# Developing future resilience from signatures of adaptation across the sorghum pangenome

**DOI:** 10.1101/2025.08.01.667986

**Authors:** Geoffrey P. Morris, Avril M. Harder, Adam L. Healey, Chloee M. McLaughlin, Brian R. Rice, Clara Cruet-Burgos, Jerry W. Jenkins, Joanna L. Rifkin, Shengqiang Shu, John J. Spiekerman, Carl J. VanGessel, Erica Agnew, Alain Audebert, Kerrie Barry, Ivan Baxter, Gregory Beurier, Lori Beth Boston, Richard E. Boyles, Siobhan M. Brady, Victoria Bunting, Jacqueline M. Chaparro, Chaney Courtney, Joseph Sékou B. Dembele, Santosh Deshpande, Cyril Diatta, Nathaniel Eck, Andrea L. Eveland, Jacques M. Faye, Daniel Fonceka, Boubacar Gano, Marie de Gracia Coquerel, David Goodstein, Jane Grimwood, Matthew E. Hudson, Jana Kholova, Katherine Johnson, Kristen K. Johnson, Dorota Kawa, Mamoutou Kouressy, Stephen Kresovich, Scott Lee, Peggy Lemaux, Robert Lowery, Delphine Luquet, Fanna Maina, Todd P. Michael, Taye T. Mindaye, John Mullet, Philip Ozersky, Christopher Plott, Jessica E. Prenni, Gael Pressoir, Jean-François Rami, Trevor W Rife, Jocelyn Saxton, Bassirou Sine, Avinash Sreedasyam, Jayson Talag, Niaba Teme, Mitchell R. Tuinstra, Vincent Vadez, John P. Vogel, Rachel Walstead, Jianan Wang, Jenell Webber, Melissa Williams, Todd C. Mockler, Jesse R. Lasky, Jeremy Schmutz, Nadia Shakoor, John T. Lovell

**Affiliations:** Department of Soil and Crop Sciences, Colorado State University, Fort Collins, CO, USA; Genome Sequencing Center, HudsonAlpha Institute for Biotechnology, Huntsville, AL, USA; Department of Horticulture and Landscape Architecture, Colorado State University, Fort Collins, CO, USA; Department of Agronomy and Horticulture, University of Nebraska, Lincoln, NE, USA; US Department of Energy Joint Genome Institute, Berkeley, CA, USA; Donald Danforth Plant Science Center, St. Louis, MO, USA; CIRAD, UMR AGAP, Montpellier, France; CIRAD, INRAE, AGAP, University Montpellier, Institut Agro, Montpellier, France; Department of Plant and Environmental Sciences, Clemson University, Clemson, SC, USA; Howard Hughes Medical Institute, UC Davis, Davis CA USA; Arizona Genomics Institute, University of Arizona, Tucson, AZ, USA; Faculté de pharmacie, Université des Sciences, des Techniques et des Technologies (USTTB), E 423 Bamako-Mali; International Crops Research Institute for the Semi-Arid Tropics (ICRISAT), Patancheru, Telangana, India; Institut Sénégalais de Recherches Agricoles, Thiès, Sénégal; DOE Center for Advanced Bioenergy and Bioproducts Innovation (CABBI) and Department of Crop Sciences, University of Illinois at Urbana-Champaign, Urbana, IL, USA; Center for Craniofacial Molecular Biology, Herman Ostrow School of Dentistry, University of Southern California, Los Angeles, CA, USA; Department of Plant Biology and Genome Center, University of California, Davis, Davis, CA, USA; Experimental and Computational Plant Development & Plant Stress Resilience, Institute of Environmental Biology, Department of Biology, Utrecht University, The Netherlands; Institut d’Economie Rurale du Mali,BP:258 Bamako-Mali; Department of Plant and Microbial Biology, University of California, Berkeley CA, USA; Innovative Genomics Institute, University of California, Berkeley CA, USA; Lone Wolf Biotech, St. Louis, MO, USA; Institut National de la Recherche Agronomique du Niger, BP: 429, Niamey, Niger; The Plant Molecular and Cellular Biology Laboratory, The Salk Institute for Biological Studies, La Jolla, CA, USA; Department of Cell and Developmental Biology, School of Biological Sciences, and Center for Marine Biotechnology and Biomedicine, University of California, San Diego, La Jolla, CA, USA; Department of Science and Conservation, San Diego Botanical Garden, Encinitas, CA, USA; Ethiopian Institute of Agricultural Research, Addis Ababa, Ethiopia; Texas A&M University, College Station, TX, USA; CHIBAS, Centre Haïtien d’Innovation en Biotechnologies et pour une Agriculture Soutenable, Croix des Bouquets, Haïti; Department of Agronomy, Purdue University, West Lafayette, Indiana, USA; IRD, Université de Montpellier, DIADE unit, Montpellier, France; School of Tropical Agriculture and Forestry, Hainan University, Haikou, China; Department of Biology, Pennsylvania State University, University Park, PA, USA

## Abstract

While the green revolution adapted a handful of crops to homogenous and high-input industrialized agriculture, much of the global population still relies on local food production from low-input smallholder farms that grow highly variable crop cultivars. The high diversity of the grain and bioenergy crop sorghum ^1–4^, and many other crops that were not homogenized during the green revolution ^5^, not only provides the raw materials for breeders to make substantial gains in cultivar improvement, but also constrains breeding efforts due to highly specialized locally adapted plant phenotypes ^6^. Here, we construct a 33-member pangenome and identify trait-associated variants in 1,988 cultivars and landraces. We then apply these resources to explore the complex interplay between historical contingency, ongoing adaptation, and the potential for future gains through climate-aware genome-enabled breeding. Specifically, our analyses conclusively demonstrate that multiple nested, deeply diverged, and previously uncharacterized structural variants in the domestication gene *SHATTERING1* distinguish the previously established multicentric origin of sorghum. We then apply landscape genomics tests to reveal how gene flow, adaptation, and secondary contact created the complex genetic mosaic in current global breeding networks. Further analysis of climate-gene associations highlights candidate loci underlying adaptation, including the biosynthetic gene cluster for the cyanogenic glucoside dhurrin. Combined, the pangenome-informed variants developed here will enable both trait discovery and subsequent marker assays to accelerate breeding and provide a framework for similar applications in other diverse and non-model crops.

## MAIN TEXT

Expanding food security and economic prosperity under rapidly changing environmental stresses will require transformative advances in global crop improvement speed and efficacy ^7–9^. New strategies for future agricultural transformations can take lessons from the past ^10–12^, including domestication, diversification, and, more recently, the Green Revolution, which developed technology packages with broadly-adapted high-yielding varieties and standardized high-input agronomic practices ^13,14^. In terms of increased agricultural productivity, the Green Revolution strategy was hugely successful for some key crops and geographies (e.g. both wheat and rice in East Asia) ^15^. However, for many indigenous African crops, including pearl millet, yams, cowpea, and sorghum, this approach has not succeeded ^16^. For example, Green Revolution-style sorghum varieties, developed for North America or South Asia, were brought directly into small-scale agriculture in Africa or incorporated into smallholder-oriented breeding programs ^17^; however, this germplasm did not conform to the demand of producers or consumers and adoption has been limited ^18^. The complexity of local adaptation in indigenous and small-holder crops to a diverse mosaic of climates and cultures ^19,20^ may underlie this stark difference in 20th century agricultural development outcomes among crops and regions.

Sorghum (*Sorghum bicolor* L. Moench) is one the most climate-resilient and phenotypically diverse major crops, adapted not only to environmental stresses, but also agronomic practices and end-uses, often in intricately interconnected fashions ^4,5^. For instance, commercial sorghum hybrids in the temperate regions of the Americas, Asia, and Australia are typically single-purpose grain, forage, or energy types ^1^; however, open-pollinated or inbred varieties grown by small-scale producers in tropical Africa, Asia, and the Americas are typically multi-purpose (grain, forage, and biomass). Further, while temperate grain sorghums are photoperiod insensitive ^6^, small-scale producers in the tropics grow photoperiod sensitive varieties adapted to narrow latitudinal bands ^21^. This diversity, while valuable to breeders as raw materials for selection ^7^, has also been a roadblock to the implementation of modern breeding approaches, which use large homogenous pools of elite breeding germplasm ^22^ to develop broadly adapted improved varieties ^18^. In response to these setbacks, localized participatory breeding approaches, which engage farmers and consumers, have been used to develop locally-improved sorghum varieties ^2,3^.

Despite its potential, breeding purely for locally adaptative trait combinations is not a panacea. Low-resourced breeding programs also need to leverage resources from the global research community and share advances (e.g. elite germplasm, traits) among national agricultural research systems (NARS) ^12^. This could be accomplished with decentralized networks of regional breeding programs, where local climate and culture are integral to breeding product profiles ^23,24^, and an *a priori* set of variants that underlie important trait variation and co-ancestry groups. Thus, a global pangenome resource — including (1) a reference genome to anchor analyses, (2) many genomes from locally adapted cultivars, and (3) whole genome resequencing of diverse germplasm — is essential to accomplish this vision.

### Anchoring sorghum genetic diversity with an improved reference genome

The ‘BTx623’ cultivar (PI 564163) has served as the reference genotype for sorghum genetics since 2009 ^25–27^, and its 2013 ‘V3’ genome assembly ^28^ remains the most important global reference resource for research and breeding. BTx623 is also well-known to the global breeding community as a parent line for commercial grain and bioenergy hybrids in the United States ^29^, the first commercial hybrids in Sudan and Niger in Central and West Africa ^24^, and as the genetic background for many trait discovery and pre-breeding resources ^30^.

Like genome assemblies that used similar technology, V3 BTx623 contained many large gaps with unknown sequences, which have been shown to bias variant detection and candidate gene annotation ^31^. Recently, long-read genome sequencing technologies allowed us to complete the repetitive portions of the genome and fill other gaps (e.g. Mimulus & Cotton ^32,33^). Consequently, our updated ‘V5’ BTx623 genome assembly represents the 10 sorghum chromosomes with only 34 contigs (contig N_50_ = 50.74Mb) — a 140-fold improvement over the 4,783 contigs used to assemble V3 (Supplementary Fig. 1A). Similar to many recent upgrades of older genome sequences ^34^, repetitive regions in V5 are far more extensive (e.g. 2.86x more contiguous centromere ^35^ blocks) while the gene-rich portions are only moderately improved (BUSCO: V3 = 98.3%, V5 = 99.7%). Importantly, V5 corrects several structural scaffolding errors in V3 (Supplementary Fig. 1B). While many of these large mis-oriented regions of V3 are highly repetitive, V5 clarifies the positions of several key genes that underlie local adaptation among breeding gene pools, including the flowering time locus *Maturity1* ^26^, the plant height genes *Dwarf2* and *Dwarf3* ^36,37^, and the dhurrin biosynthesis gene *POR* ^*38*^ (Supplementary Data 1-2). Linkage mapping and candidate gene inference in these regions will be far more accurate with a reference that is collinear with the recombination landscape.

### A 33-reference pangenome and 1,988 member diversity panel for trait discovery

As an admixed breeding line, BTx623 serves as a solid reference to anchor genome-wide genetic variation, including across our diversity panel of 1,988 unique genotypes re-sequenced with high coverage whole-genome short-read libraries. This diversity panel also spans tremendous phenotypic variation including the morphological traits associated with botanical types (*n* = 818 total libraries with botanical assignments), growth rate, and stress responses (Fig. 1), and represents many existing (e.g. biomass association panel *n* = 375) and new (e.g. terra *n* = 220) populations of interest, a host of germplasm groupings (*n* breeding = 500, *n* landrace = 737) and 746 genotypes with georeferenced GPS coordinates (Supplementary Data 3).

**Figure 1.**
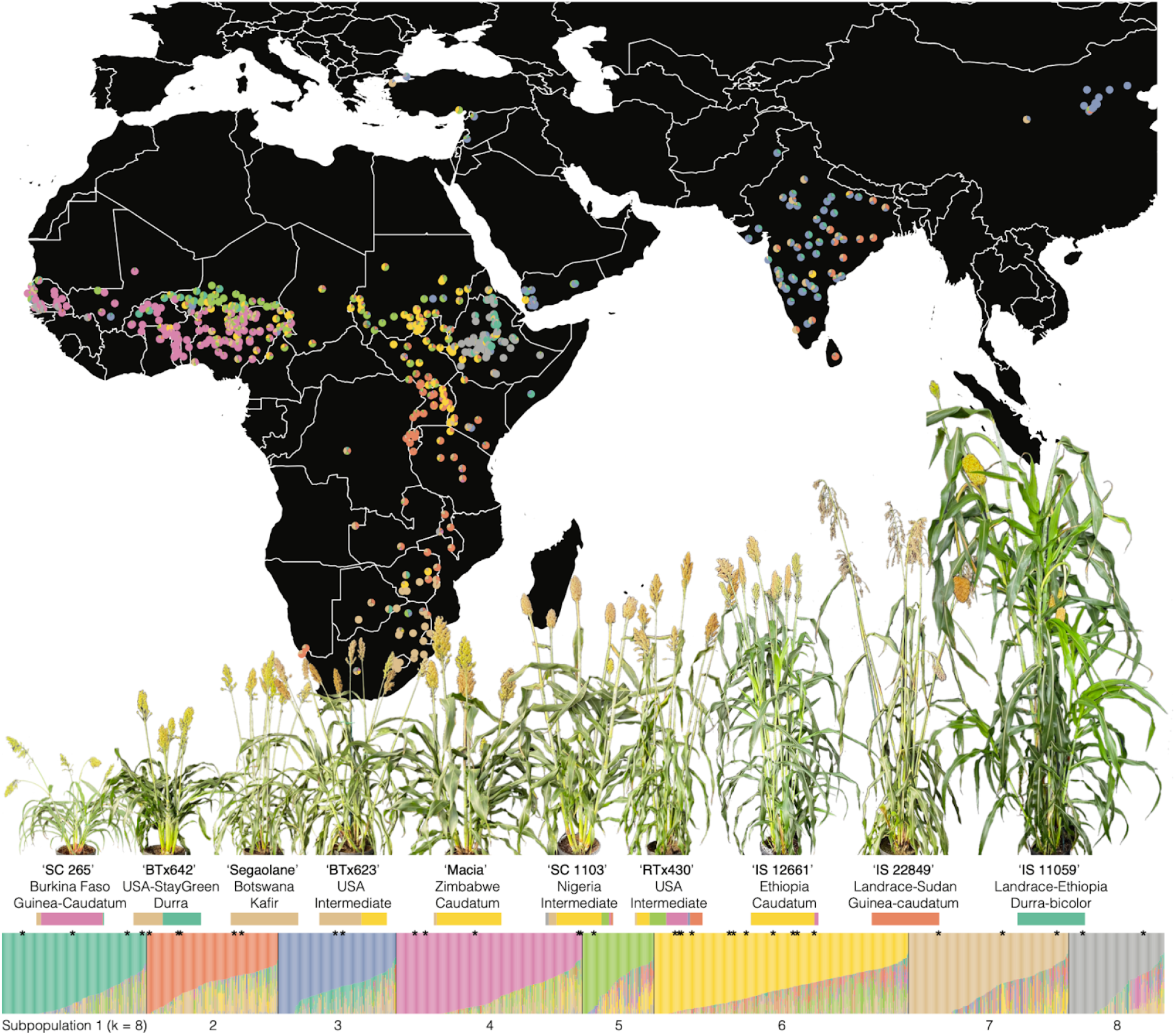
Geographic distribution of sorghum diversity panel. The whole-genome resequencing panel of 1988 unique genotypes were clustered into eight subpopulations. Ancestry proportions to subpopulation are shown for all libraries (vertical bars, barplot-bottom) and for the 693 unique African and Asian cultivars with georeferenced collection locations (pies, map-top). The 33 genotypes with reference-grade assemblies in our pan-genome resource are flagged (*) above the barplot. Ten pangenome members that span the genetic and phenotypic diversity of our pangenome are labeled (including the country of origin, and botanical type), with representative photographs (above label) and their ancestry proportions (below label).

While the BTx623 genome forms the foundation for single-reference genomic analyses, it is clear that the substantial diversity across cultivated sorghum necessitates a more comprehensive approach to genomics-enabled breeding. The recent development of multiple separately assembled genomes within many species, including sorghum ^39,40^, has revealed functionally important large-scale sequence variation ^41^, which is often not perfectly captured by short reads mapped to a single reference genome ^42^. Multiple reference genomes, when integrated into a single queryable resource (i.e., a ‘pangenome’), can be used to more accurately detect and annotate putative functional single nucleotide (‘SNP’) or small insertion-deletion (‘INDEL’) variants and are often necessary to confirm large-scale structural (‘SV’) and presence absence variation (‘PAV’). This is especially true in phenotypically diverse crops like sorghum with multiple deeply diverged gene pools ^43^ (see below). To facilitate these inquiries, we generated a 33-member pangenome resource, spanning four genetic model genotypes: ‘BTx623’ V5, readily-transformable ‘RTx430’ ^44^, stay-green & drought tolerant ‘BTx642’ ^44^, sweet ‘Wray’; nine genotypes spanning all genetic subpopulations and botanical types across our sorghum genetic diversity panel (Fig. 1, Supplementary Fig. 2); and 20 of the most important cultivars for the global sorghum improvement community (e.g. global breeding: ‘Macia’ / ‘IRAT204’, local adaptation: ‘CSM-63’ / ‘Mota Maradi’, stay-green: ‘SC 35’, aluminum tolerant: ‘SC 283’, Striga resistant: ‘SRN39’). See Supplementary Table 1 and Supplementary Note 1 for full descriptions of the pangenome members.

We characterized molecular and trait variation across the pangenome using a comprehensive suite of phenotyping approaches including field-based measurements of key agronomic traits (height, flowering time, panicle architecture, and yield components), high-throughput image-based morphology and growth traits in the greenhouse, and physiological assays including ionomic profiles, root traits, and Li-Cor-based gas exchange measurements (Supplementary Data 4). Combined, we noted very diverse phenotypes under different water availability scenarios. For example, lines RTx430 and CSM-63 (an elite African breeding line) consistently ranked highly in water-use efficiency (WUE), plant area (cm^2^), and height (cm) under both well-watered and water-limited conditions, highlighting their potential as broadly adaptable breeding candidates. Additionally, SRN39, another elite African line from Sudan, stood out for its strong performance specifically under water-limited conditions, suggesting valuable drought resilience traits.

Combined the 33-member pangenome nearly doubles the total sorghum sequence relative to the BTx623 linear reference (Fig. 2B) and includes 325 large SVs (179.6 Mb >100 kb) and 26 very large >1 Mb inversions found in multiple assemblies (Fig. 2A, Supplementary Data 5). Outside of these inversions, our sorghum genomes are remarkably collinear and lack interchromosomal translocations (e.g. ^45^), centromeric structural variants (e.g. ^32^), or other factors that are common in plants and complicate pan-genomic integration and breeding.

**Figure 2.**
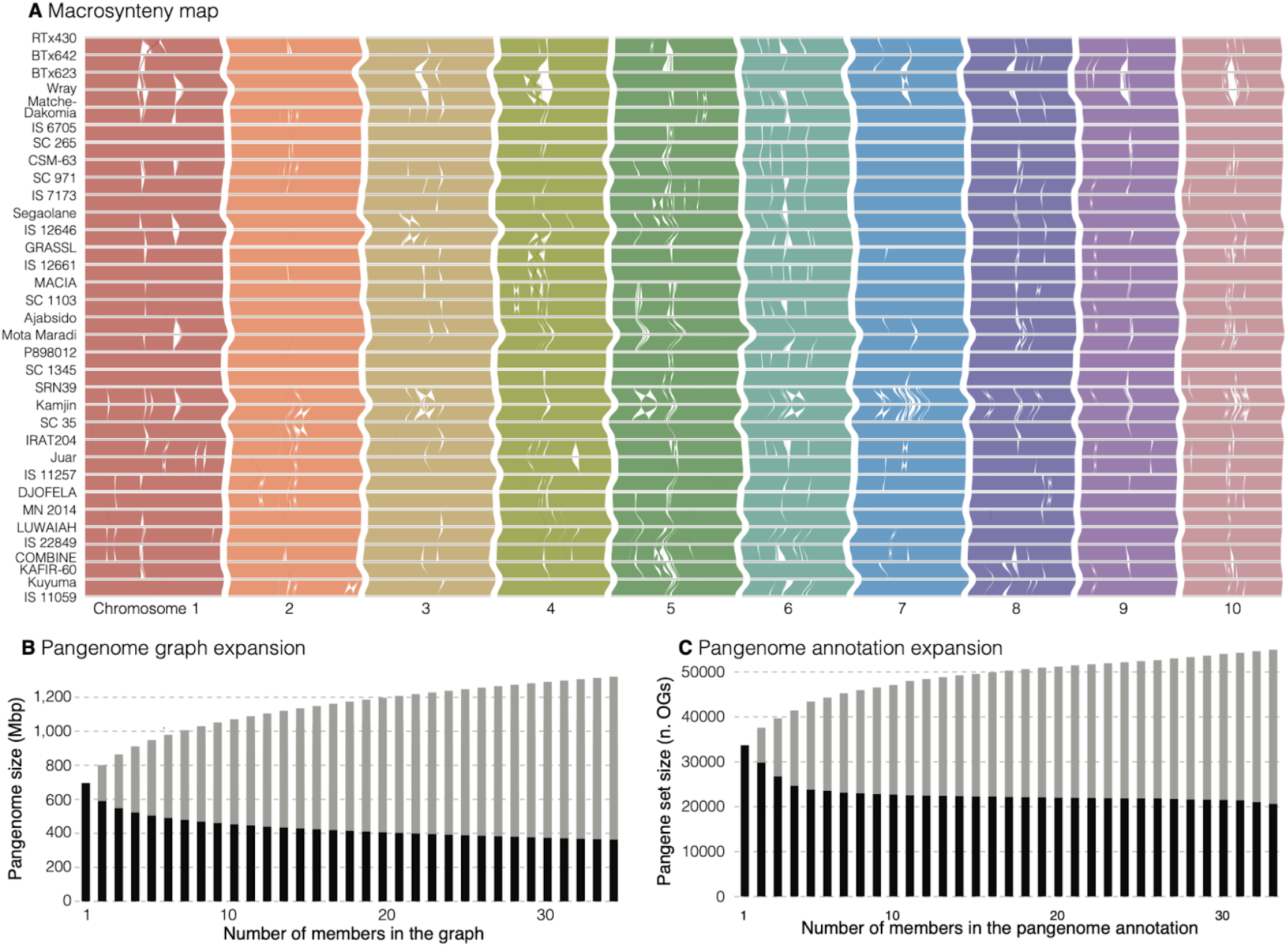
Pangenome synteny map and PAV. **A** Syntenic blocks were calculated from windowed alignments and colored by the query chromosome. To aid with visualization, all genomes were scaled to the same x-axis extent; genome sizes can be found in Supplementary Table 1. **B** Panacus and **C** OrthoFinder (phylogenetically hierarchical orthogroup) pangenome expansion curves. In both plots, black bars are the total sequence/genes found across the entire pangenome with that many members, and grey bars are the remainder that segregates across the pangenome.

The pangenome also provides an opportunity to better define structural and presence-absence variation (PAV) of protein-coding genes. However, methodological considerations can be critical. For example, pangenomes built without independent RNA and flcDNA support for each genome can under-estimate variable copy number (CNV) genes due to their reliance on sequence similarity from related species. Conversely, differing methods, sequencing support, or statistical stochasticity can inflate estimates of gene PAV through an abundance of spurious false-positive gene models ^46^. Here, we sought to leverage the strengths of both approaches by first annotating all 33 members of the pangenome with deeply sequenced short and full-length RNA reads (Supplementary Data 6), then purging rare gene models without strong support across any lines of evidence. These efforts produced a ‘pangene set’ with 988,756 gene models across 54,959 phylogenetically hierarchical orthogroups (‘gene families’; Supplementary Data 7). The majority of genes fall into the ‘core’ (100% presence; 691,667 / 70.0% of genes) or ‘soft-core’ (> 90% absence; 101,316 / 10.2% of genes) PAV categories (Fig. 2C); however, some of the most important genes for domestication and ongoing crop improvement display PAV or CNV across the pangenome. For example, several genes that are associated with resistance to the hemiparasitic plant witchweed show PAV in our pangene sets (e.g. LGS-1 sulfotransferase Sobic.005G213600). Combined, variable gene content and the abundance of large-scale structural variation indicate that our representation of the sorghum pangenome may facilitate discovery and implementation of otherwise unknown but large-effect alleles.

### Integrating single-reference and pangenome approaches to better understand multiple domestication events

Pangenome representations of multiple genomes can clearly enable trait discovery among the reference genotypes. However, the next step of identifying and reconstructing alleles across resequenced populations is both conceptually challenging and computationally intensive. The current state of the art methods require reconstruction of a pangenome graph and subsequent re-alignment of all short reads back to the updated graph whenever new references are included. This is impractical now that new genomes are produced regularly. Despite its impracticality, graph-based genotyping does capture structural variation and reduce false variant calls (e.g. pseudo-heterozygosity; Supplementary Fig. 3) and should be considered once a stable set of reference genomes have been decided on by the community. As such, a fast and easily updatable pangenome informed genotyping approach is obviously needed. This problem can be tackled with allele-to-sequence dictionaries for genomic regions of interest ^47,48^ where exact matches of diagnostic short sequences (here, 80bp ‘kmers’) are counted for each putatively causal allele. This kmer-based genotyping approach of *a priori* known alleles is agnostic to the genomic complexities that can cause misalignment and false discovery of single-reference-based variants and readily recapitulates fragment length polymorphism detected with PCR (Fig. 3A).

**Figure 3.**
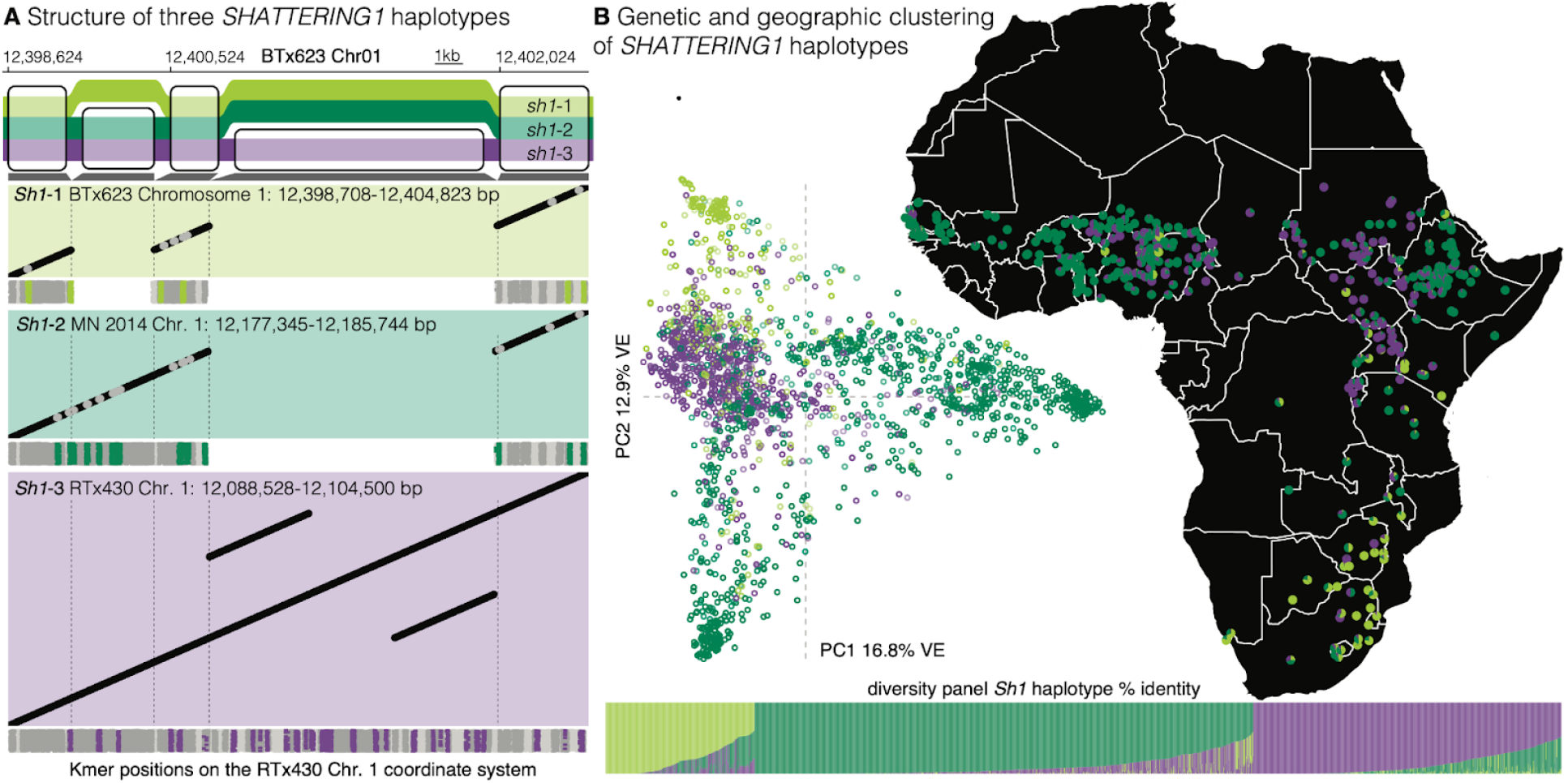
Deep divergence between domestication locus haplotypes. **A** Simplified sequence tube map showing only large INDEL variants (≥1kb) between genomes with the three *Sh1* haplotypes. Representatives of each of the haplotypes were aligned to the longest *sh1*-3 haplotype and identical (black) or variable (grey) 100bp alignments are shown in the dotplots, revealing an insertion relative to *sh1*-*1* and an exact segmental duplicate in *sh1*-3. The tubemap coordinates (BTx623) are loosely translated to RTx430 coordinate system following the grey blocks below the tubemap and the major structural mutation bounds are shown with dotted vertical lines. Positions of uninformative (dark grey) non-unique (light grey) and diagnostic (colors follow tubemap) kmers are shown below each dotplot. **B** The most closely matching haplotype is shown for resequencing libraries in genome-wide principal component space (see Supplementary Fig. 2 for details). Ancestry proportions for georeferenced African members of the diversity panel (top: pies on the map) and all libraries (bottom: bar chart) illustrate previously observed strong spatial and genome-wide population genetic structure relative to *Sh1*.

To illustrate the power of this approach, we explored *SHATTERING1* (*SH1*), a YABBY transcription factor that causes loss of seed shattering and is associated with broad geographic divergence and possibly multiple independent domestication events in sorghum ^49–51^. Initial exploration of this locus in our pangenome graph revealed known variants that distinguish the three common haplotypes: *sh1*-1 (aka *Sh1*^BTx623-like^, *Sobic*.*001G152901*) is typical of BTx623 and represents the shortest sequence due to a known 2259bp deletion that disrupts exon 2-3 relative to the undomesticated sequence; *sh1*-2 (aka *Sh1*^SC 265-like^, *SbPI156178*.*01G139400*) is the most common haplotype and harbors a putative splice modifier; and *sh1*-3 (aka *Sh1*^RTx430-like^, *SbiRTX430*.*01G157400*) exhibits many putatively functional^50^ non-synonymous SNPs (Fig. 3A). Our exploration of these haplotypes also revealed a previously undocumented 7,856bp insertion found only in *Sh1*-3 that includes a perfectly identical 2,161bp segmental duplicate (Fig. 3A). Despite substantial effort to clone and explore *SHATTERING1*, this major structural variant and untypable (by single-reference short read alignment) exact duplicate were unknown prior to our *de novo* assembly of RTx430 and our other pangenome members.

Diagnostic kmers that distinguished these three haplotypes were distributed across the entire interval, both at the insertion/deletion breakpoints and across linked shorter variants (Fig. 3A), allowing us to classify all of the short-read resequencing panel members by identity to the three major haplotypes (Fig. 3B). Previously, the three non-shattering alleles were shown to be most similar to functional *Sh1* alleles in wild sorghum populations in Tanzania, Nigeria, and Kenya, respectively ^51^. Here, we find that the landrace distribution of shattering alleles is partially congruent: *sh1-*1 in SE Africa, *sh1-*2 in West Africa (but also Ethiopia) and *sh1-*3 in East Africa (Fig. 3B), likely influenced by partially independent domestication events in different regions ^51^. However, the distribution of *Sh1* alleles is partially incongruent with genome-wide population structure (Supplementary Fig. 2). In particular, *sh1-*2 was shared by the most diverged groups of landraces: *guinea* vs *durra* types (*F*_*ST*_ = 0.073). Introgression of shattering may have contributed to domestication of two isolated wild gene pools and thus the wide dispersion of durra and introgression of local wild alleles into *guinea* types ^40,52^. Combined, these results illustrate that single-reference-based trait associations and subsequent pangenome-enabled exploration of causal alleles provides a powerful framework for trait and diagnostic marker discovery, even in complex and un-alignable regions like the *sh1*-3 exact segmental duplication.

### Exploring the balance between climate adaptation and migration in sorghum germplasm

Despite ancient divergence between *Sh1* haplotypes and evidence of multiple independent domestication events ^50,51^, the global sorghum gene pool is only moderately differentiated (k_8_ subpopulation *F*_*ST*_ = 0.170), an observation that likely results in part from abundant contemporary and ancient germplasm exchange (i.e., human-mediated gene flow). Following domestication, broadly beneficial alleles (e.g. loss of shattering) introgressed at a continental scale while local environmental and breeder selection pressures imbued cultivars with distinct trait combinations. This combination of ancient domestication alleles (Fig. 3), local adaptation, reproductive barriers, high-diversity cultivation methods ^4,53,54^, and gene flow has created distinct, partially reproductively isolated ‘botanical types’ ^39,55^ that reflect selection from local conditions as well as consumer and breeder preferences ^4,5^. To enable the efficient translation of this diversity into accelerated crop improvement, contemporary decentralized global breeding networks require a broad-based understanding of the patterns and processes shaping local adaptation and migration across the landscape.

We tested for landscape genetic signatures of migration and gene flow by analyzing spatial patterns of genetic connectivity (sharing of alleles) among 433 georeferenced African & southern Arabian cultivars (see methods) through two related approaches: (1) ‘multilocus wavelet genetic dissimilarity’ ^56^, which models allelic turnover and identifies spatial scales where genetic similarity is higher or lower than expected under random mating; and (2) ‘estimated effective migration surfaces’ ^57,58^, a graph-based method that describes spatial patterns and local rates of gene flow across the landscape at a fixed spatial scale. Both approaches clearly demonstrated the impact of human-mediated gene flow on sorghum diversity. For example, although both wild *Arabidopsis thaliana* and cultivated sorghum are selfing species with limited natural seed dispersal, Arabidopsis populations exhibited higher allelic turnover than expected under random mating at geographic scales above 7km ^56^, >17x lower than the 125 km in sorghum (Fig. 4A). Although this result indicates generally higher connectivity among sorghum populations, effective migration surfaces clearly demonstrate areas of low gene flow, especially in the Ethiopian highlands (Fig. 4B) and in the western and central Sahel (present day Senegal and Mali-Niger border, Fig. 4B-C). Consistent with locally low rates of effective migration, both the Ethiopian highlands and Sahal appear to have experienced secondary contact among domestication haplotypes (Fig. 3B) and are locations with strong evidence of local adaptation across steep environmental gradients in sorghum and other crops ^19,20,49–51^.

**Fig. 4.**
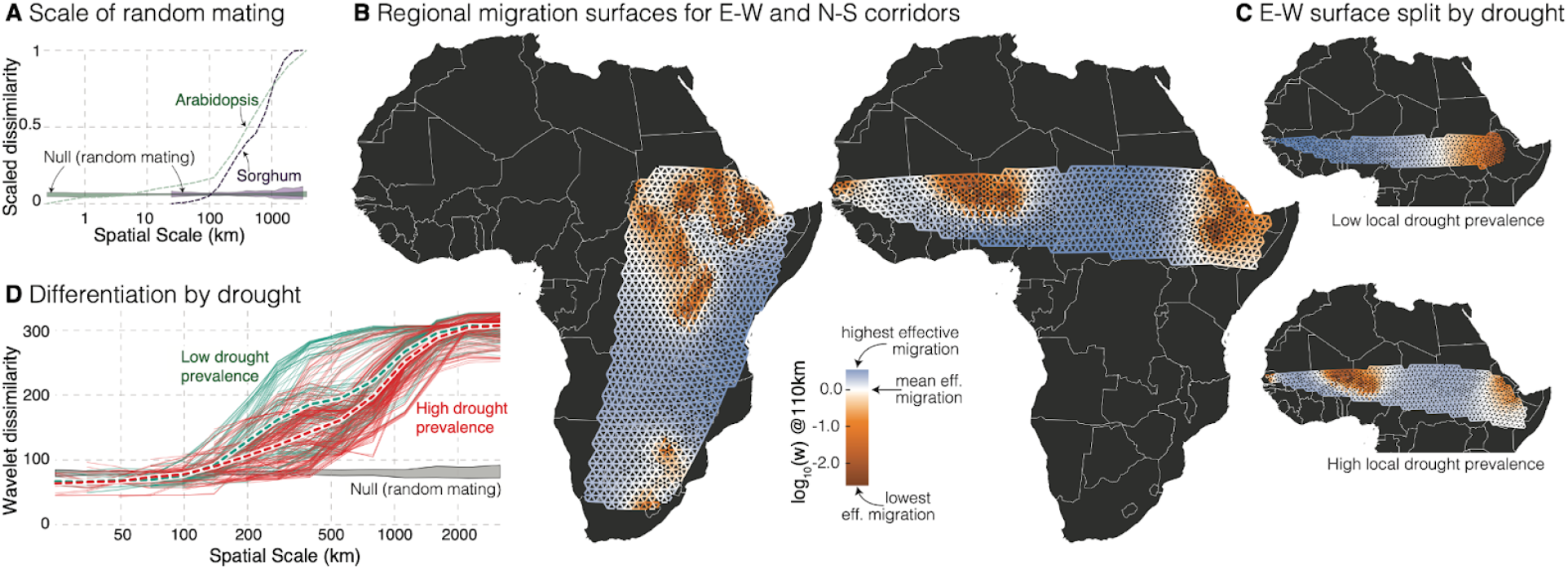
Spatial and climatic context of gene flow. **A** Comparison of the geographic scale of allelic turnover (multilocus wavelet genetic dissimilarity) in Arabidopsis (green) and sorghum (purple). Dashed lines are mean observed differentiations, and colored polygons indicate null (random mating) range based on permutation tests. **B** Estimated effective migration surfaces among sorghum populations in north-south (left) and east-west (right) corridors, which have each been shown to harbor adaptive variation in cultivated sorghum. High gene flow (blue) and low gene flow (red) are indicated by colored edges in the ‘wire-mesh’ graph. Graph grid resolution corresponds to cell spacing (distance between mid-points of adjacent cells) of 110 km. **C** Estimated effective migration surfaces between low- and high-drought prevalence sorghum populations in the east-west corridor. **D** Comparison of the geographic scale of allelic turnover between low drought prevalence (teal) and high drought prevalence (red) sorghum populations; means of each grouping are shown in white-bordered dashed lines.

These spatial analyses of allele sharing also implicate drought as a major force in historical and contemporary sorghum adaptation^59^. To test the hypothesis that drought adaptation impacts the geographic scale of gene flow, we binned accessions by those originating from high (*n* = 258) and low (*n* = 175) drought prevalence populations. Across Africa and at large spatial scales (∼150-1000 km), we found clear evidence that accessions from high drought prevalence populations had lower allelic turnover than those from less drought prone populations (Fig. 4D), suggesting alleles are more likely to move between spatially disjunct populations experiencing drought. Importantly, this pattern was robust across the continent, with similar reductions in drought-adapted allelic turnover in each of six non-overlapping ‘geographic regions’ (Supplementary Fig. 4). However, migration patterns in the Sahel complicated this finding. Here, our previous observation of generally lower effective migration rates (Fig. 4B) persisted only among cultivars from Northern, more drought prone habitats. Conversely, effective migration among cultivars from less drought-prone Southern habitats was higher than average (Fig. 4C).

The hypothesis that drought-adapted cultivars share putatively beneficial alleles across broad geographic areas can also be tested through genome-wide selection scans. Here, we applied two independent but complementary approaches: (1) *a priori* tests for different ‘extended’ larger-than expected haplotypes^60^ between high and low-drought prevalence in each of the six geographic regions; and (2) drought-agnostic scans for wavelet outliers that show higher allelic turnover than the background. Within each geographic region, the total length of extended haplotypes was not significantly different between high and low-drought prevalence locations (dry = 61.8Mb, wet = 57.4Mb; paired *t* = -0.1,*P* > 0.1). However, extended haplotype overlap between any pair of geographic regions was much greater in high-than low-drought prevalence accessions (total pairwise overlap: dry = 12.2Mb, wet = 8.4Mb; paired *t* = 3.8, *P* = 0.0013). This finding independently corroborates our previous observations of reduced allelic turnover and increased effective migration rates among cultivars in high drought regions at large spatial scales. Our drought-agnostic wavelet outlier scan also supported these findings: despite having no climatic information, wavelet outliers at large geographic scales (800-1200km) were concentrated in and near drought-associated genes (Supplementary Data 8, Supplementary Fig. 5). Individual genes included in these signatures have been previously shown to be involved in osmotic stress response ^61^, salt tolerance and dhurrin biosynthesis ^62^, and lignin deposition, particularly in response to drought stress ^63,64^, providing high-value targets for genome-enabled breeding methods.

**Fig. 5.**
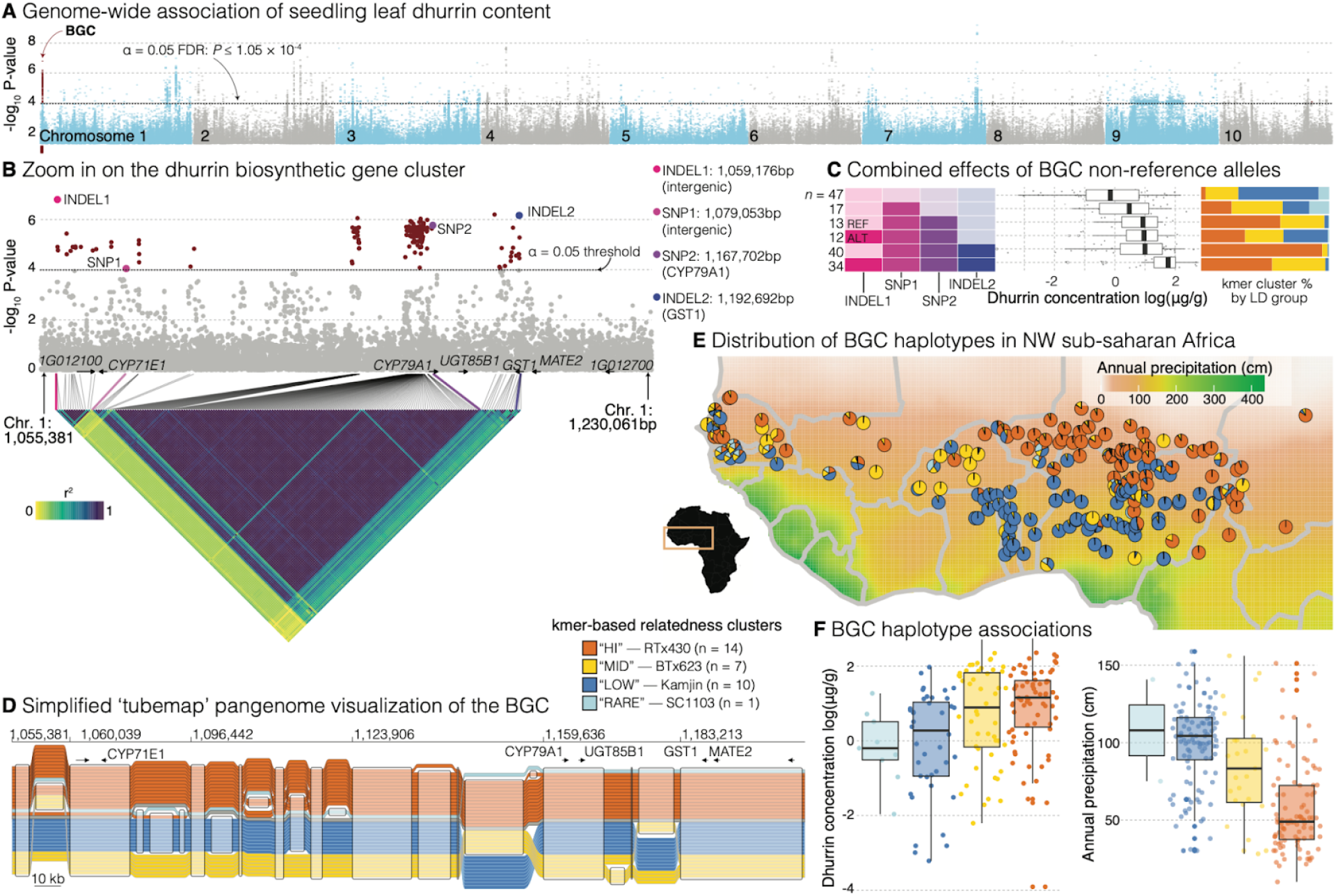
Phenotypic associations and structural variation in the dhurrin biosynthetic gene cluster. **A** GWAS ‘Manhattan’ plot across the BTx623 V5 genome coordinate system. **B** Zoom-in Manhattan plot, highlighting the five genes of known function and the two putatively non-functional bounding genes of the BGC with a linkage disequilibrium heatmap showing marker correlations for all significant variants in the interval. Four variants that are representative of the four major LD blocks in the interval are labeled. **C** Six common combinations of homozygous genotypes (BTx623 homozygote is lighter, alternative is darker) across the four major blocks in the BGC and the phenotypes of plants with those genotype combinations. The proportion of exact matches across the four kmer clusters for each of the genotype combinations are presented as colored bars to the right. See Supplementary Fig. 8 for two additional partially linked blocks and contributions of missing and heterozygous genotypes. **D** The 33 pangenome references were clustered into four BGC groups, and a ‘recombinant’ grey unclustered group for ‘IRAT204’, based on kmer similarity. The tubemap shows SVs ≥5kb with sequences shared across specific haplotypes (i.e, nodes in the pangenome graph) indicated by transparent rectangles. **E** The distribution of identity % for the four BGC pangenome groups across all northwestern sub-saharan Africa members of the diversity panel; colors follow panel D. **F** the phenotypic and climate distribution of the four clusters; annual precipitation is shown just for the region highlighted in panel E while dhurrin content is from all phenotyped members of the diversity panel.

### Future targets for molecular breeding: exploration of sequence diversity in the dhurrin biosynthesis pathway

While continent-scale gradients of abiotic stress intensity have clearly shaped sorghum diversity, local agricultural productivity is often equally affected by adaptation to complex interactions between pests, pathogens, and nutrient availability. Drought and pest adaptive responses (e.g. stomatal regulation, structural leaf defense, etc.) are obvious individual targets for breeders and natural selection alike, and some secondary metabolites offer pleiotropic resistance to both stresses. For example, the cyanogenic glycoside dhurrin not only provides resistance to chewing insect herbivory via cyanide release ^65^, but also improves constitutive dehydration avoidance ^66–68^. While part of the dhurrin biosynthesis and transport pathway is governed by a 5-gene biosynthetic gene cluster (‘BGC’) on chromosome 1 ^69,70^, the degree to which genetic variation at the BGC and other loci drive cultivated sorghum local adaptation and resistance traits is largely unexplored.

We first sought to define individual loci associated with the biosynthesis and catabolism of dhurrin by phenotyping members of the diversity panel for seedling dhurrin content (*n* = 175) and hydrogen cyanide potential (“HCNp”, *n* = 367). Contrary to expectations of a single function of dhurrin for pest defense through the production of HCNp, the two traits were only weakly genetically correlated later in vegetative growth (7-8 leaf stage; *r* = 0.16, *P* < 0.05) and uncorrelated at the seedling stage (3-4 leaf stage; *r* = 0.075, *P* = 0.33). Accordingly, none of the strongest genome-wide association (GWAS) peaks for dhurrin (Fig. 5A) and HCNp (Supplementary Data 9-11, Supplementary Figure 6) were physically proximate, potentiating a functional role of dhurrin level modification outside of pest defense potentially through known drought adaptive pathways.

Across the 1472 SNPs and short INDELs within the ∼175kb BGC interval on chromosome 1, 191 sites had significant associations with dhurrin levels (FDR < 0.05, Fig. 5B). While the majority of significant variants resided within intergenic regions, an alanine -> valine missense mutation at 1,167,702bp in CYP79A1 (*Sobic*.*001G012300*) is predicted to increase this cytochrome P450’s binding affinity for tyrosine, the substrate for the first committed step of dhurrin biosynthesis ^71^ (Supplementary Figure 7). There were also several interesting regulatory candidate variants including a 2bp CTG -> G INDEL at 1,185,567bp that disrupts predicted binding sites within an accessible chromatin region for abscisic acid-responsive transcription factors (ABF3/4), which have known functional drought-mediated effects and may affect expression of both adjacent loci (*UGT85B1*/*Sobic*.*001G012400, GST1*/*Sobic*.*001G012500*, Supplementary Table 3). The strong significance of these *cis-*regulatory motif variants meshes with a previous observation that variation in regulatory sequences may play a significant role in the evolution of dhurrin biosynthesis ^72^.

Since functional sequence variation within biosynthetic gene clusters can affect traits both directly and epistatically through modification of the pathway (e.g. abundance of precursors), it is important to consider the BGC as a whole when designing genome-enabled breeding programs to improve drought and pest adaptation through dhurrin production. Consistent with the coinheritance of and likely correlated selection on the BGC, we observed strong linkage disequilibrium (LD) blocks where all but one significant variant fell into three tightly linked clusters (Fig. 5B). These four groups of markers represent most of the unlinked variation in the region that is associated with dhurrin concentration. Combined, the total number of non-reference alleles across these four blocks (Fig. 5C) and two additional sites with higher missing data (Supplementary Figure 8) was highly predictive of dhurrin concentration due to a marginally significant additive (*t* =-2.18, *P* = 0.03) and highly significant quadratic (*t* = 4.97; *P* < 0.0001) effect on un-transformed dhurrin levels. This observation likely indicates that epistatic interactions or other non-linear effects between adjacent loci result in genotypic combinations with highly elevated dhurrin levels.

Although excluded from the association study due to high rates of missing data, several sites within the BGC exhibited an excess of heterozygotes and missing genotype calls (Supplementary Figure 8). In other systems, miss-called structural variants can be erroneously represented in single-reference short read genotyping variants as loci with high levels of missing data and excess (pseudo) heterozygosity ^73,74^, challenges that can be overcome with pangenome-enabled methods. Therefore, we applied our pangenome-kmer genotyping approach to the BGC. Haplotype sequence similarity clustering revealed that 32 of the 33 reference pangenome members fell neatly into four tight kmer-identity clusters that distinguished samples by previously typed short variants (Fig. 5C) and several previously unknown large intergenic INDELs (Fig. 5D). The unclustered member ‘IRAT204’ was intermediate and exhibited signatures of recombination between the ‘high’ and ‘mid’ clusters. The four-level clustering explained 3% more variance in seedling dhurrin content, a 15% better fit than the most predictive SNP, in part by capturing large structural differences that were not necessarily linked to SNP and small INDELs.

The kmer-based genotype clustering aggregates all variants in the interval, an approach that generally mirrors the effects of expected linked selection and combined effects of unlinked markers across BGCs both across evolutionary timespans and in ongoing breeding programs. We connected our pangenome genotyping to breeding objectives by mapping kmer clusters to phenotypic and landscape-level variation. The patterns were obvious at a continental scale: haplotypes associated with higher dhurrin content were sourced from georeferenced localities with lower annual precipitation (Kruskal-Wallis rank sum test annual precipitation by kmer de novo cluster H = 41.01, *P* < 0.0001; Fig. 5F). Critically, the evidence for this correspondence was even stronger at more local scales. For example, when we limit analysis to samples originating from West Africa, the relationship between samples with elevated dhurrin content originating from locations with lower annual precipitation is far stronger (Kruskal-Wallis rank sum test annual precipitation by kmer de novo cluster H = 77.48 *P* < 0.0001; Fig. 5F). Thus, the BGC appears to be a metabolic hub that underlies variation in pest resistance and likely is structured by adaptation to moisture availability across continental and local geographic scales.

## Conclusions

The high diversity of sorghum and many other crops that were not homogenized during the green revolution is often a product of historical and ongoing selection that has adapted cultivars to local and often stress-prone diverse agroecologies. While this diversity is a critical resource for ongoing crop improvement, variable local adaptation and consumer preference can also constrain efforts to translate breeding gains across regions: introducing broadly beneficial alleles into local breeding programs may alter grain characteristics or otherwise reduce the local value of a cultivar. Therefore, it is critical that breeders of sorghum and other locally adapted crops can quickly and effectively translate information across programs, an effort that is massively accelerated by knowledge of genes or markers that underlie trait variation and the integration of genome-enabled breeding with local consumer preferences and breeder knowledge ^19,20^. Here, we described our efforts to integrate pangenome enabled variant discovery to define targets of historical environmental adaptation, which will accelerate breeding decisions in sorghum. In addition to better characterizing large structural variants, pangenomes facilitate information transfer between genotypes used in breeding and those that are more amenable to laboratory experimentation. This is especially important in sorghum, which is notoriously recalcitrant to genome editing methods: to date none of the major breeding lines can be efficiently edited and only two genotypes (‘RTx430’ and ‘P898012’) are considered readily transformable without morphogenic regulator technology. Our pangenome bridges this gap by effectively reconstructing the putative functional alleles in breeding pedigrees and the orthologous sequence to target in transformable varieties, which will pave the way for accelerated pangenome-enabled traditional breeding and genome editing of locally adapted alleles across global sorghum germplasm.

## METHODS

### Plant material preparation and nucleic acid extractions

Young seedlings were grown in flat pans under healthy, pest- and disease-free conditions until the first fully developed leaves emerged. To optimize DNA yield and quality, seedlings were dark-treated for 24–30 hours under moist conditions. Tissue was harvested by hand in small batches, cutting ½ inch above the soil surface and immediately flash-freezing in liquid nitrogen within one minute of excision. Approximately 50g of tissue was collected in this manner, stored in pre-labeled freezer-quality ziplock bags at –80 °C, and kept frozen until extraction. DNA was extracted from young tissue using the protocol of Doyle and Doyle ^75^ with minor modifications. Flash-frozen young leaves were ground to a fine powder in a frozen mortar with liquid nitrogen followed by very gentle extraction in 2% CTAB buffer (that included proteinase K, PVP-40 and beta-mercaptoethanol) for 30min to 1h at 50 °C. After centrifugation, the supernatant was gently extracted twice with 24:1 Chloroform : Isoamyl alcohol. The aqueous phase was transferred to a new tube and 1/10th volume 3 M Sodium acetate was added, gently mixed, and DNA precipitated with iso-propanol. DNA precipitate was collected by centrifugation, washed with 70% ethanol, air dried for 5-10 minutes and dissolved thoroughly in an elution buffer at room temperature followed by RNAse treatment. DNA purity was measured with Nanodrop, DNA concentration measured with Qubit HS kit (Invitrogen) and DNA size was validated by Femto Pulse System (Agilent).

### Genome Assembly

The majority of the pangenome assemblies and BTx623 were generated by assembling the PACBIO CCS reads using the CANU ^76^ assembler, with five accessions (PI154987, PI156178, RTx430, BTx642, and PI565121) being generated using the MECAT ^77^ assembler, and one accession (Wray) being assembled using the HiFiAsm+HIC assembler ^78^. All assemblies were subsequently polished using ARROW ^79^, except BTx623 and Wray which were polished using RACON ^80^. A combination of syntenic markers from BTx642 and primary annotated genes from BTx623 were used to identify misjoins in the assembly. Syntenic markers consisted of a total of 32,400 unique, non-repetitive, non-overlapping 1.0 Kb sequences that were generated using the version 1.0 S. bicolor BTx642 genome release. The gene set consisted of a subset of 12,641 BTx642 annotated primary genes. Both the syntenic markers and the primary gene set were aligned to the polished assembly. Contigs were then ordered, oriented, and assembled into 10 chromosomes using the syntenic markers and genes. Chromosomes were numbered using the V5.0 *S. bicolor* BTx623 assembly available from Phytozome ^81^. Each chromosome join is padded with 10,000 Ns. Contigs terminating in significant telomeric sequence were identified using the (TTTAGGG)n repeat, and care was taken to make sure that they were properly oriented in the production assembly. The remaining scaffolds were screened against bacterial proteins, organelle sequences, GenBank nr and removed if found to be a contaminant. Chloroplast and mitochondrial genomes were generated using the OatK pipeline ^82^. Homozygous SNPs and INDELs were corrected in the release using the Illumina fragment reads (2×150, 400bp insert) by aligning the reads using bwa-mem ^83^ and identifying homozygous SNPs and INDELs with the GATK’s UnifiedGenotyper tool ^84^. Completeness of the euchromatic portion of the pangenome assemblies was assessed by aligning the existing *S. bicolor* BTx623 V5.1 annotated primary transcripts to the releases. The aim of the completeness analysis was to obtain a measure of completeness of the assembly, rather than a comprehensive examination of gene space. We retained genes that aligned at greater than 90% identity and 85% coverage.

### Genome Annotation

Genome annotation was performed using the pipeline developed by the DOE Joint Genome Institute and Phytozome ^81^. Each pangenome member underwent two rounds of gene prediction. In the first round gene models were independently predicted for each genome followed by the propagation of these predictions across the entire pangenome in the second round.

In the initial round, transcript assemblies were generated from 2×150 bp stranded paired-end Illumina RNA-seq reads (Supplementary Table 2) using PERTRAN ^85^ for genome-guided assembly via GSNAP ^86^, followed by splice graph construction, alignment validation, and correction. PacBio Iso-Seq CCS reads, when available, were corrected and collapsed using a GMAP-based pipeline ^86^ to refine alignments, correct splice junctions, and cluster alignments when their intron(s) matches if spliced or 95% similar if single exon. Final transcript assemblies were produced using PASA ^87^, integrating RNA-seq assemblies, corrected CCS reads, and Sanger ESTs. A species-specific repeat library was built from *S. bicolor* v3.0 using RepeatModeler ^88^. Repeats were functionally analyzed with InterProScan ^89^, incorporating Pfam ^90^ and PANTHER ^91^ databases, and those with significant hits to protein-coding domains were excluded. Genomes were soft-masked using RepeatMasker ^88^ with the curated repeat library. Gene loci were identified using transcript assembly and EXONERATE ^92^ alignments of proteins from *Arabidopsis*, soybean, poplar, *Aquilegia*, grape, rice, *Setaria viridis, Brachypodium, Panicum hallii*, pineapple, *Acorus americanus*, and Swiss-Prot (2020_06) to the repeat soft-masked genomes, with up to 2,000-bp extension on both ends unless extending into another locus on the same strand. Gene models were predicted using FGENESH+ ^93^, FGENESH_EST, EXONERATE, PASA assembly ORFs, and AUGUSTUS ^94^ trained on high-confidence PASA assembly ORFs with intron hints from RNA-seq alignments. Candidate models were selected based on EST and protein support and penalized for repeat overlaps. PASA refinement added UTRs, corrected splicing, and incorporated alternative isoforms. Cscore (BLASTP score ratio and protein coverage) was computed for PASA-refined proteins. Transcripts were retained if Cscore and coverage ≥ 0.5 or supported by ESTs; those with >20% CDS-repeat overlap required Cscore ≥ 0.9 and ≥ 70% coverage. Models with >30% TE domain overlap (Pfam) were excluded.

In the second round, each genome was hard-masked with its high-confidence gene models (supported by transcriptome and homology evidence). BLASTX and EXONERATE were used to predict new gene models by projecting high-confidence models from other pangenome members onto the hard-masked assemblies. Predicted models were retained if they showed stronger homology support than existing models and were not contradicted by transcript evidence, or if no first-round model existed at that locus. Incomplete gene models, which had low homology support without full transcriptome support, or short single exon genes (<300 bp CDS) without protein domain or good expression were manually excluded.

### Pangenome graph construction and exploration

A pangenome graph was constructed for each chromosome using Minigraph-Cactus and default settings (v2.9.3 ^95^) with the BTx623 V5 genome as the primary reference. Clipped chromosome graphs were merged and prepared for visualization using vg (v1.59.0 ^96^). We used ODGI (v0.9.0 ^97^) to inspect representation of each genome across the graph and sequenceTubeMap (v0.1.0 ^98^) to visualize variation in genomic regions of interest. For the dhurrin BGC and *Sh1* loci, we further processed this graph with vg and vcfwave (vcflib v1.0.10 ^99^) to reduce allelic complexity and retain only variants ≥5 kb and ≥1 kb in length, respectively.

### Comparative genomics

Gene families were calculated with OrthoFinder ^100^ and parsed with GENESPACE ^101^ to create saturation curves. Synteny maps were created using DEEPSPACE (github.com/jtlovell/DEEPSPACE), which aligns and tracks the positions of windowed sequence alignments and quickly visualizes large-scale structural variants. SyRI ^102^ was used for pairwise whole-genome alignment (via minimap2 ^103^) variant detection. Pangenome graph saturation curves were calculated with Panacus ^104^.

### DNA re-sequencing and variant detection

A total of 2,145 diversity samples (*n* = 1,988 unique genotypes + redundant + polishing) were resequenced at a median coverage of 43.39 (range 1.96-364.18) (Supplementary Data 3). Of these, 1,676 were used for further analysis after filtering for missing data, outlier elevated heterozygosity and collection site discrepancies. The samples were sequenced using Illumina HiSeq X10 and Illumina NovaSeq 6000 paired-end sequencing (2 × 150 bp) at HudsonAlpha Institute for Biotechnology and the Joint Genome Institute. To account for different library sizes, reads were pruned to ≤50× coverage. SNPs and short INDELs were called by aligning Illumina reads to the V5 reference with bwa mem ^83^. The resulting .bam file was filtered for duplicates using Picard (http://broadinstitute.github.io/picard) and realigned around indels using GATK 3.056 ^84^. Multi-sample SNP calling was done using SAMtools mpileup ^105^ and Varscan V2.4.089 ^106^ with a minimum coverage of eight and a minimum alternate allele count of four. Genotypes were called via a binomial test. SNPs within 25 bp of a 24-mer repeat were removed from further analyses. Only SNPs with <90% missing data and minor allele frequencies >0.001 were retained, resulting in 35,877,217 SNPs at a coverage depth between 8× and 500×. Phasing was performed using SHAPEIT3 ^107^.

### Pangenome-based genotyping

In brief, the kmer genotyping methodology extracts kmers (in this instance: 80mers that are single-copy within individual references and variable among members of the pangenome; *n*-1 references; *n* = 32). The pipeline proceeds as follows: (1) Each reference genome is individually converted into kmers, where each kmer and its frequency in the assembly is stored in a hash table. Any non-adjacent multi-copy kmers are flagged to be ignored during downstream analyses. (2) Individual reference hashes are combined in a single hash (*main_Hash*), updating the flags such that only single copy kmers among all considered references will be used downstream. To isolate kmers that are useful for genotyping for a specific region of interest (ROI; e.g. *Sh1* locus, dhurrin BGC locus), the orthologous region from each reference is extracted (from either from the pan-genome graph or best alignment bounds using minimap2 ^103^. Steps 1 and 2 above are repeated the subsetted fasta files, generating a combined subsetted hash that contains local single-copy sequences from all ROI orthologs across the pan-genome. (3) To ensure the local single copy markers do not occur elsewhere in the genome (which would prevent their use as a genotyping marker), the combined ROI hash is intersected with the main genome hash, updating flags such that any non-single copy kmers will be ignored when genotyping (*genotyping_hash*). To enable genotyping of the tandem duplicate haplotype within the *SH1* locus, the pipeline above was adjusted to allow local two-copy markers, provided both were only present within the bounds of the subsetted ROI fasta. (4) Illumina libraries were genotyped by searching for all kmers in the *genotyping_hash* and counting their frequencies. At this point, counts were either merged into a single matrix for *de novo* clustering (e.g dhurrin BGC locus), or binned based on *a priori* clusters (e.g. *SH1* locus) based on the local haplotype structure at the region of interest. *De novo* clustering as used for kmer genotyping the dhurrin BGC locus operates on a binary matrix of n × m (kmer counts × Illumina libraries), where kmer frequencies greater than one were considered present. Clusters were visualized using PCA as defined by Partitioning Around Medoids (PAM) of pairwise Jaccard distances among reference libraries. *A priori* clustering as used for kmer genotyping the *SH1* locus sorts kmers into non-overlapping bins, then counts kmer matches within each Illumina library and reports a percentage match from each bin, regardless of which reference they were derived from (e.g. LibraryA|2:97:1; for *n* matches to three haplotype bins within Library A.). Illumina library haplotypes were then assigned based on majority match to haplotype bins.

### Diversity panel source material

The sorghum diversity panel used in this study comprises 1,988 unique accessions sourced from structured diversity panels and curated regional collections. These lines span Africa, Asia, and the Americas and represent both landraces and breeding lines, providing broad coverage of sorghum’s genetic, phenotypic, and geographic variation. These include: (1) “CASP”: A set of 82 lines from a 648-line association panel, contributed by John Vogel and collaborators, previously described in Spindel et al. ^108^. This panel was developed for trait mapping under field and drought conditions. (2) “Biomass Association Panel” (BAP): A panel of 390 accessions curated by Stephen Kresovich and colleagues, representing biomass-related diversity across sorghum botanical types and global regions ^109^. The panel is widely used in bioenergy trait mapping, and 375 of the BAP accessions are included in this study. (3) “Sorghum Association Panel” (SAP): A global panel of 377 genotyped accessions developed by USDA-ARS and collaborators ^110^, representing all five major sorghum races and widely used in genome-wide association studies (GWAS). A total of 328 SAP accessions are included in this study. (3) “TERRA MEPP”: A panel developed under the DOE ARPA-E TERRA program to represent a wide range of traits relevant to field-based high-throughput phenotyping. These lines were selected to enable trait discovery under environmental variation ^111^. (4) “Purdue Inbred Calibration Set”: A set of sorghum accessions developed through Mitch Tuinstra’s breeding program at Purdue University, where emphasis lies on improving traits related to drought, cold tolerance, Striga resistance, nutritional quality, and biomass potential. These lines have played a central role in UAV-based remote sensing studies that integrate hyperspectral and LiDAR data with marker profiles, underscoring their relevance in phenomics-genomics research ^112,113^. (5) “Georeferenced Panel”: The georeferenced panel (N = 529) consisted of 166 accessions from within the BAP, 133 African accessions from the WRS, and an additional 230 accessions from NPGS-GRIN, selected to maximize spatial distribution. Georeferences were sourced from Genesys using ICRISAT IDs (“IS” numbers) and cross-referenced with NPGS-GRIN IDs (“PI” numbers) for GRIN accessions. (6) “World Reference Set (WRS)”: A global core collection of 383 sorghum accessions was established by Billot et al. ^114^ and captures broad allelic diversity within *Sorghum bicolor*, with representation from all five major cultivated races and their intermediates. The set reflects extensive geographic coverage, spanning central and eastern Africa to Western Africa, South Asia, and East Asia, with significant representation in domesticated morphotypes rather than strict race categories. Our study includes 381 of these 383 accessions, preserving nearly the full scope of genetic and geographic diversity defined in the original WRS ^114^. (7) “West African Sorghum Association Panel (WASAP)”: A 756-member diversity panel contributed by national programs in Mali, Senegal, Togo, and Niger to represent West African landraces and regionally adapted breeding lines. (8) Ethiopian Institute of Agricultural Research (EIAR): A regional collection developed and curated by EIAR to capture the exceptional diversity of Ethiopian highland sorghum. These accessions include seed-increased landraces and genotyped lines selected for local adaptation. (9) BC-NAM Parents: The panel incorporates 37 founder accessions from a West and Central African Backcross Nested Association Mapping (WCA-BCNAM) population, partially detailed by Garin et al. ^115^. These lines were selected for adaptation traits and crossed with three elite recurrent parents from the region, yielding 3,901 BC-NAM lines. The founders were chosen to maximize genetic diversity in plant height, flowering time, and environmental response across agro-ecologies varying in photoperiod, rainfall, temperature, and fertility ^115^. This panel structure reflects global efforts to unite diverse germplasm for climate-aware trait discovery and genomic breeding. Each contributor is acknowledged through authorship or citation, and the panels provide a powerful framework for both local adaptation analysis and broad-scope association mapping.

### Passport Data Collection and Curation

Passport data including botanical classification, breeding improvement status, country of origin, and geographic coordinates—were compiled from multiple sources: the GRIN-Global database, Genesis Global, the ICRISAT Sorghum Passport dataset, and previously published studies ^111,116^. Records were cross-referenced to identify and resolve discrepancies across datasets. In cases where inconsistencies could not be reconciled, both conflicting metadata entries were retained and included in the final metadata.

### Population genetics

We applied ADMIXTURE ^117^ using default settings to *n* =1988 unique genotypes of non-duplicated sequencing libraries. To generate an approximately independent set of markers, PLINK (v1.90b6.12) ^118^ we removed variants with minor allele frequency (MAF) < 0.05, more than 50% missingness, and applied LD-based pruning with a window size of 50 SNPs, step size of 5 SNPs, and an r^2^ threshold of 0.5. After filtering, we randomly selected 100,000 SNPs to estimate the optimal *K* of ancestral groups. For the remainder of population genetic analyses, we retained *n* = 433 samples georeferenced to the African continent that were designated as landraces, cultivars, or wild strains and missed a genotype call in <15% sites. To select sites that captured dynamics of population structure, we selected putatively neutral and unlinked variants using bcftools (v1.9) ^119^ with the following filtering parameters: invariant sites and sites with strand bias in variant-supporting reads (>90%) were removed. We required all retained variants to be synonymous, have a genotype call in at least 95% of individuals sampled, and a minor allele frequency (MAF) >= 0.01. Filtered genotypes were imputed with beagle ^120^ and variant effects annotated with SnpEff ^121^. We further LD-pruned the remaining SNPs in PLINK (v1.90b6.12, ^118^) using 50-SNP windows with a variance inflation factor (VIF) of 2 (VIF = “1/(1-R^2) where R^2 is the multiple correlation coefficient for a SNP being regressed on all other SNPs simultaneously,” PLINK documentation), resulting in a set of 33,823 neutral LD-pruned SNPs. We used the set of 33,823 neutral LD-pruned SNPs to estimate effective migration surfaces in “fast estimation of effective migration surfaces” (“feems”; ^57^. We ran separate analyses for the African continental axes (North - South and East - West). For each independent run of feems, we optimized the smoothing regularization parameter (λ) and utilized the optimal lambda for each analysis. Node and edge positions and weights were exported and formatted for plotting in R (R Core Team 2024). To describe patterns in allele frequency turnover at neutral sites capturing dynamics of population structure, we again used the set of 33,823 imputed, neutral, LD-pruned SNPs. Georeferenced samples (*n* = 433) were grouped into “populations” by averaging allele frequencies of samples georeferenced to the same latitude and longitude (rounded to three decimal places, ∼100 meters), resulting in 343 populations comprised of 1 and 13 individuals (mean = 1.26). We conducted “scale-specific genetic variance test” by modeling allele frequency turnover with wavelet transformations at 15 scales, spaced along a log sequence from the 0.1st to 99th percentile of distances between sites: 24km, 35km, 49km, 69km, 98km, 139km, 197km, 278km, 394km, 558km, 790km, 1119km, 1584km, 2242km, 3174km. Following methods described in ^122^, we used wavelet transformations, to determine outlier loci with more rapid spatial allele frequency turnover as compared to the overall distribution, representing loci that may be important in adaptation. We used 5,615,029 SNPs filtered for MAF > 0.01 and heterozygosity <0.9 (likely representative of genotyping errors). The top 1000 outliers from each chromosome at each spatial scale were used in downstream analyses, including a Gene Ontology (GO) enrichment using topGO (v.2.42.0), an R Bioconductor package. Using the same set of SNPs, we identified putative selective sweeps using normalized xpnsl ^60^, grouping georeferenced populations into geographic regions using Partitioning Around Medoids (PAM) and comparing individuals from high drought prevalence sites to individuals from low drought prevalence sites in each of five geographic regions (a sixth region included only high drought prevalence sites and was excluded). We retained all sites with normalized scores above the critical threshold, and converted runs of sites into blocks using the add_rle function from GENESPACE ^101^; total extended haplotypes ranged from 8-12Mb genomewide per population. We then quantified pairwise base-pair overlap of xpnsl extended haplotypes reciprocally between all geographic regions. We determined whether overlap was greater than expected by chance by comparing the observed overlap to the overlap of permuted genomic blocks, significance testing relative to 1000 permutations. We compared the amount of haplotype overlap using paired t-tests for each geographic region.

### Assignment of Landraces to Climate-Based Clusters

Following methods described in ^123^, the *Cycles* agroecosystem model was used to characterize variables representative of stage-specific water stress (vegetative, reproductive) for *n* = 326 sorghum landrace accessions’ point of origin. *Cycles* is a process-based multi-year and multi-species agroecosystem model ^124,125^ that requires a number of input files to simulate crop growth. All simulations were carried out using *Cycles* v0.13.0 (https://github.com/PSUmodeling/Cycles). The crop description file defines the physiological and management parameters that control the growth and harvest of crops used in the simulation. We used the base *Cycles sorghumMS* parameters from the default crop description file. The management (operation) file defines the daily management operations to be used in a simulated crop rotation. We activated conditional planting where *Cycles* “plants” a simulated crop once certain soil moisture and temperature levels are satisfied within a window of planting dates. We turned on the automatic nitrogen fertilization option and set planting density to 67% so that stress observed in model outputs was due entirely to climatic factors. To approximate the climate conditions the sorghum accessions were adapted to, we simulated plant growth using weather data from 1970–1989, ten years before and after the average collection year. Daily weather files at 1/4 degree resolution that overlapped with at least one of our georeferenced accessions and for years included in the simulation were fetched from the meteorological data source Global Land Data Assimilation System (GLDAS; ^126^) using the script *LDAS-forcing*.*py* sourced from https://github.com/shiyuning/LDAS-forcing. Soil physical parameters describing the average soil characteristics and land use type of each landrace accession point of origin were obtained from the ISRIC SoilGrids global database ^127^ via the HydroTerre data system ^128–130^ Using model outputs and for each landrace accession, we extracted integrative environmental variables representative of stress when in the vegetative and reproductive phase. Accessions were clustered into two groups (low and high drought prevalence) using k-means clustering implemented in the R stats package. As additional georeferenced landraces were added to the diversity set (n = 107), cluster membership was predicted using a feedforward artificial neural network (ANN) implemented in the R package neuralnet. The ANN used bioclimatic predictors from the WorldClim 2.1 dataset (1970–2000), interpolated at 10 arc-minute resolution (∼340 km^2^ per pixel). Input variables included mean temperature of the driest quarter (MTDQ), mean temperature of the warmest quarter (MTWQ), precipitation of the wettest month (PWM), precipitation seasonality (PS), precipitation of the driest quarter (PDQ), and precipitation of the warmest quarter (PWQ).

### Dhurrin phenotyping

To evaluate variation in dhurrin concentration and cyanogenic potential (HCNp) among diverse Sorghum bicolor landraces, we conducted a phenotypic analysis using accessions sourced from the USDA Germplasm Resources Information Network (GRIN). All accessions were grown under controlled conditions at the Colorado State University Plant Growth Facilities. A completely randomized block design was implemented with two replicates per accession. Four plants per accession were sown in 3-gallon pots containing Pro-Mix HP supplemented with one tablespoon of Osmocote fertilizer. Accessions were evaluated across two separate plantings, each containing two replicate blocks. Within each block, individual plants were grown and subsequently sampled for trait assessment. Sampling for HCNp was performed at the seedling stage (3–4 leaf stage) and again at the early vegetative stage (7–8 leaf stage), with tissues from the youngest fully expanded leaf collected from each plant. Dhurrin was estimated at the seedling stage for only a single replicate within one of the plantings. All plant tissue samples were processed individually and randomly assigned to plates to control for technical variation. Blocks were nested within plantings to account for potential variation.

Plates were kept on ice during sample collection and stored at –20 °C. Cyantesmo test strips (Macherey-Nagel) were cut to fit each plate, applied across alternating rows, and sealed with Axygen microplate film (Corning). Pressure was applied to each plate to limit gas transfer between wells and plates were incubated at –35 °C for 20 minutes. Strips were removed from the plates and imaged with a flatbed scanner. Images were converted to the CIELAB color space and regions of interest (ROI) were defined around each blue reaction area, with each ROI representing a single sample. ROIs were thresholded on the b* [0–128] and L* (128–255) channels. Blue intensity was calculated on a per pixel basis by subtracting the pixel L* value from 255. Blue intensity values within a ROI were summed and the resulting value was normalized by ROI area to quantify HCNp.

Dhurrin concentration was quantified using ultrahigh-performance liquid chromatography coupled with tandem mass spectrometry (UHPLC-MS/MS). Leaf samples were harvested, dried at 60 °C for approximately three days, and ground using a Bead Ruptor Elite tissue grinder (6 cycles at 4 m/s, 15 s each, with 10 s breaks). A 100 mg aliquot of ground tissue was extracted with 750 μL of 50% methanol (v/v methanol:diH2O), incubated in a 75 °C water bath for 15 minutes, cooled to room temperature (10–15 min), and supplemented with an additional 750 μL of 50% methanol. Samples were centrifuged at 11,000 rpm for 5 minutes. A 30 μL aliquot of the supernatant was diluted with 270 μL of LC-MS grade water (final 5% methanol, dilution factor 0.15 mL/mg), transferred to a 2 mL glass HPLC vial, and stored at –80 °C. Quality control (QC) was maintained using pooled QC samples prepared from aliquots of individual extracts. Blank extractions were also included. Samples were stored at –80 °C until analysis.

One microliter of extract was injected into an LX50 UHPLC system (PerkinElmer) equipped with a 20-μL sample loop (partial loop injection mode). Chromatographic separation was performed using an ACQUITY UPLC HSS T3 column (1 × 50 mm, 1.8 μm; Waters) maintained at 45 °C. The mobile phases consisted of water with 0.1% formic acid (A) and 100% acetonitrile (B). The elution gradient started at 1% B for 0.5 min, ramped linearly to 99% B over 4.5 min, returned to 1% B at 5.2 min, and was followed by a 2.8 min re-equilibration, for a total runtime of 8 min. The flow rate was set to 400 μL/min. Detection was carried out on a PerkinElmer QSight 420 triple quadrupole mass spectrometer equipped with an electrospray ionization (ESI) source operating in positive mode and selected reaction monitoring (SRM). The optimized SRM transitions for dhurrin were based on an authentic standard: Q1/Q3 = 333.9/144.9 for quantification and 333.9/184.9 for qualification. Source parameters included: drying gas temperature at 120 °C, hot-surface induced desolvation (HSID) temperature at 200 °C, electrospray voltage at 5000 V, and nebulizer gas flow at 350. MS acquisition was scheduled around a retention time of 1.14 min with 1 min time windows. The dwell time for each transition was 100 ms.Data acquisition and processing were performed using Simplicity 3Q™ software (v3.0, PerkinElmer). Quantification was based on standard curves generated from authentic dhurrin standards using linear regression. Concentrations were expressed as μg/g fresh weight, adjusted for sample weight and dilution factors.

### Whole plant phenotyping

Across the pangenome, we observed substantial phenotypic and molecular variation. Field-based measurements collected in 2023 and 2024 across multiple locations captured key agronomic traits including plant height, flowering time, panicle architecture, and yield components, with data separated by replication and site to enable genotype × environment interaction analyses. High-resolution temporal vegetation indices, extracted from drone-based overhead imagery, further characterized canopy development and biomass dynamics over the growing season. Controlled-environment phenotyping complemented field trials to quantify water use efficiency and drought response under well-watered and water-limited conditions using automated indoor imaging systems. Growth curve data across treatments enabled classification of genotypes by their drought response strategies, including early vigor and maintenance of growth under stress. Greenhouse-based evaluations provided additional trait measurements under standardized conditions, including early vigor and developmental timing. Visual documentation of phenotype diversity was collected across environments, including a curated set of representative field and greenhouse images for each genotype. Additional datasets, such as ionomic profiles, root traits, and Li-Cor-based gas exchange measurements, are being compiled to further link physiological traits to structural genomic variation. These multimodal datasets form the foundation for integrative analyses such as principal component analysis (PCA), enabling projection of the pangenome lines in trait space relative to the broader diversity panel. Together, these data highlight phenotypic axes of variation relevant to both local adaptation and global breeding priorities.

### Dhurrin and Cyanide genome-wide associations

Genome-wide association studies (GWAS) were performed using a linear mixed model (LMM) implemented in GEMMA v0.98.3 ^131^. Only directly called genotypes were used; no imputation was performed. Variants were filtered to retain those with a minor allele frequency (MAF) ≥ 0.05. For each trait, association statistics were computed using the Wald test. Markers with missing or undefined p-values were excluded from downstream analysis. The filtered results were used to generate Manhattan plots and assess overlap with a priori candidate genes.

### In Silico Ligand Docking of Tyrosine to CYP79A1 Protein Variants

To evaluate the impact of sequence variants on the binding affinity of CYP79A1 for its substrate tyrosine, in silico ligand docking was performed using SwissDock (https://www.swissdock.ch). Homology models of wild-type and mutant CYP79A1 proteins were generated via SWISS-MODEL (https://swissmodel.expasy.org) using FASTA sequences as input. The resulting PDB files were manually cleaned to remove water molecules and extraneous heteroatoms. The MOL2 file for the substrate L-tyrosine was downloaded from the PubChem database and used as the ligand for docking. PDB files of CYP79A1 containing either an alanine or valine at position 211 were used as the target structures. Each docking run was performed under default settings, using the “Docking with AutoDock Vina” option to scan for potential ligand-binding regions. SwissDock returned a series of clusters, each associated with an estimated binding free energy (ΔG) in kcal/mol. The binding affinity of tyrosine to each CYP79A1 variant was compared by identifying the lowest predicted ΔG value across all clusters. More negative ΔG values were interpreted as indicative of stronger substrate binding. Predicted docking structures were visualized using PyMOL (https://www.pymol.org/) in order to confirm that the substrate tyrosine localized in the active site of each protein model.

### Dhurrin BGC Cis-Element Analysis

To test whether intergenic variants associated with seedling dhurrin content reside in regulatory regions with accessible chromatin and potential transcription factor (TF) binding sites, the Plant PAN 4.0 and SorghumBase ^132^ resources were used. Genomic coordinates of intergenic variants within the BGC were examined using the Ensembl Plants genome browser hosted through SorghumBase to assess whether they overlapped with known accessible chromatin regions (ACRs). Specifically, the ACR track derived from ATAC-seq profiling of 7-day-old Sorghum bicolor leaf tissue, published by Oka et al. (2022), was used, which provides a high-resolution map of chromatin accessibility across the genome. To test whether these intergenic variants also overlapped with putative TF binding sites, each variant sequence was extracted along with 10 bp of flanking sequence on both sides. These sequences were input into PlantPAN 4.0, a promoter and cis-element prediction tool that integrates binding site data from multiple plant-specific motif databases. Only predicted TF binding sites that directly overlapped the variant position were retained for downstream analysis, under the assumption that such overlap may indicate mutation of a functional cis-regulatory element. To maximize confidence in predicted TF-DNA interactions, only hits that included both a known TF family and a corresponding Arabidopsis gene identifier were considered for downstream analysis.

## ACKNOWLEDGEMENTS

The work (proposals: 503014, 504730, 2015; Award DOIs: 10.46936/10.25585/60001093, 10.46936/10.25585/60001224, 10.46936/10.25585/60001015) conducted by the U.S. Department of Energy Joint Genome Institute (https://ror.org/04xm1d337), a DOE Office of Science User Facility, is supported by the Office of Science of the U.S. Department of Energy operated under Contract No. DE-AC02-05CH11231. This work was also supported by Gates Foundation projects: “Green Evolution - Accelerating Dryland Cereals Improvement for Africa” (INV-053669) to GM and the “Sorghum Genomics Toolkit” to TM, GM, DF, JFR, and JS and “Mining useful alleles for climate change adaptation from CGIAR gene banks”. The information presented herein was funded in part by the Advanced Research Projects Agency-Energy (ARPA-E), U.S. Department of Energy, under Award Numbers DE-AR0000594. SMB, DK and TT acknowledge support from NIOO via grant OPP1082853 via grant OPP1082853 “RSM Systems Biology for Sorghum”. PGL was supported by the US Cooperative Extension Service through the Division of Agriculture and Natural Resources of the University of California and DOE Grant DE-SC0014081. This study is made possible by the support of the American People provided to the Feed the Future Innovation Lab for Collaborative Research on Sorghum and Millet through the United States Agency for International Development (USAID) under associate award no. AID-OAA-A13-00047, “Feed the Future Innovation Lab for Genomics-Assisted Sorghum Breeding” and “Feed the Future Innovation Lab for Crop Improvement”. MEH is supported by the DOE Center for Advanced Bioenergy and Bioproducts Innovation (U.S. Department of Energy,Office of Science, Office of Biological and Environmental Research under award no. DE-SC0018420). The contents are the sole responsibility of the authors and do not necessarily reflect the views of USAID or the United States Government. We dedicate this work to the memory of Dr. Todd C. Mockler, a founding contributor to the sorghum pangenome effort. We honor his vision by advancing this work and building on the groundwork he helped establish. His loss is deeply felt, and his legacy lives on through the data, tools, and collaborations he inspired.

## DATA AVAILABILITY

Reference genome assembly and annotation files of all pangenome members are available at https://phytozome-next.jgi.doe.gov/. All raw sequence reads are currently being deposited in the NCBI SRA database. BioProjects will be listed in Supplementary Table 1 (assemblies and annotations), and Supplementary Data 3 (resequencing diversity panel) once posted. In the meantime, please contact the authors for pre-publication access to the raw read data.

## DESCRIPTION OF SUPPLEMENTARY INFORMATION

**Supplementary Information** | Document with Supplementary Tables 1-3 and Supplementary Figures 1-8.

**Supplementary Note 1** | **Phenotypic description of each of the pangenome members**. Includes source photographs that had backgrounds removed for Fig. 1 and information related to pedigree, botanical type and other genotype-specific characteristics.

**Supplementary Data 1** | **Syntenic blocks between V3 and V5 BTx623**. Minimap2 was used to align the V3 (query) to the V5 (target) assemblies using the default for closely related whole-genome alignments (−xasm5). All other parameters were default. Data presented as the raw .paf file with standard column names. File name: “SIData1_v3vsv5_mm2syntenicBlocks.paf”.

**Supplementary Data 2** | **Synteny and global orthologs between V3 and V5**. Annotated .bed files containing orthofinder- and GENESPACE-defined orthogroups (with columns globOG = global orthogroups [deprecated], globHOG = global phylogenetic hierarchical orthogroups ‘HOGs’, og = synteny-constrained HOGs), tandem arrays (columns arrayID, arrayRep), and information about the genes (e.g. peptide length). File name: “SIData2_v3vsv5_genespaceOrthofinderOrthogroups.txt”.

**Supplementary Data 3** | **Metadata for the diversity panel**. Sample and library information for all members of the diversity panel, with flags showing which libraries were used for each analysis. File name: “SIData3_diversityPanelMetadata.csv”

**Supplementary Data 4** | **High-throughput phenotypic characterization of the pangenome members**. Minimap2 was used to align the V3 (query) to the V5 (target) assemblies using the default for closely related whole-genome alignments (−xasm5). All other parameters were default. Data presented as the raw .paf file with standard column names. File name: “SIData4_PlantCVDeltaWueHeightArea.csv”.

**Supplementary Data 5** | **Structural variants among reference genomes**. Summary of SVs identified from Minimap2 alignments with SyRI at 3 minimum variant lengths (50bp, 10kb, 1Mb) for each of the 32 reference genomes aligned to BTx623 V5. Counts and total lengths are provided for deletions, insertions, duplications, inversions, inverted duplications, inverted translocations, and translocations. File name: “SIData5_SyRI_SV_summary.txt”.

**Supplementary Data 6** | **Pangenome annotation metadata**. Library ID, number of reads, condition/tissue and reference genomeID for all RNA sequencing libraries used for protein-coding gene annotation. File name: “SIData6_RNAseq.libraryStats_aligned2Refs.csv”

**Supplementary Data 7** | **Pangenome pangene sets**. Format following Supplementary Data 2 but for all 33 members of the pangenome. File name: “SIData7_pangenomeAnnot_genespaceOrthofinderOrthogroups.txt”.

**Supplementary Data 8** | **Gene ontology enrichment tests**. Standard GO enrichment format with the final column “scale” giving the spatial scale in km with which the outliers were defined for 15 wavelet scales. File name: “SIData8_waveletGOenrichmentStats.csv”.

**Supplementary Data 9** | **Genome wide association mapping results for dhurrin concentration**.

Gzip-compressed text file containing the output of GEMMA in default format, which contains results in fields: chromosome, strand [not filled], bp position, number of missing calls, alternative allele, reference allele, allele frequency, beta statistic, standard error, and the following statistics (see GEMMA manual): logl_H1, l_remle, l_mle, p_wald, p_lrt, p_score. File name: “SIData9_GEMMMAdhuSamp.assoc.txt.gz”.

**Supplementary Data 10** | **Genome wide association mapping results for seedling hydrogen cyanide potential (HCNp)**. See description for Supplementary Data 9. File name: “SIData10_GEMMMASeedHCNSamp.assoc.txt.gz”.

**Supplementary Data 11** | **Genome wide association mapping results for vegetative hydrogen cyanide potential (HCNp)**. See description for Supplementary Data 9. File name: “SIData11_GEMMMAVegHCNSamp.assoc.txt.gz”.

