## Supplementary Information for "Developing future resilience from signatures of adaptation across the sorghum pangenome"

**Running title:** Sorghum pangenome climate adaptation


**SUPPLEMENTARY FIGURES** **2**

Supplementary Figure 1 2

Supplementary Figure 2 3

Supplementary Figure 3 4

Supplementary Figure 4 5

Supplementary Figure 5 6

Supplementary Figure 6 7

Supplementary Figure 7 8

Supplementary Figure 8 9

**SUPPLEMENTARY TABLES** **10**

Supplementary Table 1 10

Supplementary Table 2 12

Supplementary Table 3 13

### SUPPLEMENTARY FIGURES

#### Supplementary Figure 1

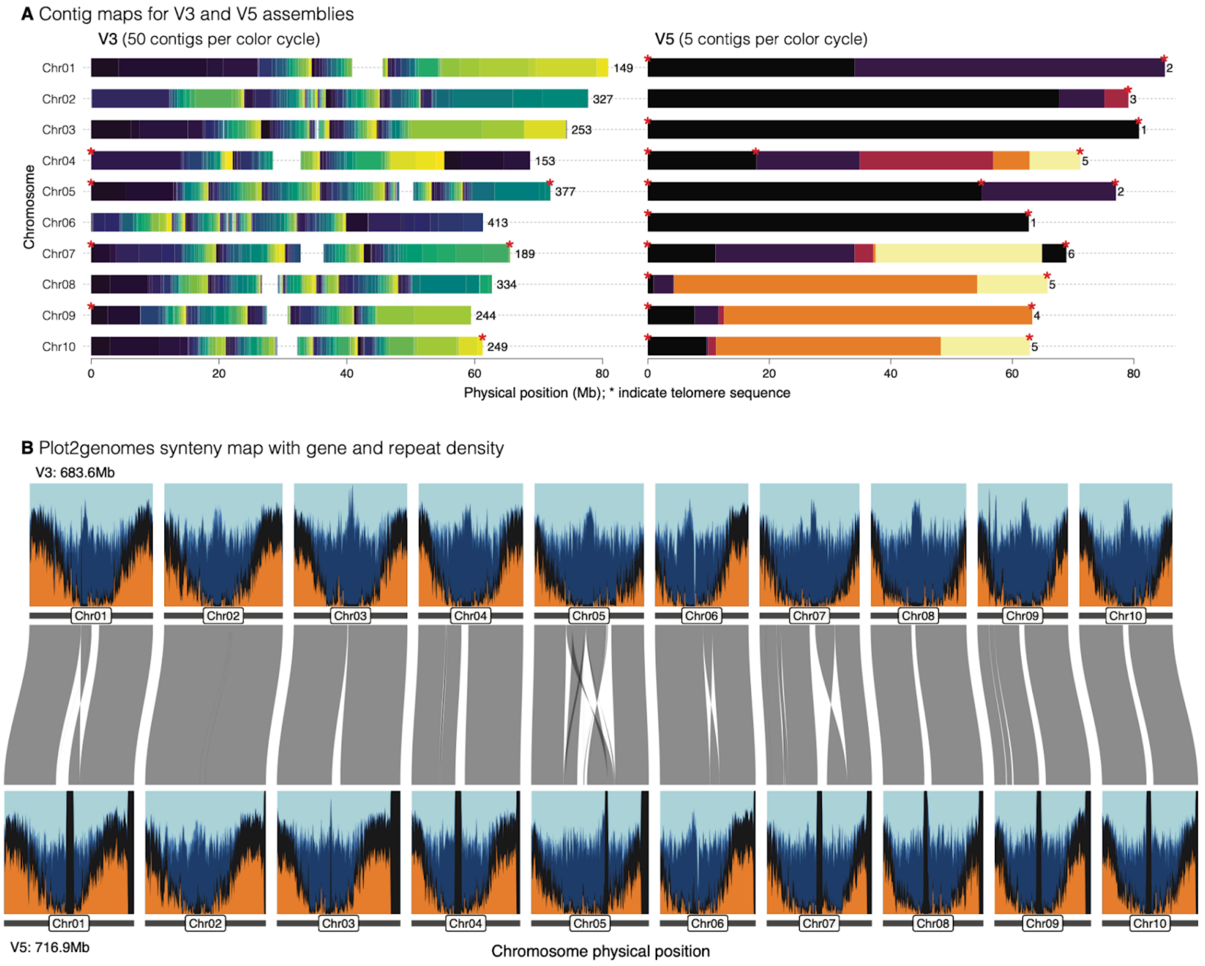


**Supplementary Figure 1 | Genome assembly and annotation comparison between V3 and V5 BTx623 resources. A** Contig maps showing the position of gaps and telomeres in V3 (left) and V5 (right) illustrate an approximate 100-fold increase in assembly quality between the versions. Red *s indicate the presence of dense blocks of putative telomeric repeats. **B** Synteny map accompanied by gene (orange) and repeat density (blues: dark = T3, light = other; 100kb-overlapping 500kb sliding windows) show the position of structural variants and increased repeat representation between the two versions.

#### Supplementary Figure 2

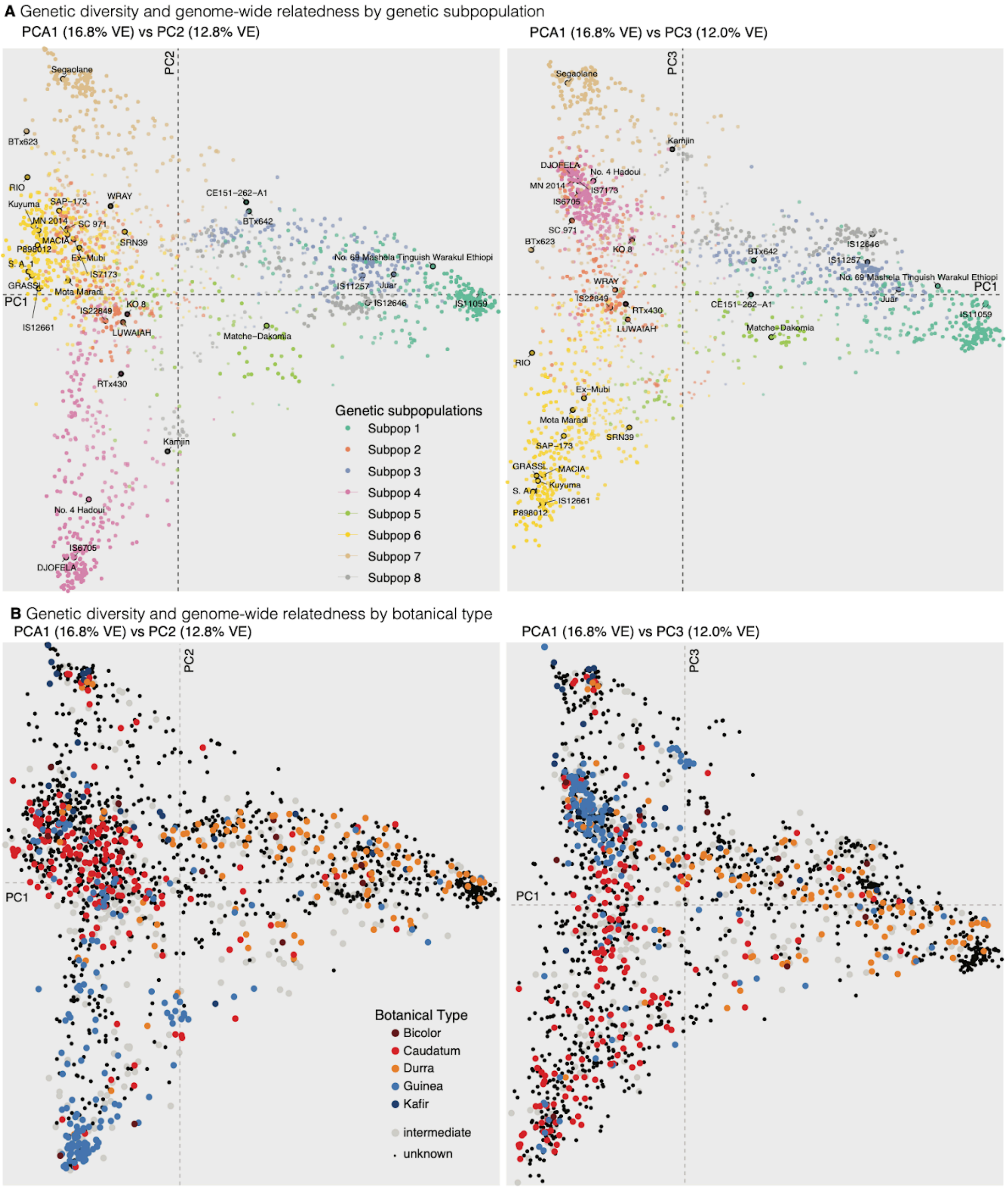


**Supplementary Figure 2 | Genetic principal components.** Coordinates from SNP- and short INDEL-based principal components are shown for the first two axes (left) and first and third axes (right), color-coded by the eight admixture subpopulations (**A)** and the five major botanical types (**B**).

#### Supplementary Figure 3

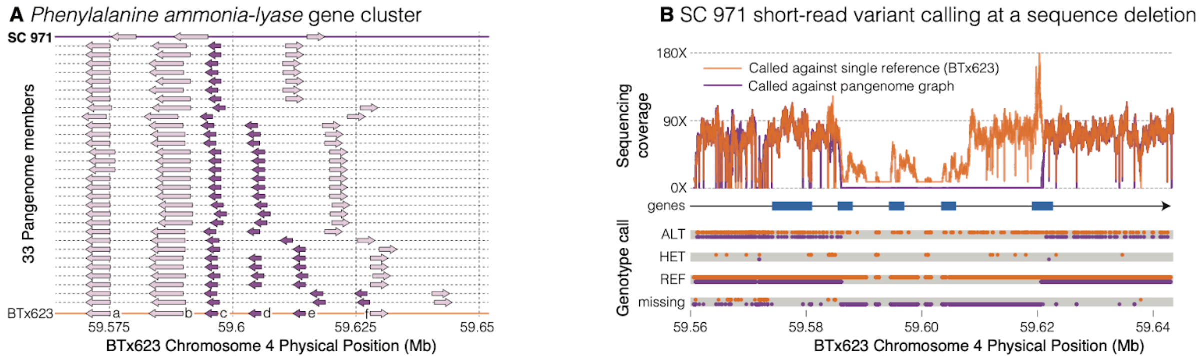


**Supplementary Figure 3 | Comparison of graph and single-reference genotyping approaches at a known presence-absence variation site. A** Gene arrow plot indicating presence/absence of phenylalanine ammonia lyase copies on chromosome 4 across the 33 pangenome members. The BTx623 reference and SC 971, which harbors the largest deletion of three . From left to right the seven gene models in BTx623 are: a: Sobic.004G220300, b: Sobic.004G220400, c: Sobic.004G220500, d: Sobic.004G220600, e: Sobic.004G220650, and f: Sobic.004G220700. PI numbers (or genome IDs) from bottom to top: BTx623, PI656057, PI597980, PI656044, PI656027, PI565121, PI276837, PI513676, PI569459, RTx430, Wray, PI276816, PI329501, PI656023, PI180348, PI534133, PI655981, PI660557, PI533766, PI660565, PI585966, PI570071, PI156178, PI660563, PI656031, PI656015, PI655988, PI576434, PI154844, BTx642, PI656050, PI656111. Arrow directions indicate gene orientations on Chr04. Positions are provided in the BTx623 coordinate system, as BTx623 is the primary reference in the pangenome graph. Positions of genes in other genomes are approximate. Gene PAV was inferred using odgi untangle and a minimum Jaccard similarity of 0.8 with respect to BTx623 gene sequences. **B** Comparison of depth and genotype calls between linear and graph alignments of phenylalanine ammonia lyase array on sorghum chromosome 4. To generate genotype calls, short reads were aligned to both the linear V5 BTx623 reference genome (BWA) and the Minigraph-Cactus pangenome graph with 33 haplotypes and BTx623 as the primary reference (vg giraffe). Graph-aligned reads were ‘surjected’ to BTx623 coordinate space and an identical variant-calling workflow was applied to linear- and graph-based BAM files. Concordant with the PAV results, graph-based alignments produce missing calls for SC 971 at genes c-e, whereas linear-based alignments primarily produce homozygous reference genotype calls at these loci. Additionally, linear-based alignments produced heterozygous calls not observed in the graph-based calls across the region, likely due to the reference bias inherent in linear-based analyses.

#### Supplementary Figure 4

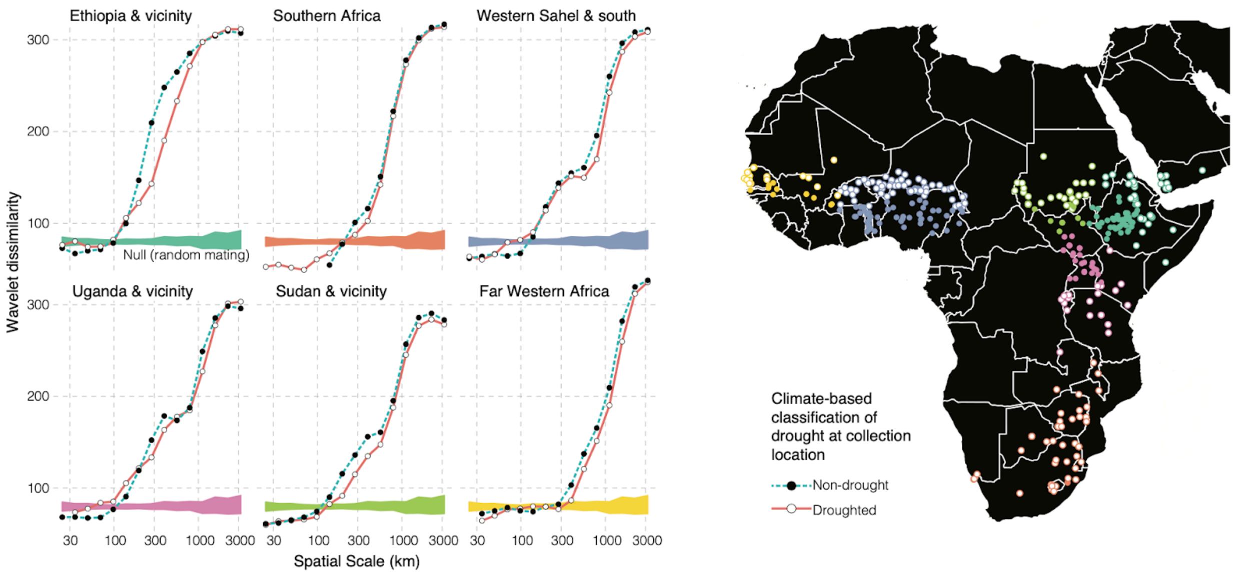


**Supplementary Figure 4 | Drought-associated differentiation in discrete geographic regions.** Plots on the left follow that of Fig. 4A,D, but are split by six PAM-clustered geographic groups (see map on right for positions of groupings).

#### Supplementary Figure 5

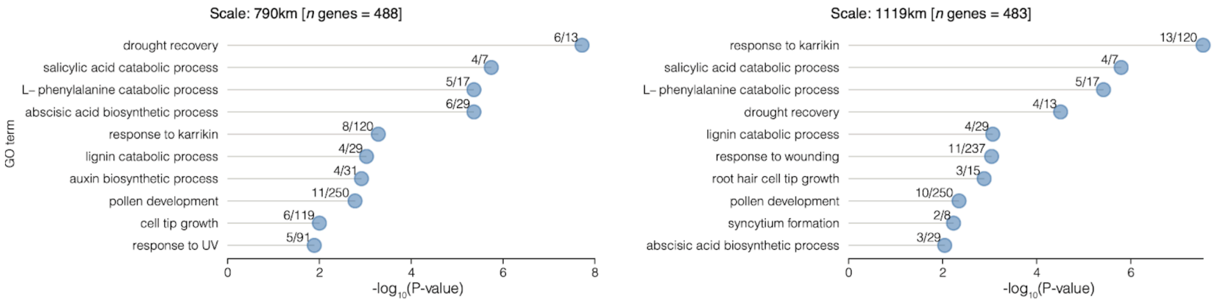


**Supplementary Figure 5 | Summary of the top 10 gene ontology enrichments for 800-1200km scales.** Standard GO enrichment plots showing the significance (x-axis) and contribution of genes (numbers next to points) for the top 10 enrichments for wavelet dissimilarity scans at 790km (left) and 1,119km (right).

### Supplementary Figure 6

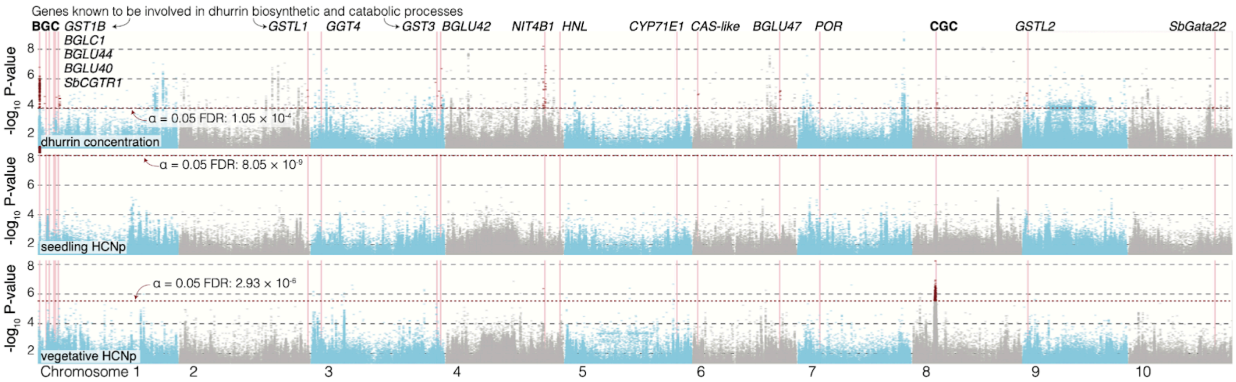


**Supplementary Figure 6 | Genome-wide dhurrin and HCNp GWAS scans.** ‘Manhattan’ plots for dhurrin concentration, total seedling and vegetative cyanide potential. The biosynthetic gene cluster (BGC), catabolic gene cluster (CGC) and 17 genes in the dhurrin pathways are highlighted. Significant marker associates within 1Mb of these genes are colored dark red.

#### Supplementary Figure 7

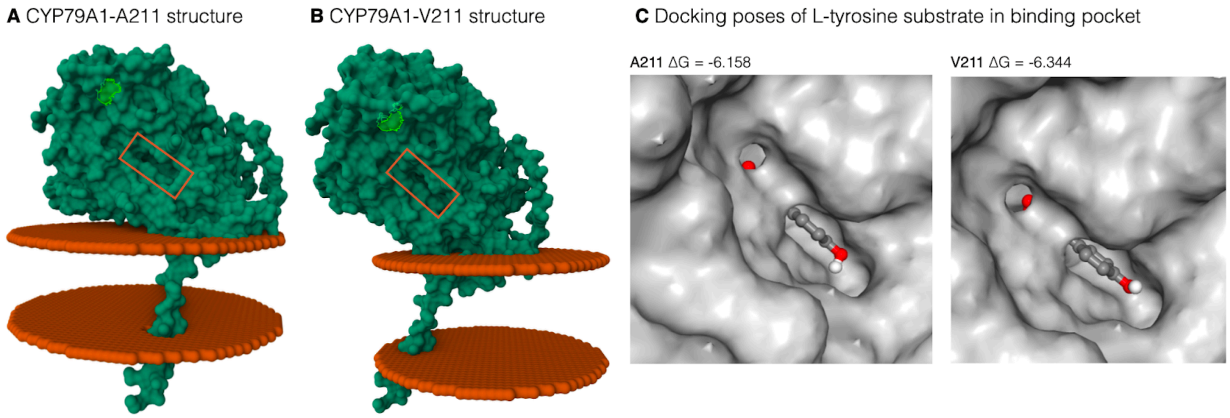


**Supplementary Figure 7 | Structural and regulatory context of a function-altering variant in CYP79A1.** Homology models of the CYP79A1 enzyme containing either an **A** alanine (A211) or **B** valine (V211) residue at position 211 were embedded within the ER membrane using OPM PPM3.0. In each model, residue 211 is outlined in green and the substrate-binding pocket for tyrosine is boxed in red. **C** Predicted docking poses of the substrate L-tyrosine in the binding pocket of CYP79A1 A211 (left) and V211 (right) variants. Ligand docking was performed using SwissDock with the AutoDock Vina protocol. The most energetically favorable pose for each variant is shown, along with the predicted binding free energy (ΔG, kcal/mol), where more negative values indicate stronger binding affinity.

#### Supplementary Figure 8

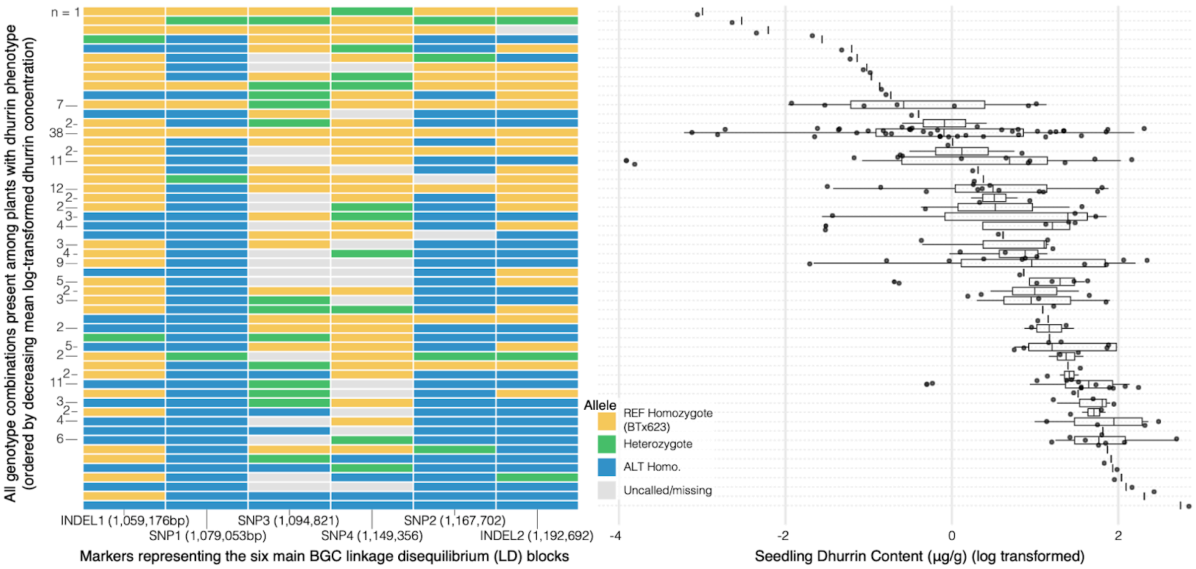


**Supplementary Figure 8 | Phenotypic effects of resequencing short variant allelic combinations.** All combinations of genotypes (BTx623 genotype homozygote is yellow, heterozygote is green, alternative homozygote is blue, missing genotype call is gray) across six relatively unlinked sites in the BGC and the log-transformed seedling dhurrin content of plants with those genotype combinations. INDEL1-2 and SNP1-2 are the same as in Figure 5C. Here, we include two additional SNPs that show high levels of missingness (SNP3-4: Chr01 1,094,821, 1,149,356). The number of phenotyped samples with a given allelic combination is denoted to the left (unlabeled rows have only a single library with that combination of genotypes).

### SUPPLEMENTARY TABLES

#### Supplementary Table 1

**Supplementary Table 1 | Genome assembly and annotation statistics.** Combined statistics and genome IDs / common names for the 31 new reference genomes presented here and *two previously published genomes used in the pangenome resource.

| Genome Name | Common Name | Genome (Mb) | Scaffold N50 (Mb) | Contig N50 (Mb) | Genome in chromosomes (%) | n. primary transcripts | n. alt. transcripts | Annotation BUSCO (%) |
|---|---|---|---|---|---|---|---|---|
| BTx623 | BTx623 | 719.9 | 71.2 | 50.7 | 99.59 | 35,052 | 12,145 | 99.7 |
| BTx642* | BTx642 | 680.1 | 64.5 | 12.1 | 99.45 | 35,115 | 14,233 | 99.4 |
| Wray | Wray | 736.8 | 72 | 63.9 | 98.72 | 30,610 | 24,639 | 99.4 |
| RTx430* | RTx430 | 677.7 | 64.6 | 4 | 98.09 | 34,601 | 12,280 | 99.2 |
| PI_329301 | IS 11059 | 683.3 | 64.1 | 15.7 | 98.93 | 28,942 | 10,981 | 99.7 |
| PI_156178 | MN 2014 | 681.1 | 65.3 | 20 | 99.62 | 29,002 | 12,066 | 99.4 |
| PI_180348 | Juar (IS 12876) | 690.6 | 68.7 | 15.1 | 99.33 | 28,924 | 12,177 | 99.6 |
| PI_276837 | IS 12661 | 693.7 | 64.9 | 18.7 | 98.94 | 29,687 | 12,468 | 99.7 |
| PI_569459 | LUWAIAH (IS 19523) | 716.2 | 65.6 | 5.3 | 95.01 | 29,924 | 12,100 | 99.7 |
| PI_570071 | IS 22849 | 667.5 | 64.7 | 4.3 | 98.33 | 28,385 | 11,753 | 98.2 |
| PI_656031 | IRAT204 | 741.4 | 65.2 | 20.7 | 92.8 | 30,537 | 12,824 | 99.6 |
| PI_656044 | Kuyuma | 686.5 | 64.6 | 14.1 | 98.95 | 28,959 | 11,433 | 99.6 |
| PI_154844 | GRASSL | 701.2 | 65.4 | 28.4 | 99.02 | 29,739 | 11,917 | 99.4 |
| PI_276816 | IS 12646 | 699.3 | 65.7 | 21.4 | 98.85 | 30,355 | 11,801 | 99.5 |
| PI_329501 | IS 11257 | 765.8 | 65.4 | 2.6 | 89.84 | 31,566 | 11,821 | 99.2 |
| PI_513676 | Matche-Dakomia (IS 20380) | 688.8 | 65.1 | 29.3 | 99.06 | 29,556 | 11,236 | 99.4 |
| PI_533766 | SC 265 | 687.8 | 65.6 | 22.4 | 99.35 | 29,614 | 12,439 | 99.3 |
| PI_534133 | SC 35 | 688.6 | 66.6 | 15.4 | 98.86 | 29,931 | 12,386 | 99.4 |
| PI_565121 | MACIA | 689.1 | 65.2 | 26.1 | 99.86 | 29,238 | 11,807 | 99.7 |
| PI_576434 | SC 1103 | 688.7 | 65.6 | 20.1 | 99.04 | 29,381 | 12,068 | 99.5 |
| PI_585966 | DJOFELA (IS 26395) | 681.7 | 64.5 | 9.5 | 98.72 | 28,332 | 10,781 | 97.7 |
| PI_597980 | SC 1345 (CSM-90) | 692.1 | 65.9 | 23 | 99.15 | 29,153 | 12,008 | 99.6 |
| PI_655981 | CSM-63 | 692.1 | 65.2 | 22 | 99.38 | 29,231 | 12,377 | 99.5 |
| PI_655988 | COMBINE KAFIR-60 | 696.8 | 66.3 | 24.3 | 99.33 | 28,761 | 10,279 | 99.7 |
| PI_656015 | Ajabsido | 689.4 | 64.4 | 22.6 | 98.68 | 29,779 | 11,923 | 99.6 |
| PI_656023 | Segaolane | 696.2 | 66 | 27.9 | 99.28 | 29,639 | 12,074 | 99.4 |

| | | | | | | | | |
|---|---|---|---|---|---|---|---|---|
| PI_656027 | SRN39 | 696.7 | 65.4 | **18**.3 | 99.32 | 29,589 | 11,520 | 99.7 |
| PI_656050 | Mota Maradi | 698.8 | 65.8 | 20.9 | 99.61 | 29,337 | 12,294 | 99.5 |
| PI_656057 | P898012 | 694.7 | 65.8 | 17.8 | 99.1 | 29,332 | 12,140 | 99.4 |
| PI_656111 | SC 971 | 690.9 | 64.7 | 12.1 | 98.63 | 29,662 | 12,432 | 99.4 |
| PI_660557 | IS 6705 | 690.1 | 65.9 | 21.5 | 99.04 | 29,411 | 11,907 | 99.3 |
| PI_660563 | IS 7173 | 698.1 | 65.5 | 27.7 | 99.05 | 29,937 | 11,818 | 99.4 |
| PI_660565 | Kamjin (IS 3620) | 707.6 | 67.5 | 25.4 | 99 | 30,367 | 12,568 | 99.4 |

#### Supplementary Table 2

**Supplementary Table 2**. Illumina RNA-seq and PacBio Iso-Seq data used for genome annotation of 31 new reference genomes presented in this study.

| **Genome Name** | **Illumina RNA-seq read pairs (Billion)** | **PacBio Iso-Seq CCS reads (Million)** |
|---|---|---|
| BTx623 | 4 | 19.73 |
| Wray | 4 | 24 |
| PI_329301 | 0.79 | 7.65 |
| PI_156178 | 0.76 | 9.8 |
| PI_180348 | 0.9 | 8.61 |
| PI_276837 | 0.73 | - |
| PI_569459 | 0.27 | 9.94 |
| PI_570071 | 1.83 | 9.95 |
| PI_656031 | 0.7 | - |
| PI_656044 | 0.71 | 6.04 |
| PI_154844 | 0.67 | - |
| PI_276816 | 1.45 | - |
| PI_329501 | 0.95 | 7.72 |
| PI_513676 | 0.81 | - |
| PI_533766 | 0.74 | - |
| PI_534133 | 1.19 | - |
| PI_565121 | 0.84 | - |
| PI_576434 | 0.61 | - |
| PI_585966 | 0.59 | 6.9 |
| PI_597980 | 0.9 | - |
| PI_655981 | 0.79 | - |
| PI_655988 | 0.74 | 5.72 |
| PI_656015 | 0.76 | - |
| PI_656023 | 0.86 | - |
| PI_656027 | 0.88 | - |
| PI_656050 | 0.71 | - |
| PI_656057 | 0.8 | - |
| PI_656111 | 0.79 | - |
| PI_660557 | 0.78 | - |
| PI_660563 | 0.92 | - |
| PI_660565 | 1.47 | - |

#### Supplementary Table 3

**Supplementary Table 3**. Predicted transcription factor binding sites and their overlap with regions of accessible chromatin for select intergenic sites strongly associated with the dhurrin seedling content phenotype.

| Site (BTx623 coordinates) | Location | log10(P wald) Seedling dhurrin content | TF Recognition Site(s) Identified through PlantPAN (Y/N) | TF ID or Motif name | Accessible chromatin region overlap with site identified through SorghumBase (Y/N) |
|---|---|---|---|---|---|
| INDEL Chr01:1,059,176bp | Intergenic between Sobic.001G012100 (BGC 5' Border Gene) and *CYP71E1* | -5.76581 | Yes | AT3G14230 (RAP2.2), AT5G65410 (AtHB25), AT1G23420 (INO), AT4G29940 (PRHA), AT1G75240 (AtHB33), AT1G09030 (NF-YB4), AT1G75520 (SRS5) | No |
| SNP Chr01:1,079,053bp | Intergenic between *CYP71E1* and *CYP79A1* | -3.32338 | Yes | AT1G23380 (KNAT6), AT1G77920 (TGA7) | No |
| SNP Chr01:1,094,821bp | Intergenic between *CYP71E1* and *CYP79A1* | | No | NA | No |
| SNP Chr01:1,149,356bp | Intergenic between *CYP71E1* and *CYP79A1* | | Yes | AT2G42540 (AtCOR15A) | No |
| INDEL Chr01:1,185,567bp | Intergenic between *UGT85B1* and *GST1* | -4.876768 | Yes | AT3G19290 (ABF4), AT4G34000 (ABF3) | Yes |
