## Supplementary Note 1 for "Developing future resilience from signatures of adaptation across the sorghum pangenome"

# PI 534133

**Other name(s):** SC 35, No. 69 Mashela Tinguish, Warakul Ethiopi
**Species:** Sorghum bicolor subsp. bicolor
**Group:** ICRISAT Reference Collection, Sorghum Association Panel & Geo-Reference Collection

| | |
|---|---|
| **Flowering Days Under Short Days** | 73 days after planting |
| **Seed Color** | Yellow |
| **Seedling Color** | Green |
| **Mid-Rib Color** | White |
| **Lodging** | No |
| **Stem diameter (mm) 3rd internode** | 31.2, 22.9 |
| **Stem diameter (mm) 6th internode** | 20.2, 18.0 |
| **Main Stem Height: LC; PD (cm)** | 69; 16 |
| **Total Number of Tillers, with panicles** | 5, 5 |
| **Tiller 1 Stem Height: LC; PD (cm)** | 65; 17.5 |
| **Tiller 2 Stem Height: LC; PD (cm)** | 67; 40 |
| **Tiller 3 Stem Height: LC; PD (cm)** | 63; 35 |

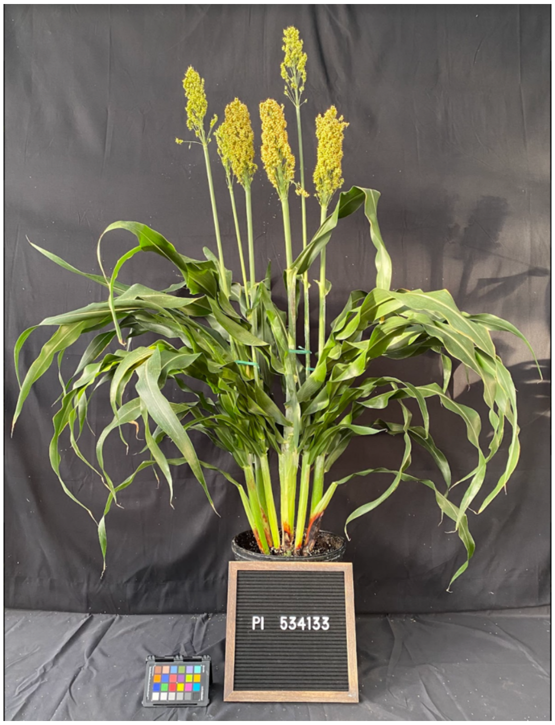


- *Stem diameter: recorded both axes means. if oval-shaped "x, y".*
- *Measurements highlighted in green and photos were recorded 18 weeks after planting (WAP).*
- *Growth conditions: 28°C (day)/22°C (night); 16h light:8h dark; Light Intensity: 400 uMol, 50% RH, Soil type: Berger BM7.*

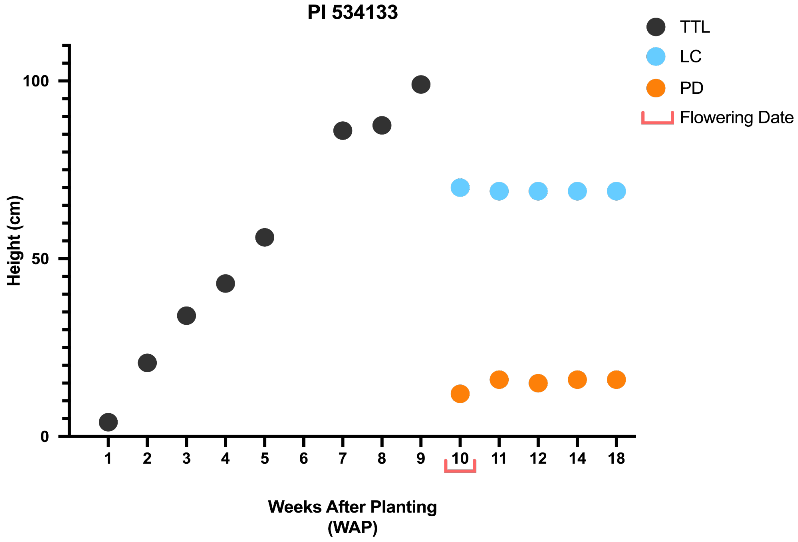


- *The plant was measured up to the last collar (LC) as well as the peduncle to the base of the panicle (PD). Before panicle formation, the height was measured up to the top of the tallest leaf (TTL).*

# PI 656031

**Other name(s):** IRAT204, CE151-262-A1
**Species:** Sorghum bicolor subsp. bicolor
**Group:** ICRISAT Reference Collection & Sorghum Association Panel

| | |
|---|---|
| **Flowering Days Under Short Days** | 75 days after planting |
| **Seed Color** | White |
| **Seedling Color** | Green |
| **Mid-Rib Color** | Tan |
| **Lodging** | No |
| Stem diameter (mm) 3rd internode | 17.2, 15.1 |
| Stem diameter (mm) 6th internode | 13.1 |
| Main Stem Height: LC; PD (cm) | 77; 6.5 |
| Total Number of Tillers, with panicles | 14, 10 |
| Tiller 1 Stem Height: LC; PD (cm) | 67; 24.5 |
| Tiller 2 Stem Height: LC; PD (cm) | 77; 15.5 |
| Tiller 3 Stem Height: LC; PD (cm) | 54; 14 |

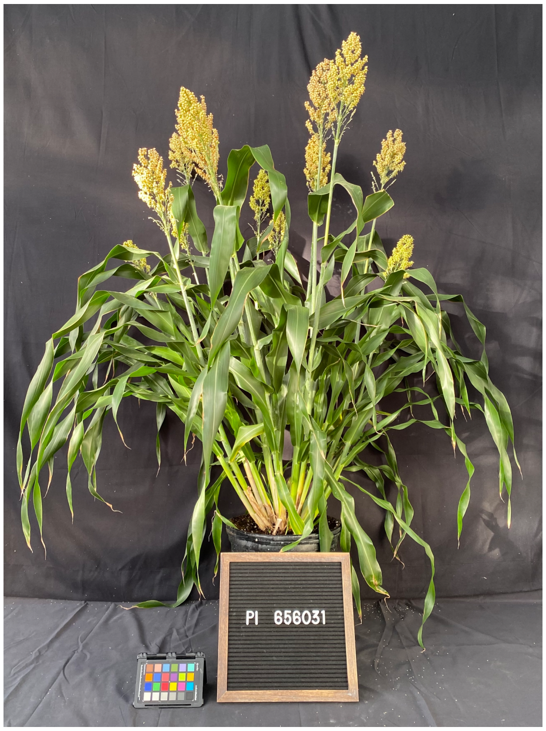


- *Stem diameter: recorded both axes means. if oval-shaped "x, y".*
- *Measurements highlighted in green and photos were recorded 18 weeks after planting (WAP).*
- *Growth conditions: 28°C (day)/22°C (night); 16h light:8h dark; Light Intensity: 400 uMol, 50% RH, Soil type: Berger BM7.*

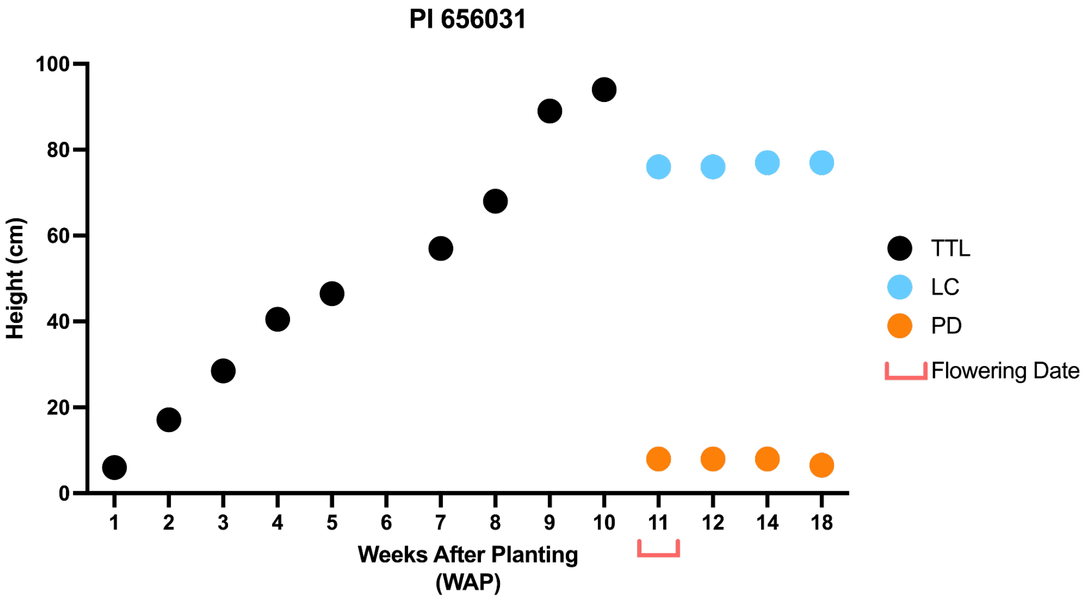


- *The plant was measured up to the last collar (LC) as well as the peduncle to the base of the panicle (PD). Before panicle formation, the height was measured up to the top of the tallest leaf (TTL).*

# PI 513676

**Other name(s):** Matche-Dakomia, IS 20380
**Species:** Sorghum bicolor subsp. bicolor
**Group:** ICRISAT Reference Collection

| | |
|---|---|
| **Flowering Days Under Short Days** | 52 days after planting |
| **Seed Color** | White |
| **Seedling Color** | Green |
| **Mid-Rib Color** | White |
| **Lodging** | No |
| **Stem diameter (mm) 3rd internode** | 12.6 |
| **Stem diameter (mm) 6th internode** | NA |
| **Main Stem Height: LC; PD (cm)** | 90; 28 |
| **Total Number of Tillers, with panicles** | 3, 3 |
| **Tiller 1 Stem Height: LC; PD (cm)** | 94; 22.5 |
| **Tiller 2 Stem Height: LC; PD (cm)** | 99; 22.5 |
| **Tiller 3 Stem Height: LC; PD (cm)** | 105; 12.5 |

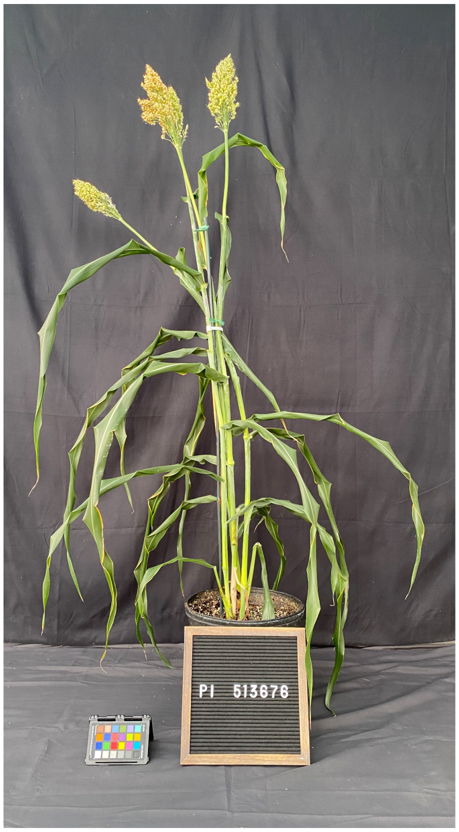


- *Stem diameter: recorded both axes means. if oval-shaped "x, y".*
- *Measurements highlighted in green and photos were recorded 18 weeks after planting (WAP).*
- *Growth conditions: 28°C (day)/22°C (night); 16h light:8h dark; Light Intensity: 400 uMol, 50% RH, Soil type: Berger BM7.*

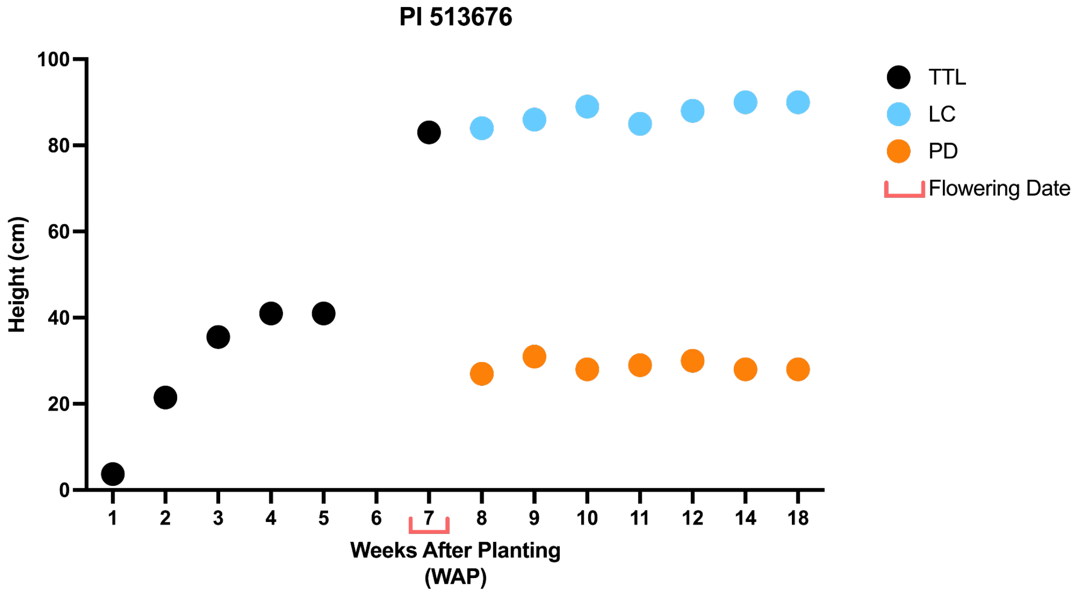


- *The plant was measured up to the last collar (LC) as well as the peduncle to the base of the panicle (PD). Before panicle formation, the height was measured up to the top of the tallest leaf (TTL).*

# PI 660557

**Other name(s):** IS 6705
**Species:** Sorghum bicolor subsp. bicolor
**Group:** ICRISAT Reference Collection

| | |
|---|---|
| **Flowering Days Under Short Days** | 68 days after planting |
| **Seed Color** | Yellow |
| **Seedling Color** | Green |
| **Mid-Rib Color** | White |
| **Lodging** | No |
| **Stem diameter (mm) 3rd internode** | 12.8 |
| **Stem diameter (mm) 6th internode** | 11.7 |
| **Main Stem Height: LC; PD (cm)** | 236; 17 |
| **Total Number of Tillers, with panicles** | 8, 7 |
| **Tiller 1 Stem Height: LC; PD (cm)** | 117; 3 |
| **Tiller 2 Stem Height: LC; PD (cm)** | 173; 21 |
| **Tiller 3 Stem Height: LC; PD (cm)** | 173; 38 |

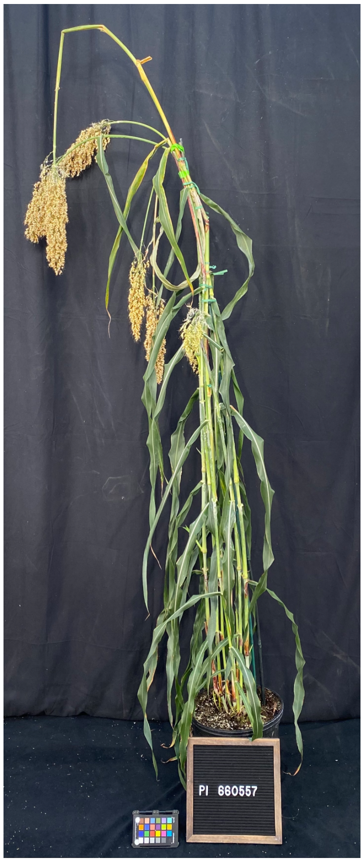


- *Stem diameter: recorded both axes means. if oval-shaped "x, y".*
- *Measurements highlighted in green and photos were recorded 18 weeks after planting (WAP).*
- *Growth conditions: 28°C (day)/22°C (night); 16h light:8h dark; Light Intensity: 400 uMol, 50% RH, Soil type: Berger BM7.*

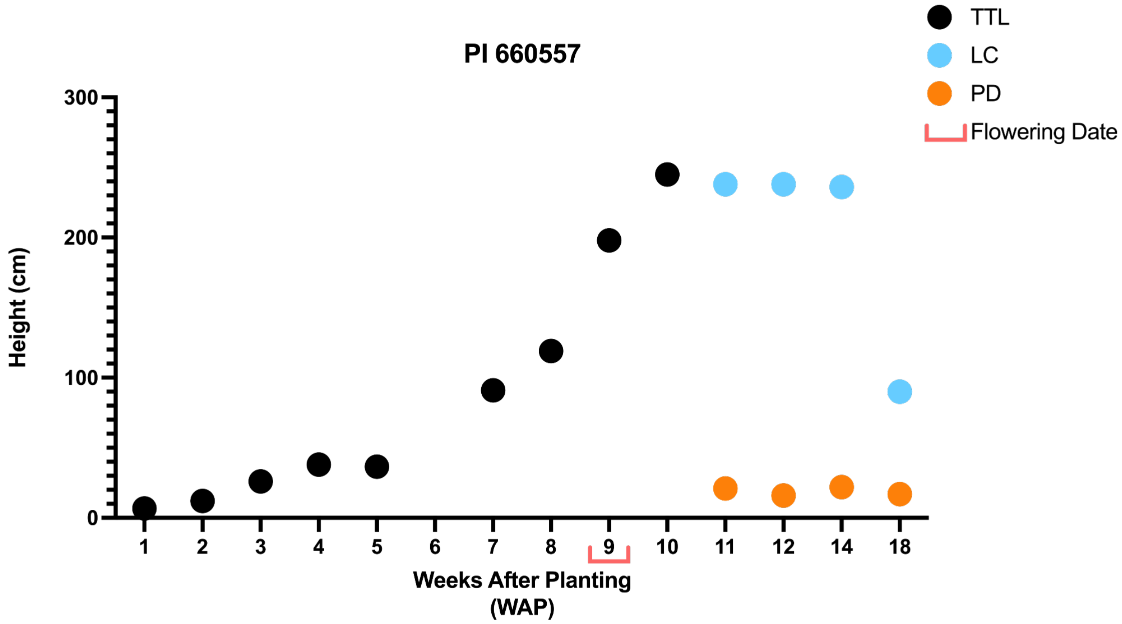


- *The plant was measured up to the last collar (LC) as well as the peduncle to the base of the panicle (PD). Before panicle formation, the height was measured up to the top of the tallest leaf (TTL).*

# PI 533766

**Other name(s):** SC 265, No. 4 Hadoui
**Species:** Sorghum bicolor subsp. bicolor
**Group:** ICRISAT Reference Collection & Sorghum Association Panel

| | |
|---|---|
| **Flowering Days Under Short Days** | 81 days after planting |
| **Seed Color** | Yellow |
| **Seedling Color** | Green |
| **Mid-Rib Color** | Tan |
| **Lodging** | No |
| Stem diameter (mm) 3rd internode | 16.9, 13.1 |
| Stem diameter (mm) 6th internode | 10.7, 8.9 |
| **Main Stem Height: LC; PD (cm)** | 57; 32.5 |
| **Total Number of Tillers, with panicles** | 6, 6 |
| **Tiller 1 Stem Height: LC; PD (cm)** | 45; 33.5 |
| **Tiller 2 Stem Height: LC; PD (cm)** | 45; 23 |
| **Tiller 3 Stem Height: LC; PD (cm)** | 49; 31.5 |

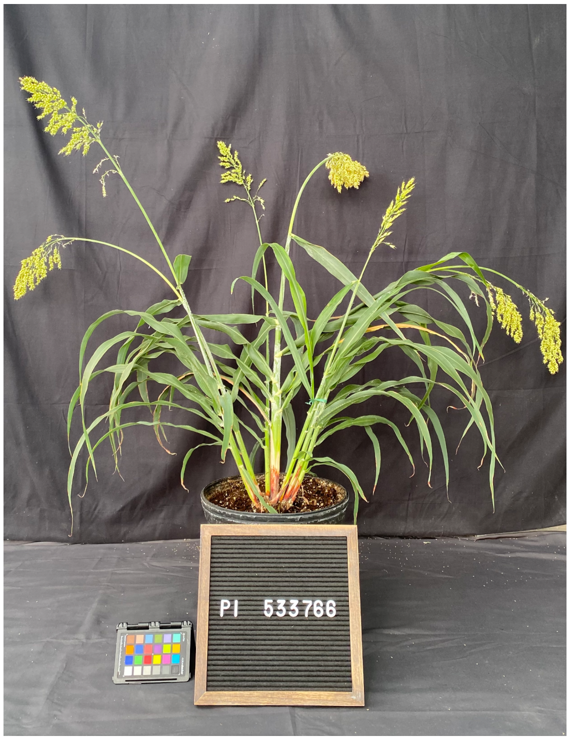


- *Stem diameter: recorded both axes means. if oval-shaped "x, y".*
- *Measurements highlighted in green and photos were recorded 18 weeks after planting (WAP).*
- *Growth conditions: 28°C (day)/22°C (night); 16h light:8h dark; Light Intensity: 400 uMol, 50% RH, Soil type: Berger BM7.*

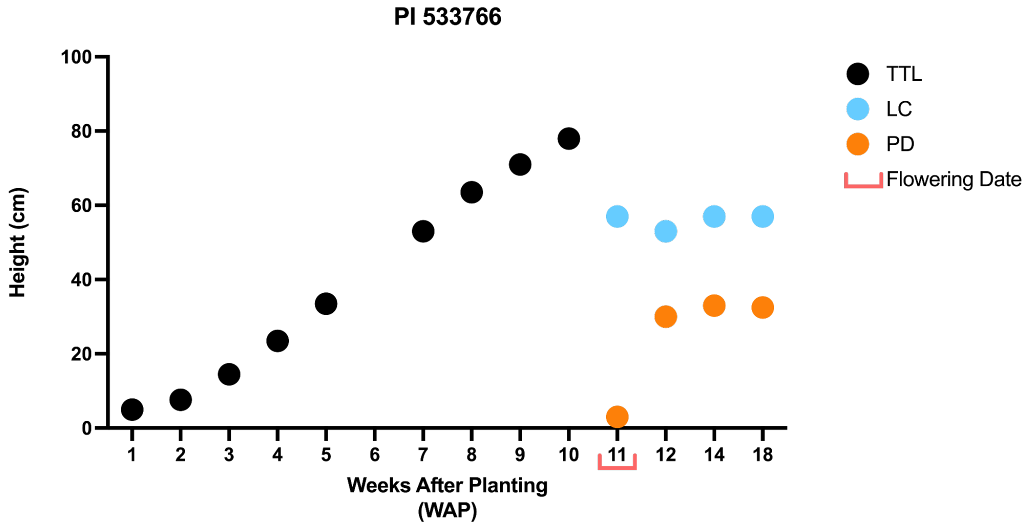


- *The plant was measured up to the last collar (LC) as well as the peduncle to the base of the panicle (PD). Before panicle formation, the height was measured up to the top of the tallest leaf (TTL).*

# PI 655981

**Other name(s):** CSM-63, 87C424
**Species:** Sorghum bicolor subsp. bicolor
**Group:** ICRISAT Reference Collection, Sorghum Association Panel & Bioenergy Association Panel

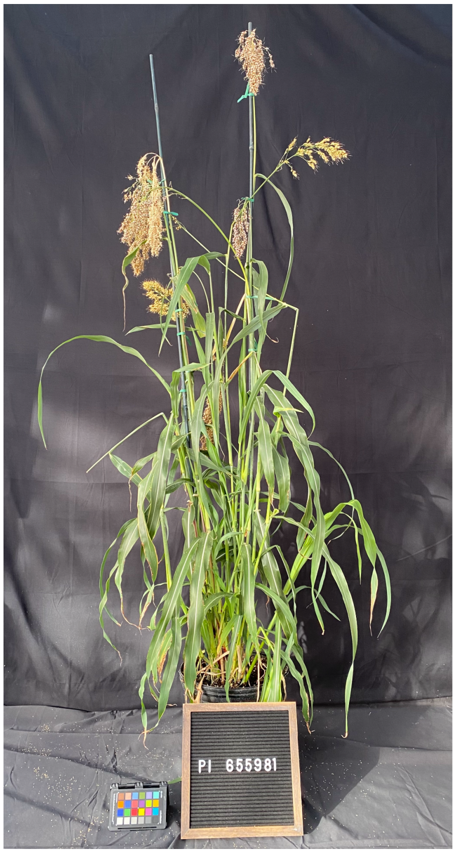


| | |
|---|---|
| **Flowering Days Under Short Days** | 47 days after planting |
| **Seed Color** | Brown-purple |
| **Seedling Color** | Green |
| **Mid-Rib Color** | White |
| **Lodging** | Yes |
| **Stem diameter (mm) 3rd internode** | 6.6 |
| **Stem diameter (mm) 6th internode** | 4.0 |
| **Main Stem Height: LC; PD (cm)** | 130; 27.5 |
| **Total Number of Tillers, with panicles** | 16, 9 |
| **Tiller 1 Stem Height: LC; PD (cm)** | 109; 13 |
| **Tiller 2 Stem Height: LC; PD (cm)** | 116; 16 |
| **Tiller 3 Stem Height: LC; PD (cm)** | 108; 21 |

- *Stem diameter: recorded both axes means. if oval-shaped "x, y".*
- *Measurements highlighted in green and photos were recorded 18 weeks after planting (WAP).*
- *Growth conditions: 28°C (day)/22°C (night); 16h light:8h dark; Light Intensity: 400 uMol, 50% RH, Soil type: Berger BM7.*

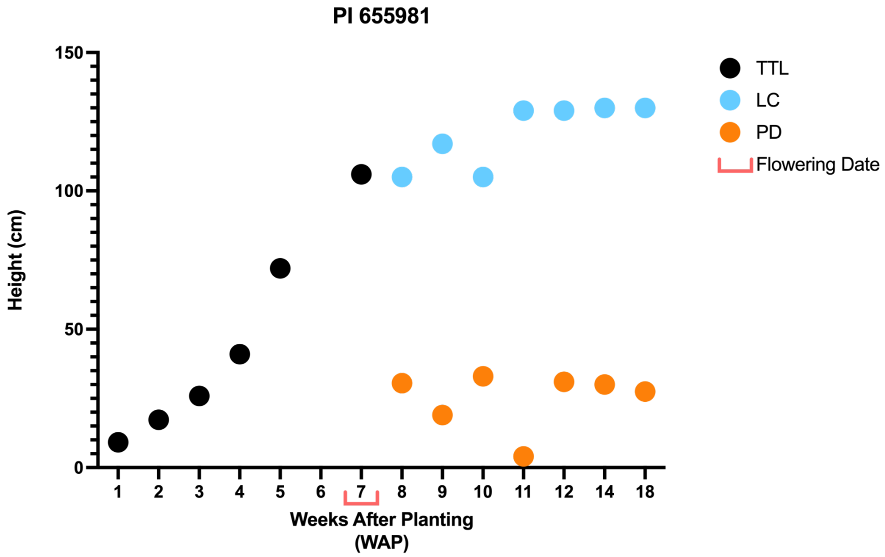


- *The plant was measured up to the last collar (LC) as well as the peduncle to the base of the panicle (PD). Before panicle formation, the height was measured up to the top of the tallest leaf (TTL).*

# PI 656111

**Other name(s):** SC 971
**Species:** Sorghum bicolor subsp. bicolor
**Group:** ICRISAT Reference Collection & Sorghum Association Panel

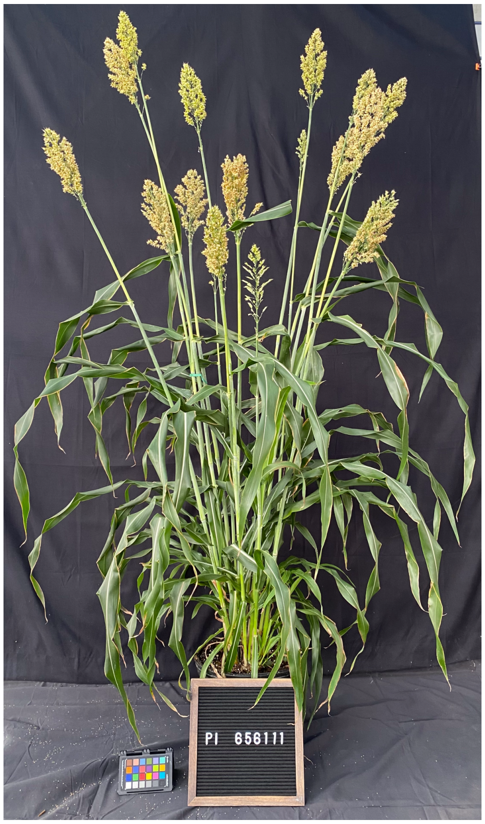


| | |
|---|---|
| **Flowering Days Under Short Days** | 54 days after planting |
| **Seed Color** | Tan |
| **Seedling Color** | Green |
| **Mid-Rib Color** | White |
| **Lodging** | No |
| **Stem diameter (mm) 3rd internode** | 10.6 |
| **Stem diameter (mm) 6th internode** | 7.0 |
| **Main Stem Height: LC; PD (cm)** | 85; 23 |
| **Total Number of Tillers, with panicles** | 17, 16 |
| **Tiller 1 Stem Height: LC; PD (cm)** | 91; 30 |
| **Tiller 2 Stem Height: LC; PD (cm)** | 120; 26 |
| **Tiller 3 Stem Height: LC; PD (cm)** | 85; 18 |

- *Stem diameter: recorded both axes means. if oval-shaped "x, y".*
- *Measurements highlighted in green and photos were recorded 18 weeks after planting (WAP).*
- *Growth conditions: 28°C (day)/22°C (night); 16h light:8h dark; Light Intensity: 400 uMol, 50% RH, Soil type: Berger BM7.*

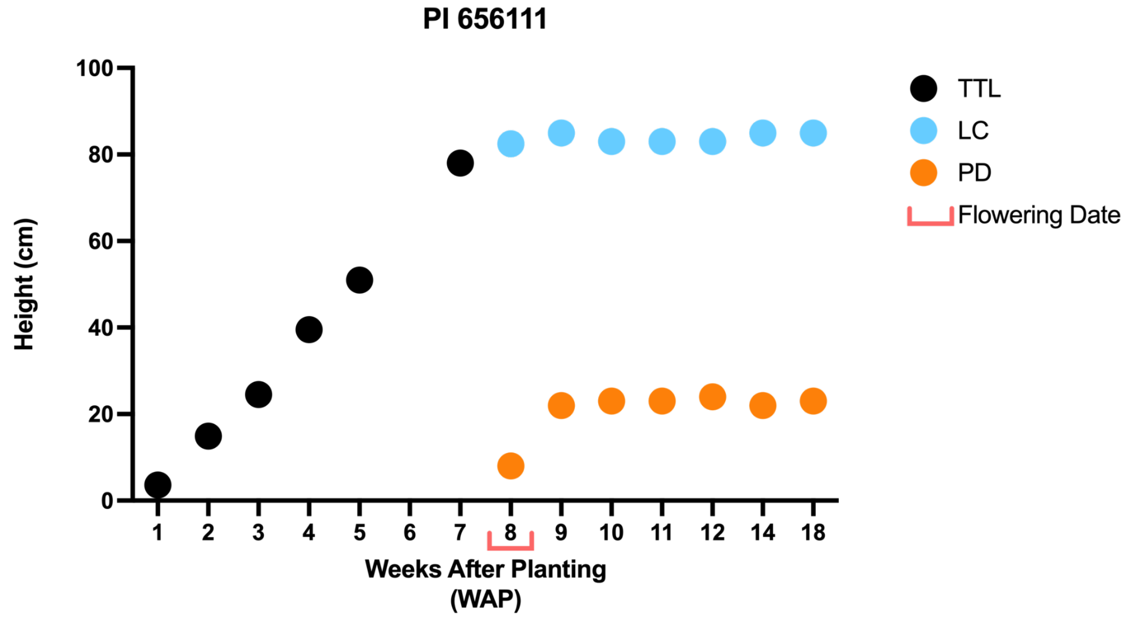


- *The plant was measured up to the last collar (LC) as well as the peduncle to the base of the panicle (PD). Before panicle formation, the height was measured up to the top of the tallest leaf (TTL).*

# PI 660563

**Other name(s):** IS 7173
**Species:** Sorghum bicolor subsp. bicolor
**Group:** ICRISAT Reference Collection

| | |
|---|---|
| Flowering Days Under Short Days | 59 days after planting |
| Seed Color | White |
| Seedling Color | Green |
| Mid-Rib Color | White |
| Lodging | No |
| Stem diameter (mm) 3rd internode | 11.0 |
| Stem diameter (mm) 6th internode | 7.6 |
| Main Stem Height: LC; PD (cm) | 139; 43 |
| Total Number of Tillers, with panicles | 19, 15 |
| Tiller 1 Stem Height: LC; PD (cm) | 171; 44 |
| Tiller 2 Stem Height: LC; PD (cm) | 167; 45 |
| Tiller 3 Stem Height: LC; PD (cm) | 162; 51 |

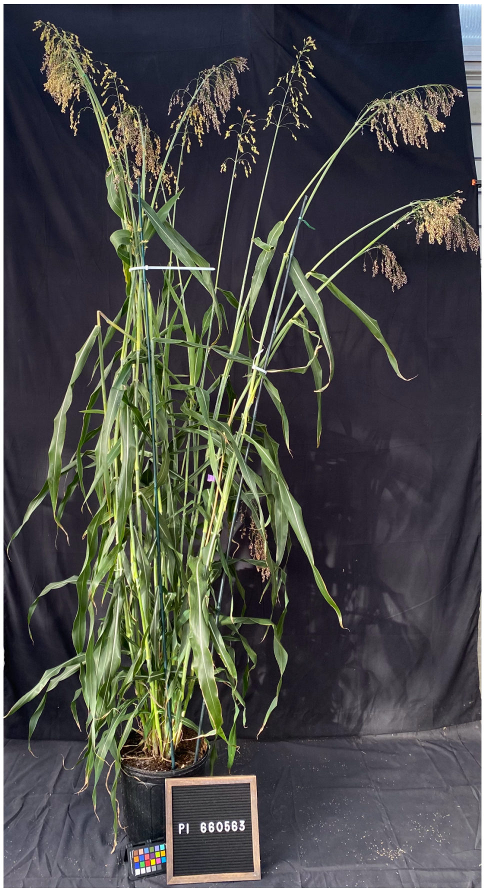


- *Stem diameter: recorded both axes means. if oval-shaped "x, y".*
- *Measurements highlighted in green and photos were recorded 18 weeks after planting (WAP).*
- *Growth conditions: 28°C (day)/22°C (night); 16h light:8h dark; Light Intensity: 400 uMol, 50% RH, Soil type: Berger BM7.*

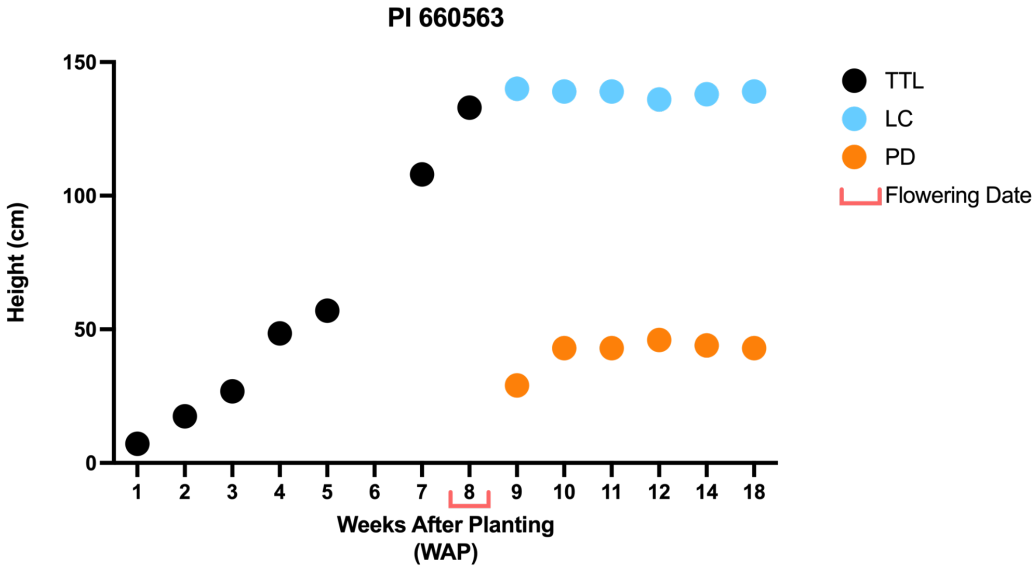


- *The plant was measured up to the last collar (LC) as well as the peduncle to the base of the panicle (PD). Before panicle formation, the height was measured up to the top of the tallest leaf (TTL).*

# PI 656023

**Other name(s):** Segaolane
**Species:** Sorghum bicolor subsp. bicolor
**Group:** ICRISAT Reference Collection & Sorghum Association Panel

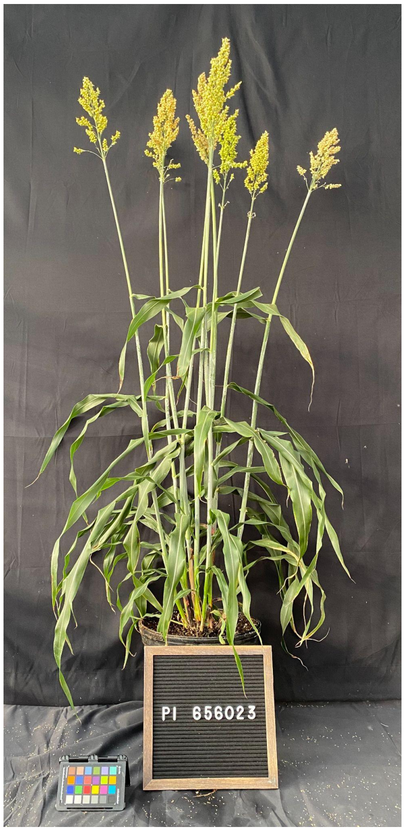


| | |
|---|---|
| **Flowering Days Under Short Days** | 47 days after planting |
| **Seed Color** | Yellow |
| **Seedling Color** | Green |
| **Mid-Rib Color** | White |
| **Lodging** | No |
| **Stem diameter (mm) 3rd internode** | 31.2, 22.9 |
| **Stem diameter (mm) 6th internode** | 20.2, 18.0 |
| **Main Stem Height: LC; PD (cm)** | 64; 31 |
| **Total Number of Tillers, with panicles** | 6, 4 |
| **Tiller 1 Stem Height: LC; PD (cm)** | 77; 32.5 |
| **Tiller 2 Stem Height: LC; PD (cm)** | 63; 2 |
| **Tiller 3 Stem Height: LC; PD (cm)** | 71; 35.5 |

- *Stem diameter: recorded both axes means. if oval-shaped "x, y".*
- *Measurements highlighted in green and photos were recorded 18 weeks after planting (WAP).*
- *Growth conditions: 28°C (day)/22°C (night); 16h light:8h dark; Light Intensity: 400 uMol, 50% RH, Soil type: Berger BM7.*

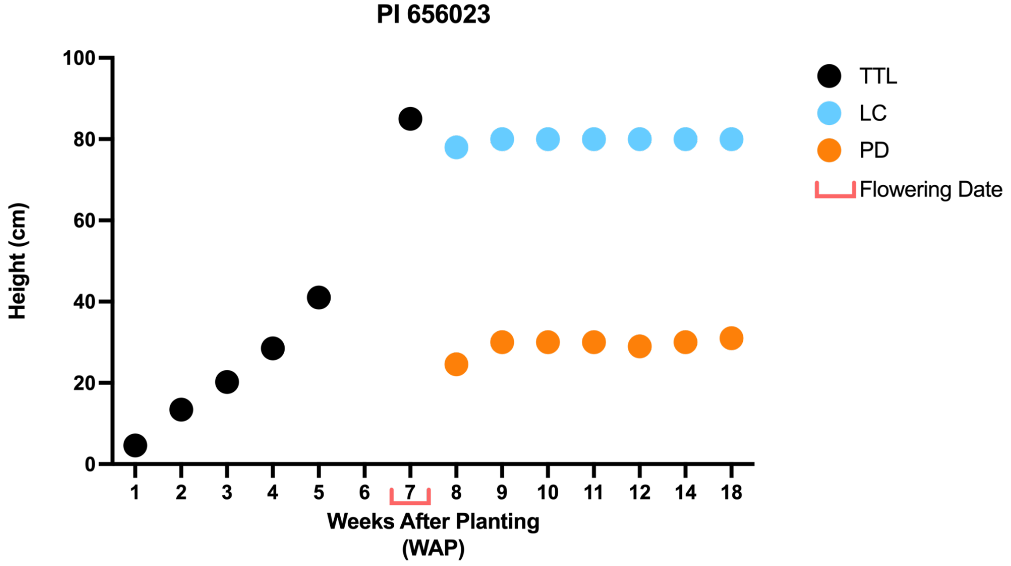


- *The plant was measured up to the last collar (LC) as well as the peduncle to the base of the panicle (PD). Before panicle formation, the height was measured up to the top of the tallest leaf (TTL).*

# PI 276816

**Other name(s):** IS 12646
**Species:** Sorghum bicolor subsp. bicolor
**Group:** ICRISAT Reference Collection

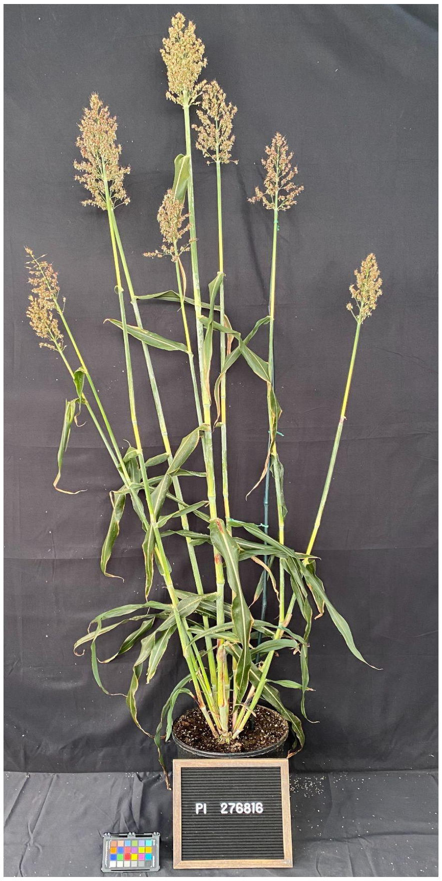


| | |
|---|---|
| **Flowering Days Under Short Days** | 59 days after planting |
| **Seed Color** | Brown-purple |
| **Seedling Color** | Purple |
| **Mid-Rib Color** | White |
| **Lodging** | Yes |
| **Stem diameter (mm) 3rd internode** | 15.5 |
| **Stem diameter (mm) 6th internode** | 11.7 |
| **Main Stem Height: LC; PD (cm)** | 165; 14 |
| **Total Number of Tillers, with panicles** | 8, 8 |
| **Tiller 1 Stem Height: LC; PD (cm)** | 135; 29 |
| **Tiller 2 Stem Height: LC; PD (cm)** | 129; 24 |
| **Tiller 3 Stem Height: LC; PD (cm)** | 112; 25 |

- *Stem diameter: recorded both axes means. if oval-shaped "x, y".*
- *Measurements highlighted in green and photos were recorded 18 weeks after planting (WAP).*
- *Growth conditions: 28°C (day)/22°C (night); 16h light:8h dark; Light Intensity: 400 uMol, 50% RH, Soil type: Berger BM7.*

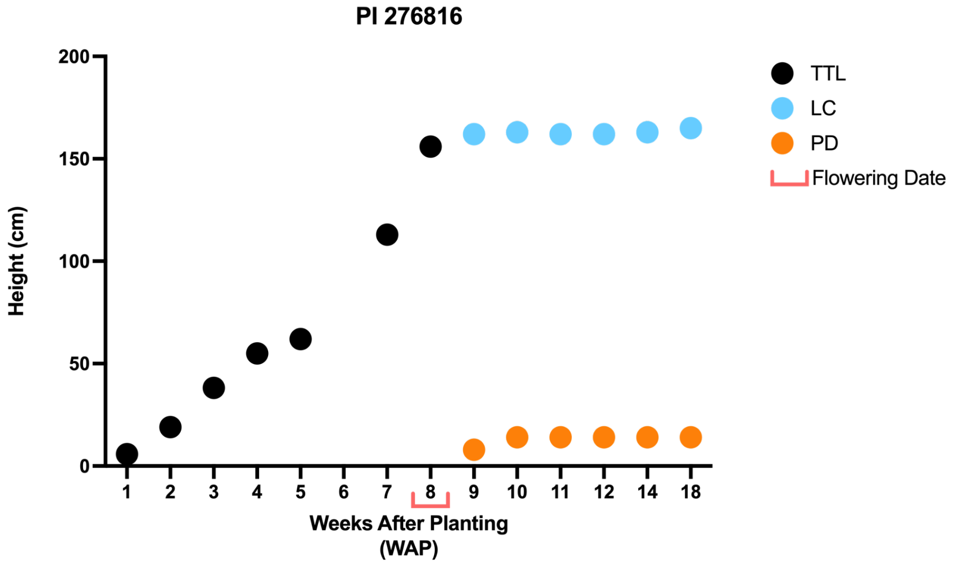


- *The plant was measured up to the last collar (LC) as well as the peduncle to the base of the panicle (PD). Before panicle formation, the height was measured up to the top of the tallest leaf (TTL).*

# PI 154844

**Other name(s):** GRASSL
**Species:** Sorghum bicolor subsp. bicolor
**Group:** ICRISAT Reference Collection & Bioenergy Association Panel

| | |
|---|---|
| Flowering Days Under Short Days | 66 days after planting |
| Seed Color | Red |
| Seedling Color | Purple |
| Mid-Rib Color | Tan |
| Lodging | No |
| Stem diameter (mm) 3rd internode | 17.5, 15.8 |
| Stem diameter (mm) 6th internode | 16.7 |
| Main Stem Height: LC; PD (cm) | 188; 26.5 |
| Total Number of Tillers, with panicles | 9, 7 |
| Tiller 1 Stem Height: LC; PD (cm) | 120; 12 |
| Tiller 2 Stem Height: LC; PD (cm) | 187; 16 |
| Tiller 3 Stem Height: LC; PD (cm) | 175; 21 |

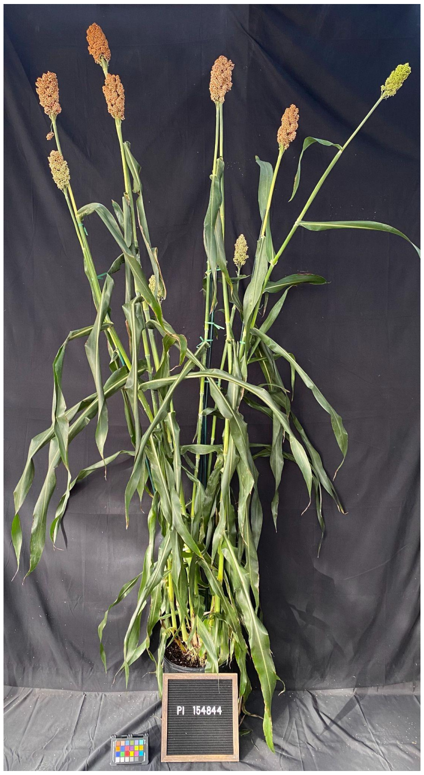


- *Stem diameter: recorded both axes means. if oval-shaped "x, y".*
- *Measurements highlighted in green and photos were recorded 18 weeks after planting (WAP).*
- *Growth conditions: 28°C (day)/22°C (night); 16h light:8h dark; Light Intensity: 400 uMol, 50% RH, Soil type: Berger BM7.*

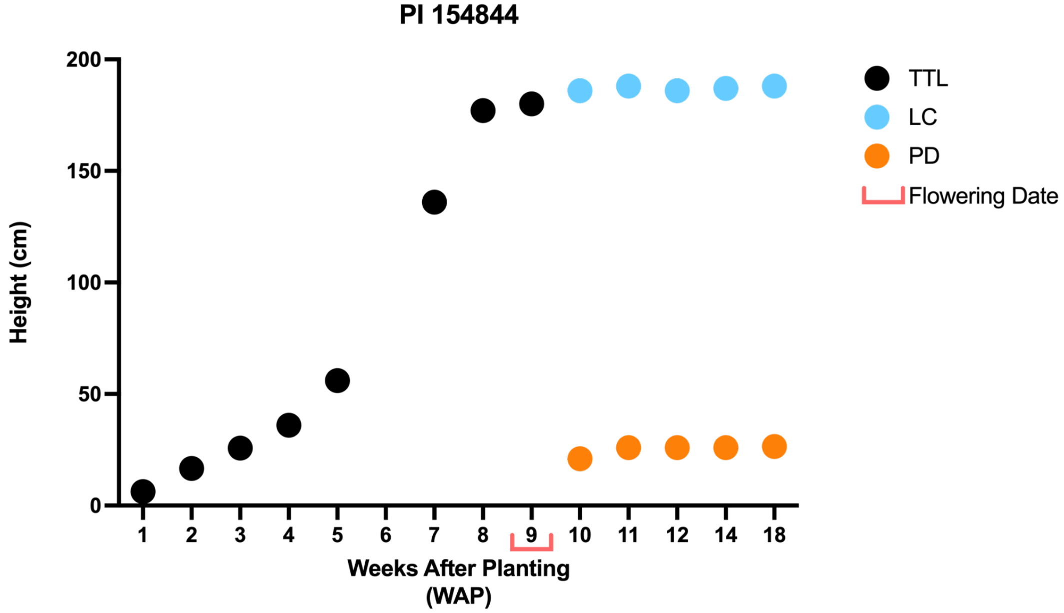


- *The plant was measured up to the last collar (LC) as well as the peduncle to the base of the panicle (PD). Before panicle formation, the height was measured up to the top of the tallest leaf (TTL).*

# PI 276837

**Other name(s):** IS 12661
**Species:** Sorghum bicolor subsp. bicolor
**Group:** ICRISAT Reference Collection, Sorghum Association Panel & Bioenergy Association Panel

| | |
|---|---|
| **Flowering Days Under Short Days** | 59 days after planting |
| **Seed Color** | White |
| **Seedling Color** | Green |
| **Mid-Rib Color** | Tan |
| **Lodging** | No |
| **Stem diameter (mm) 3rd internode** | 11.0 |
| **Stem diameter (mm) 6th internode** | 12.6, 9.2 |
| **Main Stem Height: LC; PD (cm)** | 146; 10.5 |
| **Total Number of Tillers, with panicles** | 12, 9 |
| **Tiller 1 Stem Height: LC; PD (cm)** | 169; 17.5 |
| **Tiller 2 Stem Height: LC; PD (cm)** | 138; 21 |
| **Tiller 3 Stem Height: LC; PD (cm)** | 168; 14 |

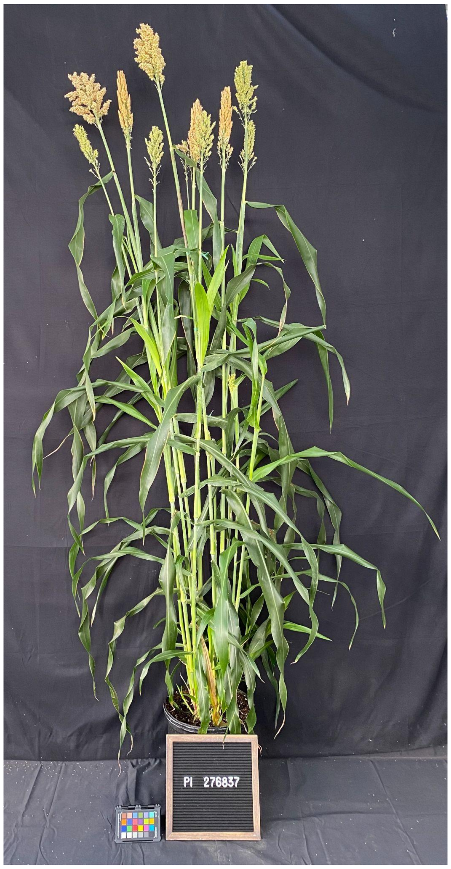


- *Stem diameter: recorded both axes means. if oval-shaped "x, y".*
- *Measurements highlighted in green and photos were recorded 18 weeks after planting (WAP).*
- *Growth conditions: 28°C (day)/22°C (night); 16h light:8h dark; Light Intensity: 400 uMol, 50% RH, Soil type: Berger BM7.*

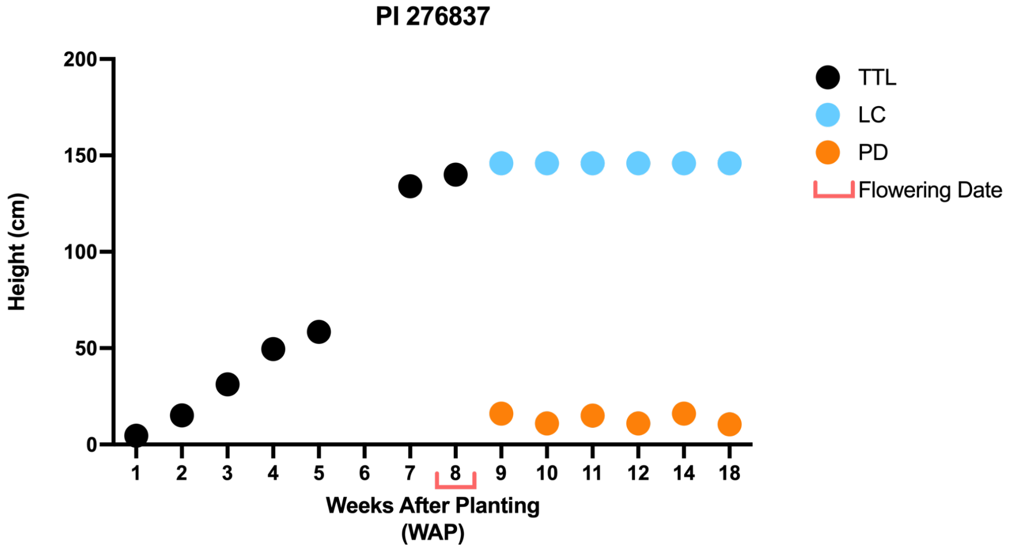


- *The plant was measured up to the last collar (LC) as well as the peduncle to the base of the panicle (PD). Before panicle formation, the height was measured up to the top of the tallest leaf (TTL).*

# PI 565121

**Other name(s):** MACIA
**Species:** Sorghum bicolor subsp. bicolor
**Group:** ICRISAT Reference Collection & Sorghum Association Panel

| | |
|---|---|
| **Flowering Days Under Short Days** | 66 days after planting |
| **Seed Color** | White |
| **Seedling Color** | Green |
| **Mid-Rib Color** | Tan |
| **Lodging** | No |
| **Stem diameter (mm) 3rd internode** | 20.7, 18.4 |
| **Stem diameter (mm) 6th internode** | 19.2 |
| **Main Stem Height: LC; PD (cm)** | 95 |
| **Total Number of Tillers, with panicles** | 6, 6 |
| **Tiller 1 Stem Height: LC; PD (cm)** | 67 |
| **Tiller 2 Stem Height: LC; PD (cm)** | 95; 15 |
| **Tiller 3 Stem Height: LC; PD (cm)** | 90; 9 |

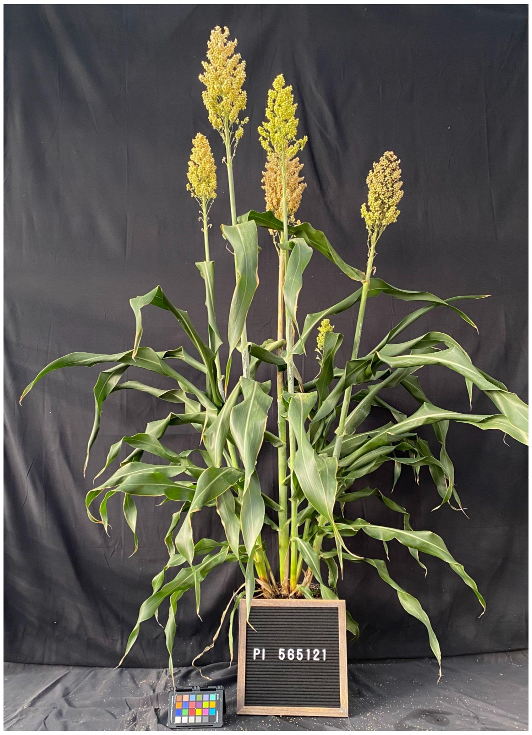


- *Stem diameter: recorded both axes means. if oval-shaped "x, y".*
- *Measurements highlighted in green and photos were recorded 18 weeks after planting (WAP).*
- *Growth conditions: 28°C (day)/22°C (night); 16h light:8h dark; Light Intensity: 400 uMol, 50% RH, Soil type: Berger BM7.*

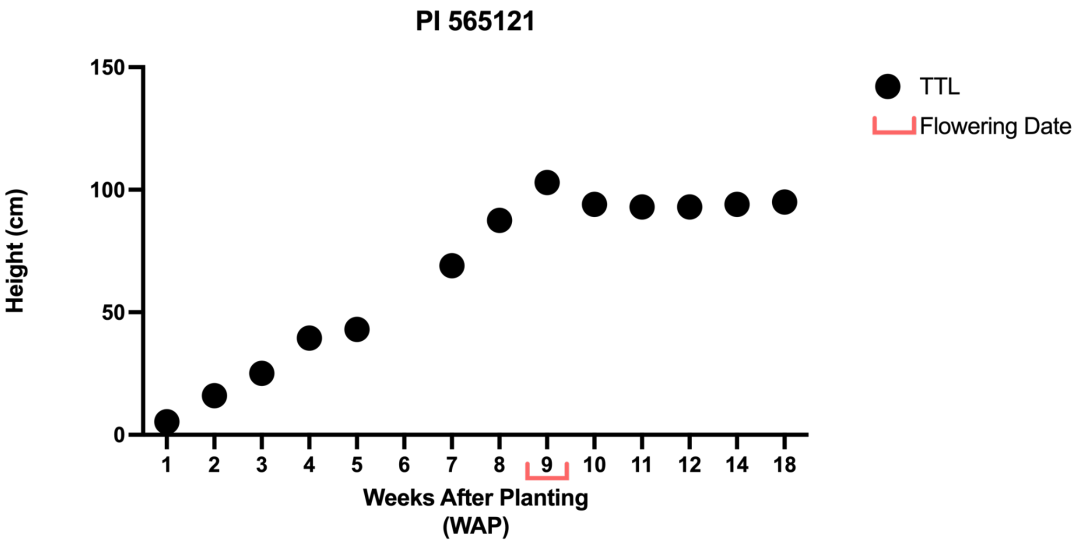


- *The plant was measured up to the last collar (LC) as well as the peduncle to the base of the panicle (PD). Before panicle formation, the height was measured up to the top of the tallest leaf (TTL).*

# PI 576434

**Other name(s):** SC 1103, Ex-Mubi
**Species:** Sorghum bicolor subsp. bicolor
**Group:** ICRISAT Reference Collection & Sorghum Association Panel

| | |
|---|---|
| **Flowering Days Under Short Days** | 73 days after planting |
| **Seed Color** | White |
| **Seedling Color** | Green |
| **Mid-Rib Color** | White |
| **Lodging** | No |
| **Stem diameter (mm) 3rd internode** | 23.4, 21.3 |
| **Stem diameter (mm) 6th internode** | 21.1, 20.0 |
| **Main Stem Height: LC; PD (cm)** | 142; 13 |
| **Total Number of Tillers, with panicles** | 5, 5 |
| **Tiller 1 Stem Height: LC; PD (cm)** | 134; 13 |
| **Tiller 2 Stem Height: LC; PD (cm)** | 132; 12 |
| **Tiller 3 Stem Height: LC; PD (cm)** | 105; 10 |

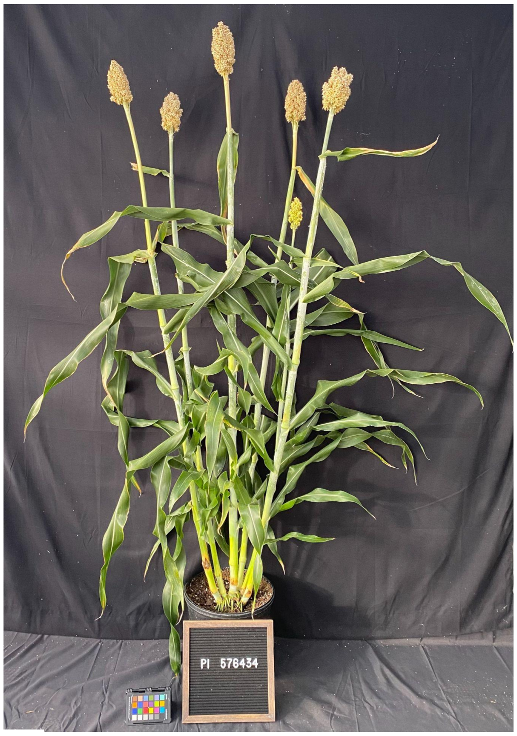


- *Stem diameter: recorded both axes means. if oval-shaped "x, y".*
- *Measurements highlighted in green and photos were recorded 18 weeks after planting (WAP).*
- *Growth conditions: 28°C (day)/22°C (night); 16h light:8h dark; Light Intensity: 400 uMol, 50% RH, Soil type: Berger BM7.*

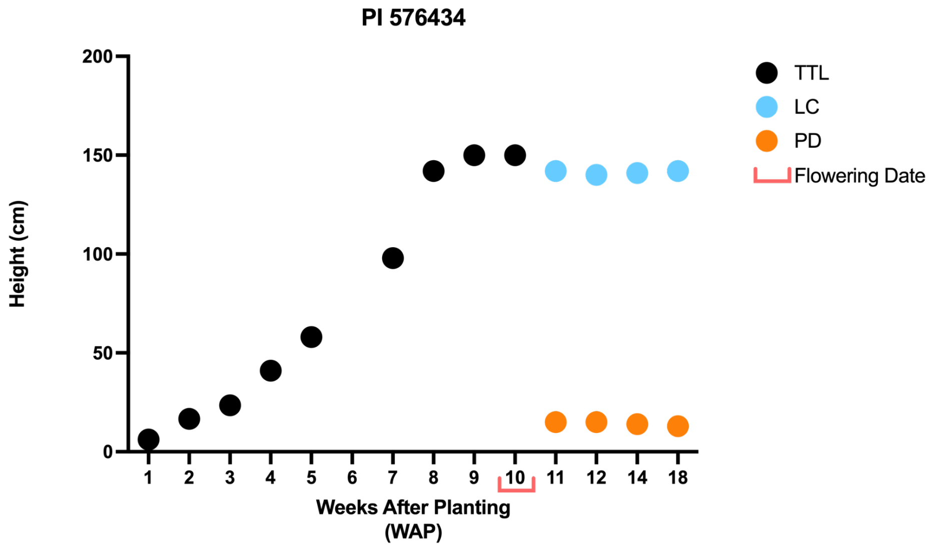


- *The plant was measured up to the last collar (LC) as well as the peduncle to the base of the panicle (PD). Before panicle formation, the height was measured up to the top of the tallest leaf (TTL).*

# PI 656015

**Other name(s):** Ajabsido
**Species:** Sorghum bicolor subsp. bicolor
**Group:** ICRISAT Reference Collection, Sorghum Association Panel & Bioenergy Association Panel

| | |
|---|---|
| **Flowering Days Under Short Days** | 47 days after planting |
| **Seed Color** | White |
| **Seedling Color** | Green |
| **Mid-Rib Color** | White |
| **Lodging** | No |
| **Stem diameter (mm) 3rd internode** | 9.4 |
| **Stem diameter (mm) 6th internode** | NA |
| **Main Stem Height: LC; PD (cm)** | 77; 15.5 |
| **Total Number of Tillers, with panicles** | 17, 10 |
| **Tiller 1 Stem Height: LC; PD (cm)** | 78; 11 |
| **Tiller 2 Stem Height: LC; PD (cm)** | 87; 11 |
| **Tiller 3 Stem Height: LC; PD (cm)** | 62; 13 |

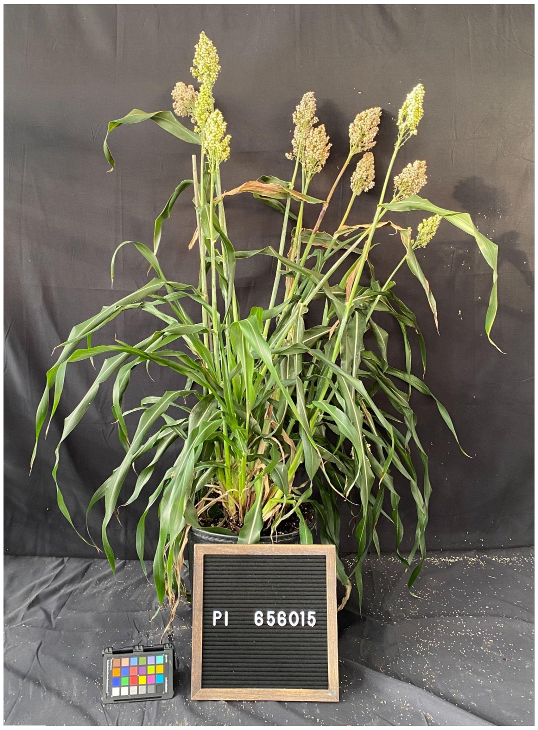


- *Stem diameter: recorded both axes means. if oval-shaped "x, y".*
- *Measurements highlighted in green and photos were recorded 18 weeks after planting (WAP).*
- *Growth conditions: 28°C (day)/22°C (night); 16h light:8h dark; Light Intensity: 400 uMol, 50% RH, Soil type: Berger BM7.*

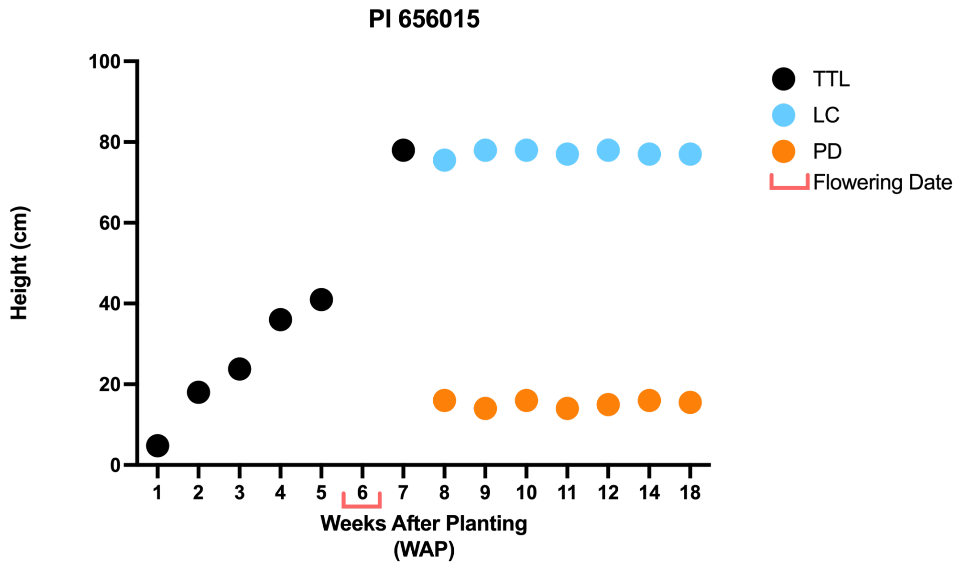


- *The plant was measured up to the last collar (LC) as well as the peduncle to the base of the panicle (PD). Before panicle formation, the height was measured up to the top of the tallest leaf (TTL).*

# PI 656050

**Other name(s):** Mota Maradi
**Species:** Sorghum bicolor subsp. bicolor
**Group:** ICRISAT Reference Collection & Sorghum Association Panel

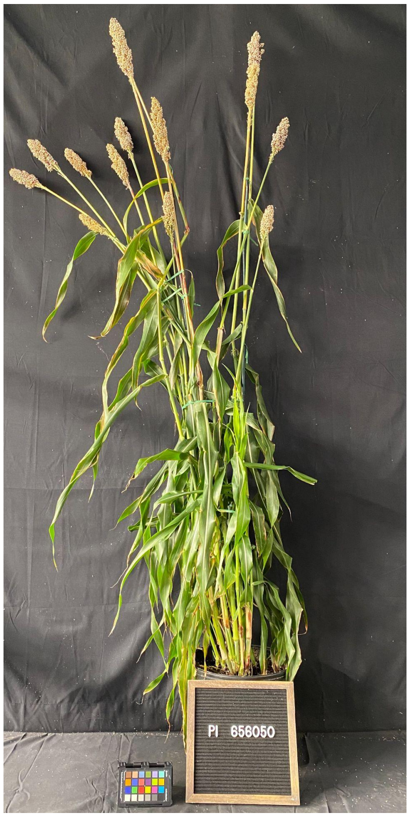


| | |
|---|---|
| **Flowering Days Under Short Days** | 52 days after planting |
| **Seed Color** | White |
| **Seedling Color** | Purple |
| **Mid-Rib Color** | White |
| **Lodging** | No |
| **Stem diameter (mm) 3rd internode** | 10.5 |
| **Stem diameter (mm) 6th internode** | NA |
| **Main Stem Height: LC; PD (cm)** | 112; 27.5 |
| **Total Number of Tillers, with panicles** | 14, 13 |
| **Tiller 1 Stem Height: LC; PD (cm)** | 106; 9.5 |
| **Tiller 2 Stem Height: LC; PD (cm)** | 109; 12 |
| **Tiller 3 Stem Height: LC; PD (cm)** | 113; 33 |

- *Stem diameter: recorded both axes means. if oval-shaped "x, y".*
- *Measurements highlighted in green and photos were recorded 18 weeks after planting (WAP).*
- *Growth conditions: 28°C (day)/22°C (night); 16h light:8h dark; Light Intensity: 400 uMol, 50% RH, Soil type: Berger BM7.*

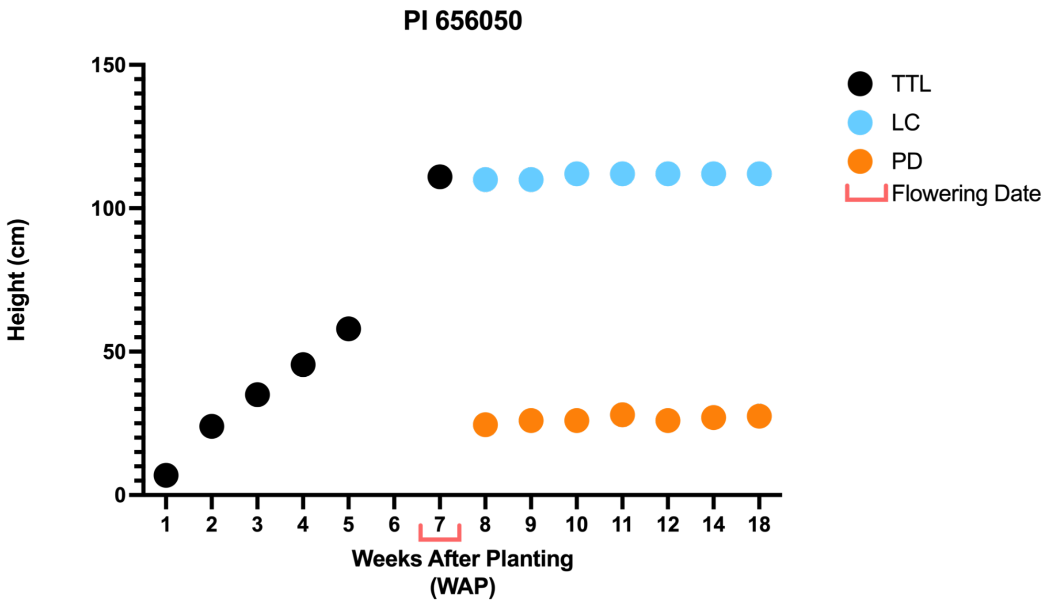


- *The plant was measured up to the last collar (LC) as well as the peduncle to the base of the panicle (PD). Before panicle formation, the height was measured up to the top of the tallest leaf (TTL).*

# PI 656057

**Other name(s):** P898012
**Species:** Sorghum bicolor subsp. bicolor
**Group:** ICRISAT Reference Collection & Sorghum Association Panel

| | |
|---|---|
| **Flowering Days Under Short Days** | 52 days after planting |
| **Seed Color** | White |
| **Seedling Color** | Purple |
| **Mid-Rib Color** | Tan |
| **Lodging** | No |
| **Stem diameter (mm) 3rd internode** | 10.2 |
| **Stem diameter (mm) 6th internode** | 6.3 |
| **Main Stem Height: LC; PD (cm)** | 83; 20 |
| **Total Number of Tillers, with panicles** | 11, 9 |
| **Tiller 1 Stem Height: LC; PD (cm)** | 83; 23 |
| **Tiller 2 Stem Height: LC; PD (cm)** | 76; 20.5 |
| **Tiller 3 Stem Height: LC; PD (cm)** | 84 |

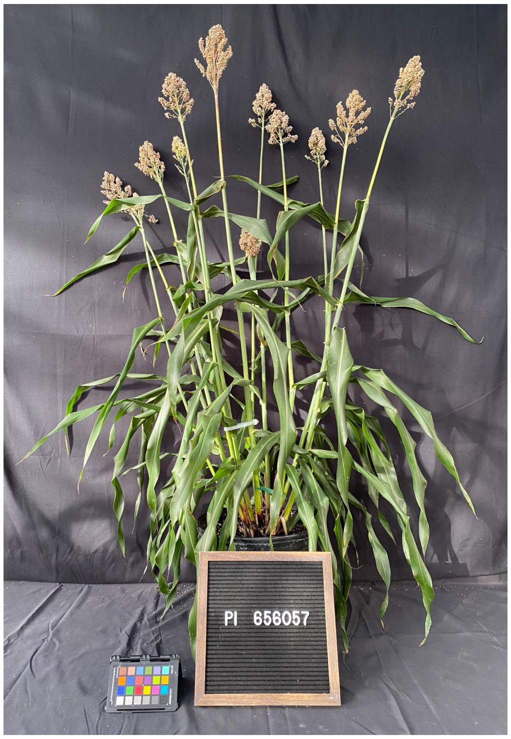


- *Stem diameter: recorded both axes means. if oval-shaped "x, y".*
- *Measurements highlighted in green and photos were recorded 18 weeks after planting (WAP).*
- *Growth conditions: 28°C (day)/22°C (night); 16h light:8h dark; Light Intensity: 400 uMol, 50% RH, Soil type: Berger BM7.*

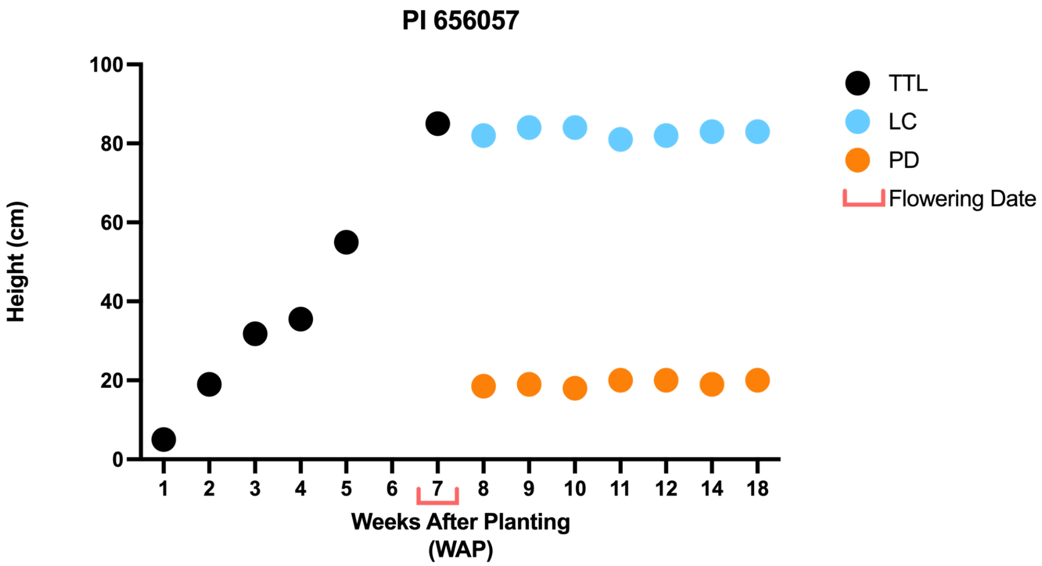


- *The plant was measured up to the last collar (LC) as well as the peduncle to the base of the panicle (PD). Before panicle formation, the height was measured up to the top of the tallest leaf (TTL).*

# PI 597980

**Other name(s):** SC 1345, CSM-90, SAP-173
**Species:** Sorghum bicolor subsp. bicolor
**Group:** ICRISAT Reference Collection & Sorghum Association Panel

| | |
|---|---|
| **Flowering Days Under Short Days** | 47 days after planting |
| **Seed Color** | White |
| **Seedling Color** | Purple |
| **Mid-Rib Color** | White |
| **Lodging** | Tan |
| **Stem diameter (mm) 3rd internode** | 13.2, 11.0 |
| **Stem diameter (mm) 6th internode** | NA |
| **Main Stem Height: LC; PD (cm)** | 71 |
| **Total Number of Tillers, with panicles** | 13, 12 |
| **Tiller 1 Stem Height: LC; PD (cm)** | 68 |
| **Tiller 2 Stem Height: LC; PD (cm)** | 82; 1 |
| **Tiller 3 Stem Height: LC; PD (cm)** | 79 |

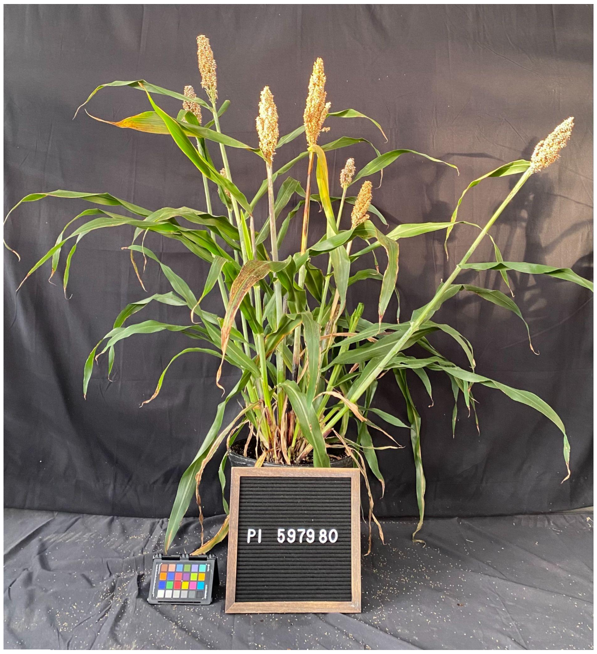


- *Stem diameter: recorded both axes means. if oval-shaped "x, y".*
- *Measurements highlighted in green and photos were recorded 18 weeks after planting (WAP).*
- *Growth conditions: 28°C (day)/22°C (night); 16h light:8h dark; Light Intensity: 400 uMol, 50% RH, Soil type: Berger BM7.*

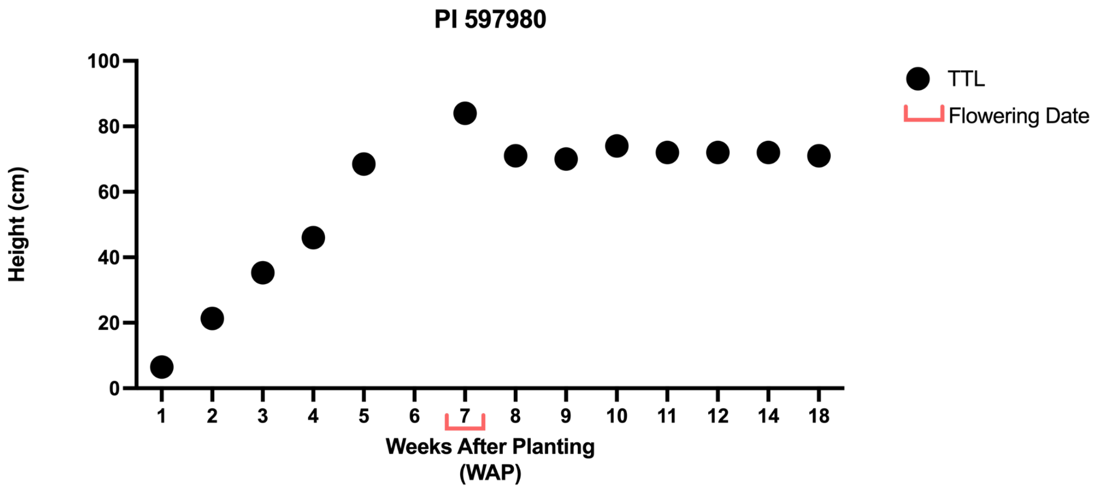


- *The plant was measured up to the last collar (LC) as well as the peduncle to the base of the panicle (PD). Before panicle formation, the height was measured up to the top of the tallest leaf (TTL).*

# PI 656027

**Other name(s):** SRN39
**Species:** Sorghum bicolor subsp. bicolor
**Group:** ICRISAT Reference Collection & Sorghum Association Panel

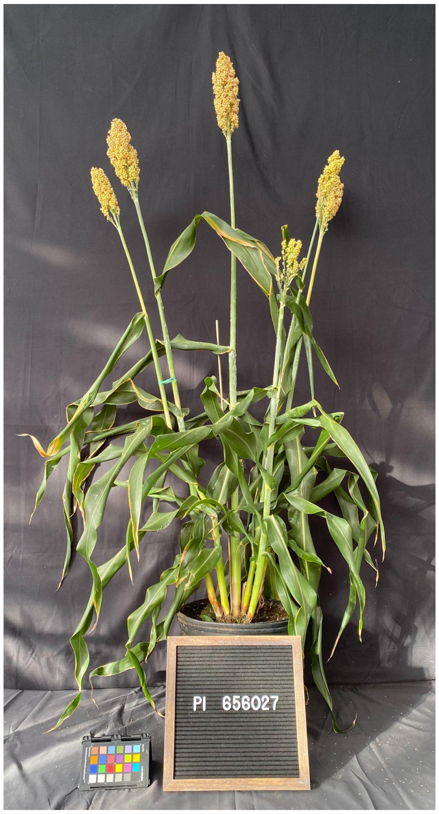


| | |
|---|---|
| **Flowering Days Under Short Days** | 60 days after planting |
| **Seed Color** | Tan |
| **Seedling Color** | Green |
| **Mid-Rib Color** | Tan |
| **Lodging** | No |
| **Stem diameter (mm) 3rd internode** | 16.3 |
| **Stem diameter (mm) 6th internode** | 15.5 |
| **Main Stem Height: LC; PD (cm)** | 95; 23 |
| **Total Number of Tillers, with panicles** | 5, 4 |
| **Tiller 1 Stem Height: LC; PD (cm)** | 80; 23 |
| **Tiller 2 Stem Height: LC; PD (cm)** | 79; 19 |
| **Tiller 3 Stem Height: LC; PD (cm)** | 72 |

- *Stem diameter: recorded both axes means. if oval-shaped "x, y".*
- *Measurements highlighted in green and photos were recorded 18 weeks after planting (WAP).*
- *Growth conditions: 28°C (day)/22°C (night); 16h light:8h dark; Light Intensity: 400 uMol, 50% RH, Soil type: Berger BM7.*

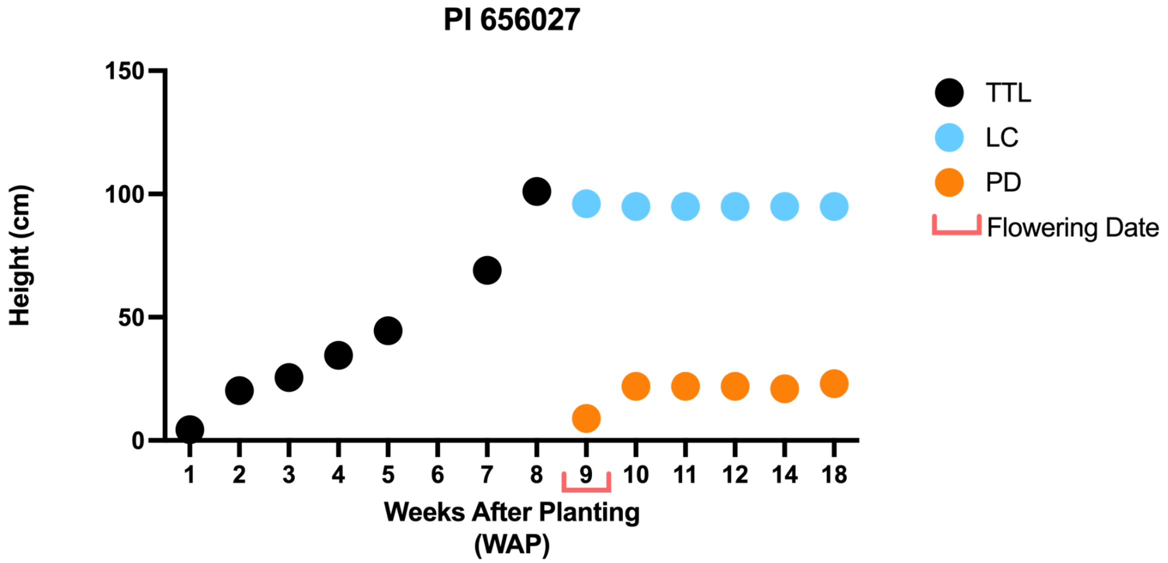


- *The plant was measured up to the last collar (LC) as well as the peduncle to the base of the panicle (PD). Before panicle formation, the height was measured up to the top of the tallest leaf (TTL).*

# PI 660565

**Other name(s):** Kamjin, IS 3620
**Species:** Sorghum bicolor subsp. bicolor
**Group:** ICRISAT Reference Collection

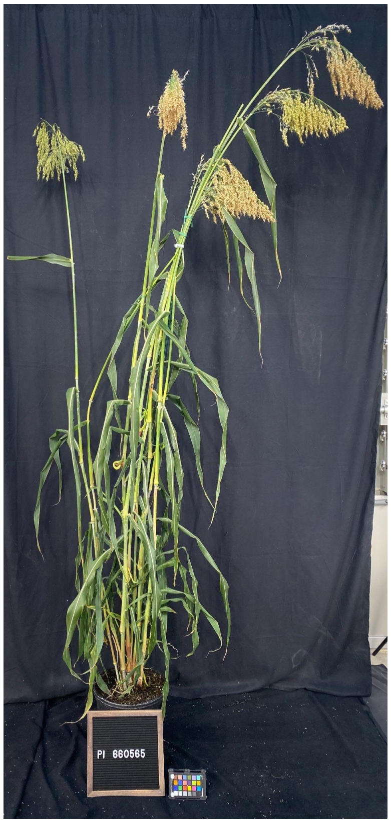


| | |
|---|---|
| Flowering Days Under Short Days | 77 days after planting |
| Seed Color | Tan |
| Seedling Color | Purple |
| Mid-Rib Color | White |
| Lodging | No |
| Stem diameter (mm) 3rd internode | 11.9 |
| Stem diameter (mm) 6th internode | 11.8 |
| Main Stem Height: LC; PD (cm) | 183; 19 |
| Total Number of Tillers, with panicles | 8, 7 |
| Tiller 1 Stem Height: LC; PD (cm) | 179; 31 |
| Tiller 2 Stem Height: LC; PD (cm) | 149; 31 |
| Tiller 3 Stem Height: LC; PD (cm) | 158; 33 |

- *Stem diameter: recorded both axes means. if oval-shaped "x, y".*
- *Measurements highlighted in green and photos were recorded 18 weeks after planting (WAP).*
- *Growth conditions: 28°C (day)/22°C (night); 16h light:8h dark; Light Intensity: 400 uMol, 50% RH, Soil type: Berger BM7.*

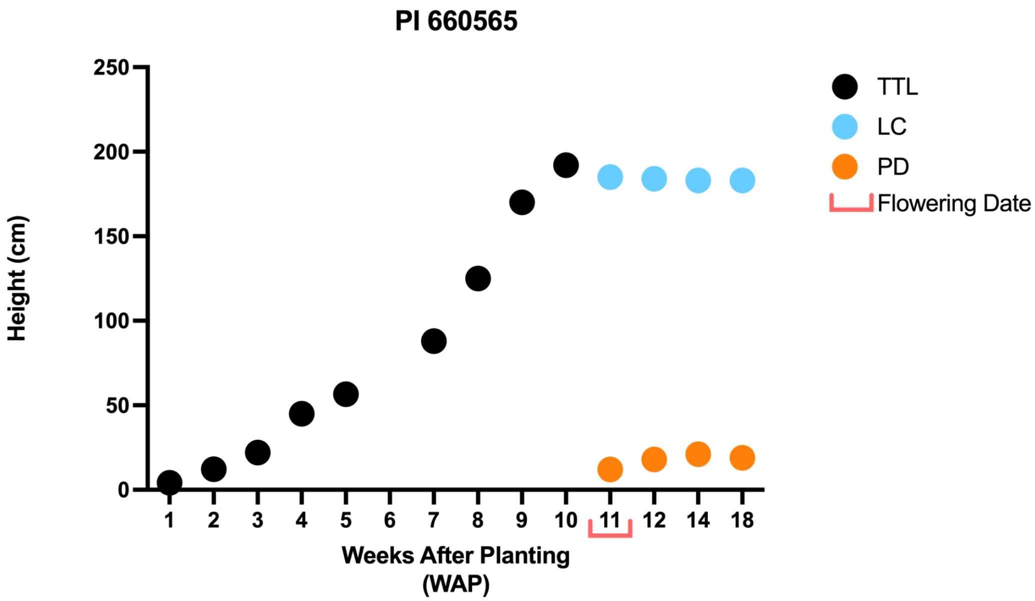


- *The plant was measured up to the last collar (LC) as well as the peduncle to the base of the panicle (PD). Before panicle formation, the height was measured up to the top of the tallest leaf (TTL).*

# PI 180348

**Other name(s):** Juar, IS 12876
**Species:** Sorghum bicolor subsp. bicolor
**Group:** ICRISAT Reference Collection & Bioenergy Association Panel

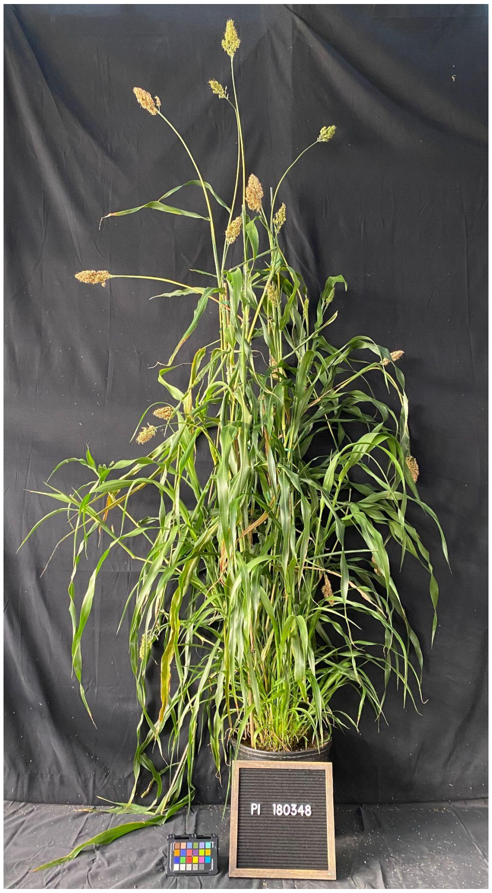


| | |
|---|---|
| **Flowering Days Under Short Days** | 47 days after planting |
| **Seed Color** | NA |
| **Seedling Color** | Green |
| **Mid-Rib Color** | Tan |
| **Lodging** | Yes |
| **Stem diameter (mm) 3rd internode** | 7.3 |
| **Stem diameter (mm) 6th internode** | 5.3 |
| **Main Stem Height: LC; PD (cm)** | 139; 25.5 |
| **Total Number of Tillers, with panicles** | 35, 17 |
| **Tiller 1 Stem Height: LC; PD (cm)** | 166; 42.5 |
| **Tiller 2 Stem Height: LC; PD (cm)** | 164; 33 |
| **Tiller 3 Stem Height: LC; PD (cm)** | 153; 38 |

- *Stem diameter: recorded both axes means. if oval-shaped "x, y".*
- *Measurements highlighted in green and photos were recorded 18 weeks after planting (WAP).*
- *Growth conditions: 28°C (day)/22°C (night); 16h light:8h dark; Light Intensity: 400 uMol, 50% RH, Soil type: Berger BM7.*

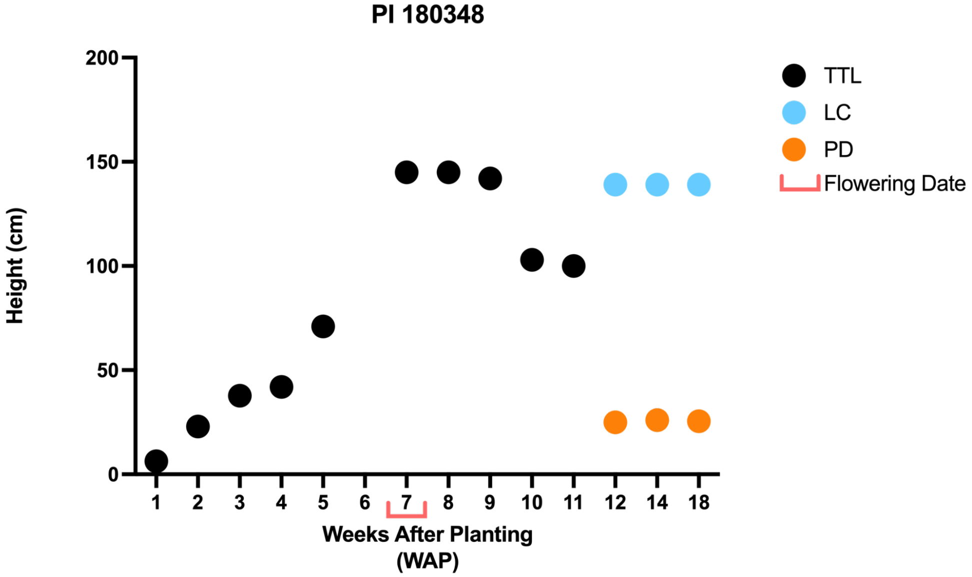


- *The plant was measured up to the last collar (LC) as well as the peduncle to the base of the panicle (PD). Before panicle formation, the height was measured up to the top of the tallest leaf (TTL).*

## PI 329501

**Other name(s):** IS 11257
**Species:** Sorghum bicolor subsp. bicolor
**Group:** ICRISAT Reference Collection, Geo-Reference Collection & Bioenergy Association Panel

| | |
|---|---|
| **Flowering Days Under Short Days** | 81 days after planting |
| **Seed Color** | Red |
| **Seedling Color** | Green |
| **Mid-Rib Color** | White |
| **Lodging** | No |
| **Stem diameter (mm) 3rd internode** | 21.4 |
| **Stem diameter (mm) 6th internode** | 22.0 |
| **Main Stem Height: LC; PD (cm)** | 224 |
| **Total Number of Tillers, with panicles** | 6, 4 |
| **Tiller 1 Stem Height: LC; PD (cm)** | 202; 19 |
| **Tiller 2 Stem Height: LC; PD (cm)** | 207; 11 |
| **Tiller 3 Stem Height: LC; PD (cm)** | 173; 27 |

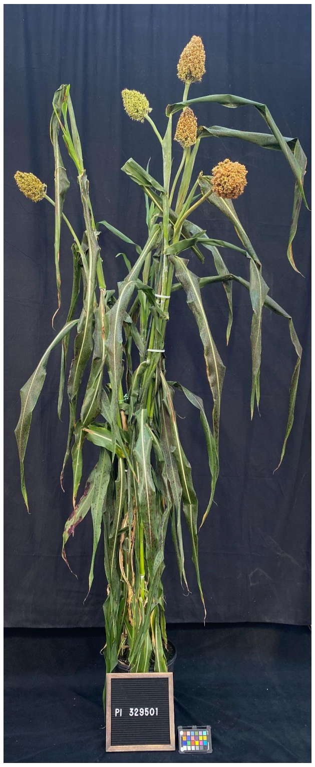


- *Stem diameter: recorded both axes means. if oval-shaped "x, y".*
- *Measurements highlighted in green and photos were recorded 18 weeks after planting (WAP).*
- *Growth conditions: 28°C (day)/22°C (night); 16h light:8h dark; Light Intensity: 400 uMol, 50% RH, Soil type: Berger BM7.*

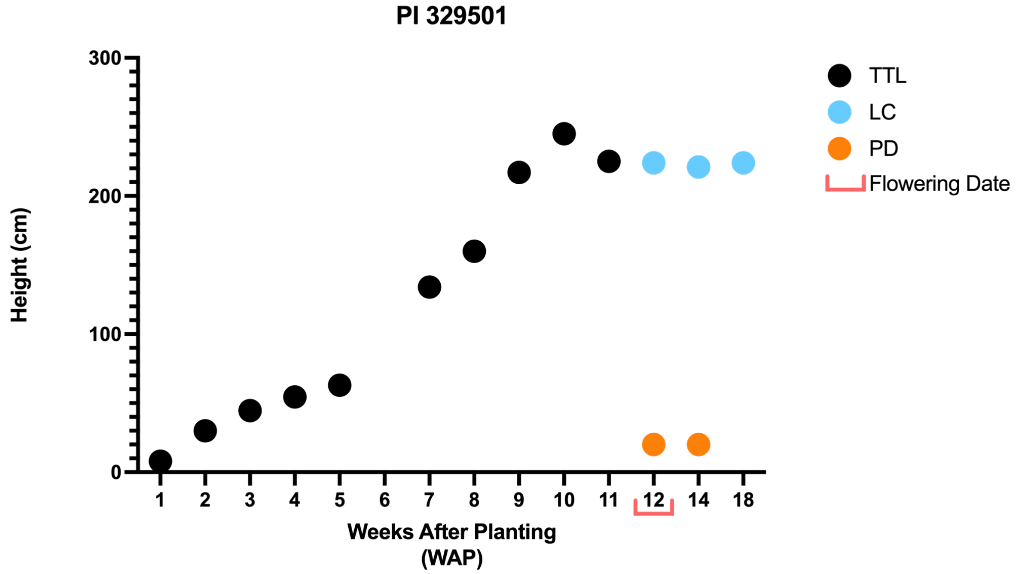


- *The plant was measured up to the last collar (LC) as well as the peduncle to the base of the panicle (PD). Before panicle formation, the height was measured up to the top of the tallest leaf (TTL).*

# PI 156178

**Other name(s):** MN 2014
**Species:** Sorghum bicolor subsp. bicolor
**Group:** ICRISAT Reference Collection & Bioenergy Association Panel

| | |
|---|---|
| **Flowering Days Under Short Days** | 77 days after planting |
| **Seed Color** | Red |
| **Seedling Color** | Green |
| **Mid-Rib Color** | White |
| **Lodging** | No |
| **Stem diameter (mm) 3rd internode** | 13.2 |
| **Stem diameter (mm) 6th internode** | 8.8 |
| **Main Stem Height: LC; PD (cm)** | 159; 14 |
| **Total Number of Tillers, with panicles** | 11, 11 |
| **Tiller 1 Stem Height: LC; PD (cm)** | 150; 22.5 |
| **Tiller 2 Stem Height: LC; PD (cm)** | 156; 14.5 |
| **Tiller 3 Stem Height: LC; PD (cm)** | 122; 15 |

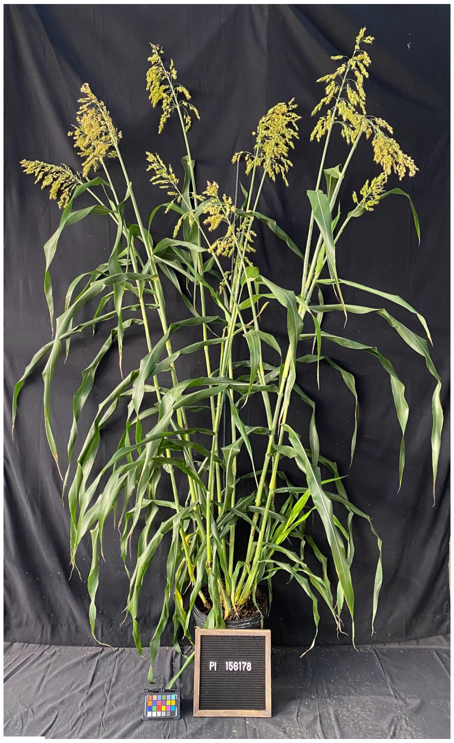


- *Stem diameter: recorded both axes means. if oval-shaped "x, y".*
- *Measurements highlighted in green and photos were recorded 18 weeks after planting (WAP).*
- *Growth conditions: 28°C (day)/22°C (night); 16h light:8h dark; Light Intensity: 400 uMol, 50% RH, Soil type: Berger BM7.*

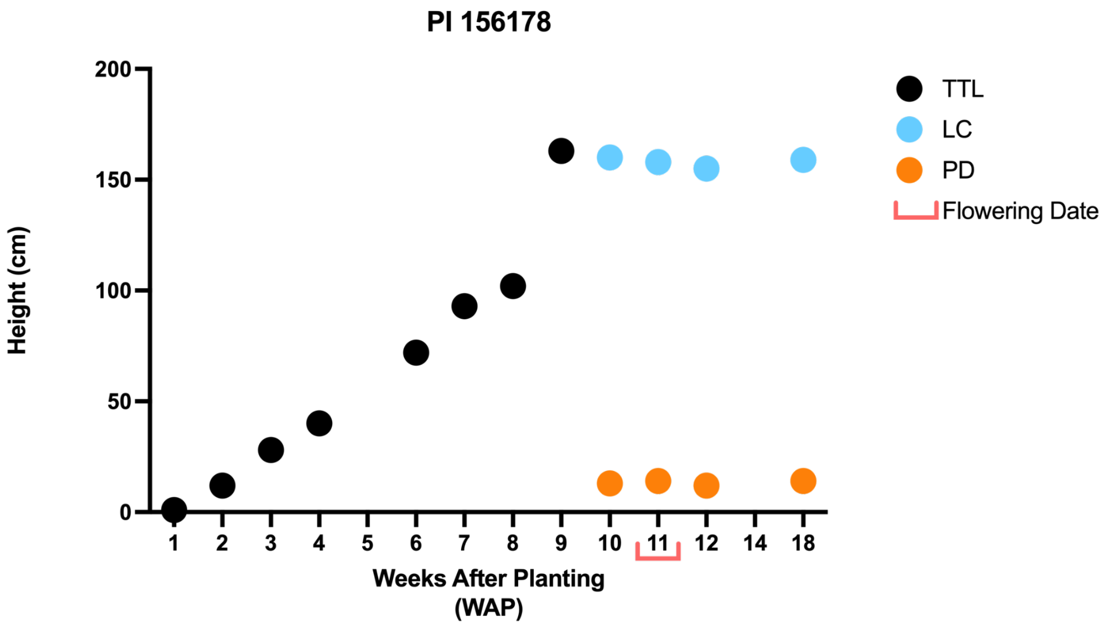


- *The plant was measured up to the last collar (LC) as well as the peduncle to the base of the panicle (PD). Before panicle formation, the height was measured up to the top of the tallest leaf (TTL).*

## PI 569459

**Other name(s):** LUWAIAH, IS 19523
**Species:** Sorghum bicolor subsp. bicolor
**Group:** ICRISAT Reference Collection & Bioenergy Association Panel

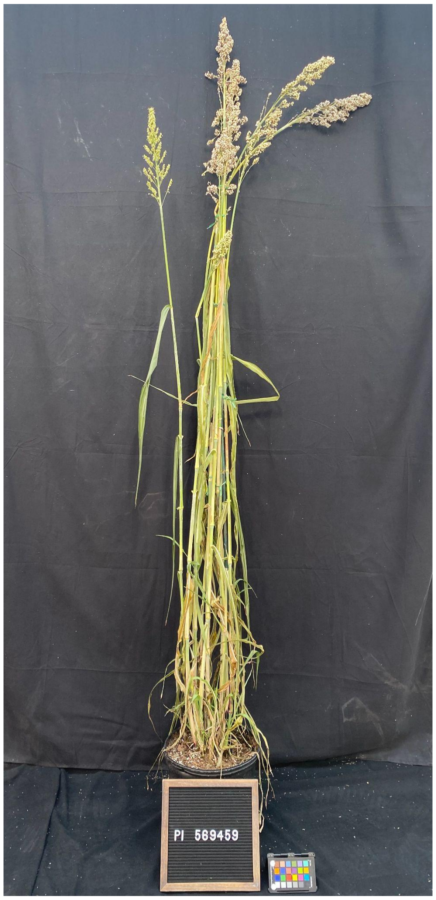


| | |
|---|---|
| **Flowering Days Under Short Days** | 52 days after planting |
| **Seed Color** | White |
| **Seedling Color** | Green |
| **Mid-Rib Color** | White |
| **Lodging** | Yes |
| **Stem diameter (mm) 3rd internode** | 9.9 |
| **Stem diameter (mm) 6th internode** | 6.9 |
| **Main Stem Height: LC; PD (cm)** | 149; 41 |
| **Total Number of Tillers, with panicles** | 13, 11 |
| **Tiller 1 Stem Height: LC; PD (cm)** | 149; 39 |
| **Tiller 2 Stem Height: LC; PD (cm)** | 129; 17.5 |
| **Tiller 3 Stem Height: LC; PD (cm)** | 127; 39.5 |

- *Stem diameter: recorded both axes means. if oval-shaped "x, y".*
- *Measurements highlighted in green and photos were recorded 18 weeks after planting (WAP).*
- *Growth conditions: 28°C (day)/22°C (night); 16h light:8h dark; Light Intensity: 400 uMol, 50% RH, Soil type: Berger BM7.*

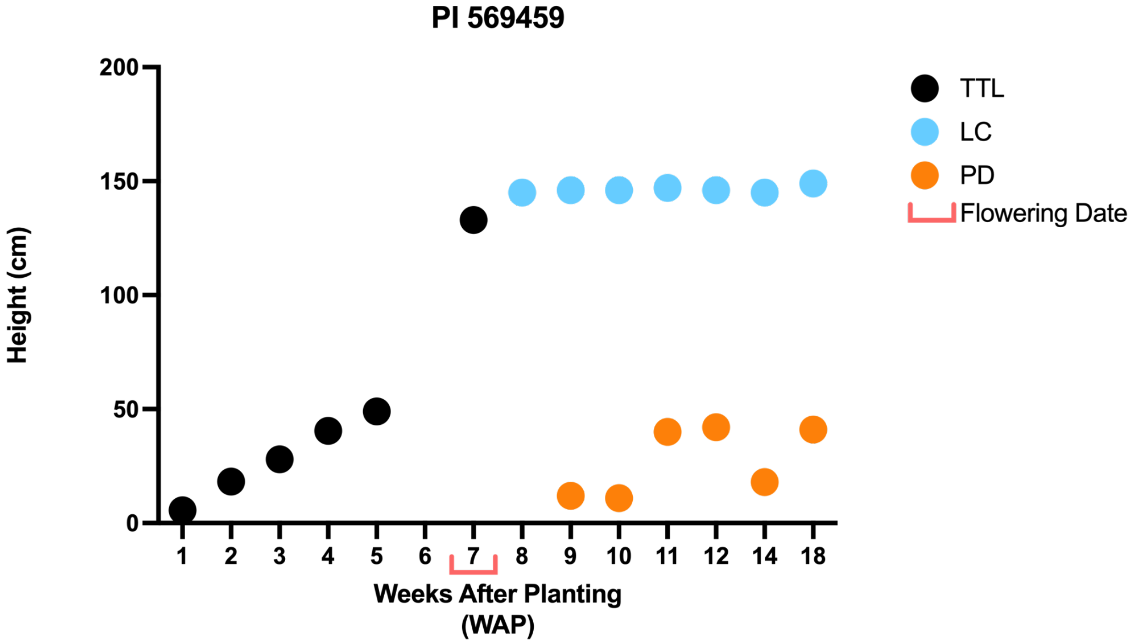


- *The plant was measured up to the last collar (LC) as well as the peduncle to the base of the panicle (PD). Before panicle formation, the height was measured up to the top of the tallest leaf (TTL).*

# PI 570071

**Other name(s):** IS 22849
**Species:** Sorghum bicolor subsp. bicolor
**Group:** ICRISAT Reference Collection & Bioenergy Association Panel

| | |
|---|---|
| Flowering Days Under Short Days | 54 days after planting |
| Seed Color | Red |
| Seedling Color | Green |
| Mid-Rib Color | White |
| Lodging | Yes |
| Stem diameter (mm) 3rd internode | 12.8 |
| Stem diameter (mm) 6th internode | 7.6 |
| Main Stem Height: LC; PD (cm) | 158; 27 |
| Total Number of Tillers, with panicles | 8, 4 |
| Tiller 1 Stem Height: LC; PD (cm) | 122; 28 |
| Tiller 2 Stem Height: LC; PD (cm) | 140; 33 |
| Tiller 3 Stem Height: LC; PD (cm) | 138; 28.5 |

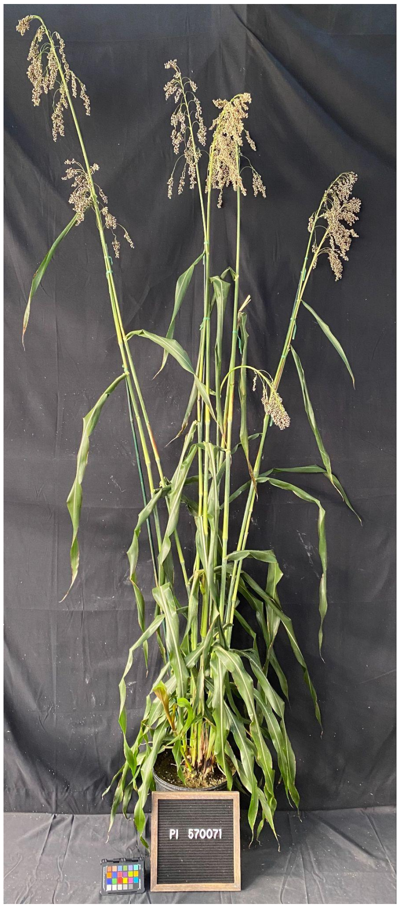


- *Stem diameter: recorded both axes means. if oval-shaped "x, y".*
- *Measurements highlighted in green and photos were recorded 18 weeks after planting (WAP).*
- *Growth conditions: 28°C (day)/22°C (night); 16h light:8h dark; Light Intensity: 400 uMol, 50% RH, Soil type: Berger BM7.*

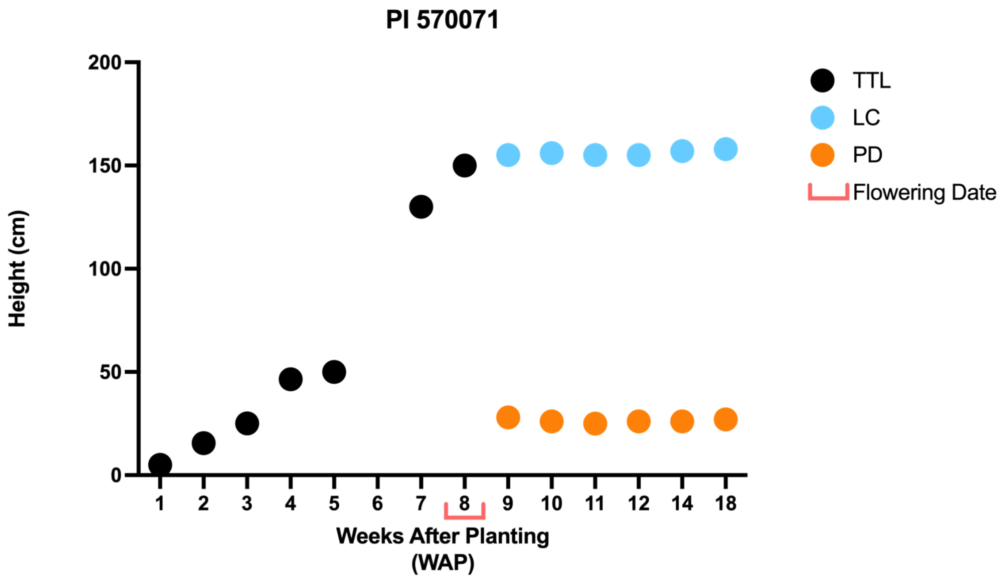


- *The plant was measured up to the last collar (LC) as well as the peduncle to the base of the panicle (PD). Before panicle formation, the height was measured up to the top of the tallest leaf (TTL).*

# PI 655988

**Other name(s):** COMBINE KAFIR-60
**Species:** Sorghum bicolor subsp. bicolor
**Group:** ICRISAT Reference Collection & Sorghum Association Panel

| | |
|---|---|
| **Flowering Days Under Short Days** | 60 days after planting |
| **Seed Color** | White |
| **Seedling Color** | Green |
| **Mid-Rib Color** | Tan |
| **Lodging** | No |
| **Stem diameter (mm) 3rd internode** | 15.7, 14.7 |
| **Stem diameter (mm) 6th internode** | 10.5 |
| **Main Stem Height: LC; PD (cm)** | 90; 10 |
| **Total Number of Tillers, with panicles** | 5, 4 |
| **Tiller 1 Stem Height: LC; PD (cm)** | 88; 19 |
| **Tiller 2 Stem Height: LC; PD (cm)** | 86; 12 |
| **Tiller 3 Stem Height: LC; PD (cm)** | 77; 19 |

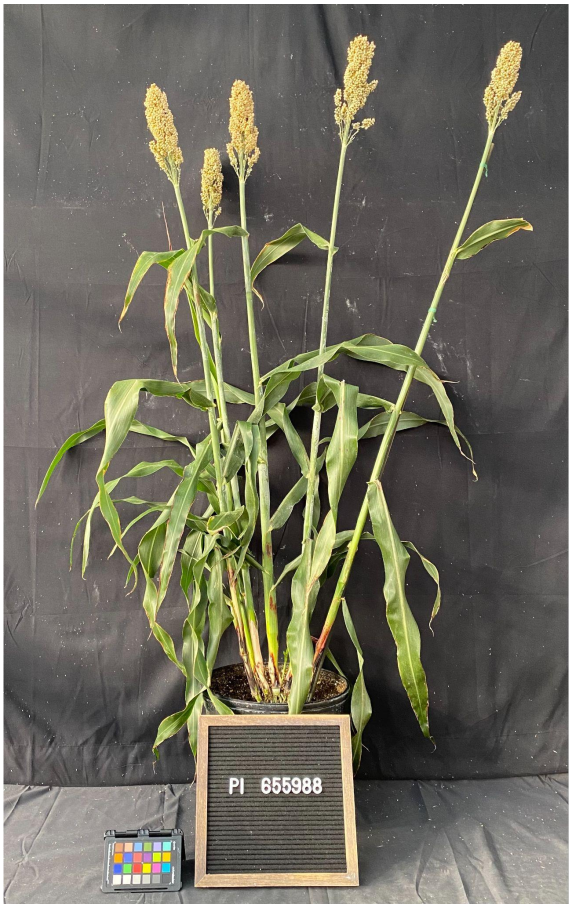


- *Stem diameter: recorded both axes means. if oval-shaped "x, y".*
- *Measurements highlighted in green and photos were recorded 18 weeks after planting (WAP).*
- *Growth conditions: 28°C (day)/22°C (night); 16h light:8h dark; Light Intensity: 400 uMol, 50% RH, Soil type: Berger BM7.*

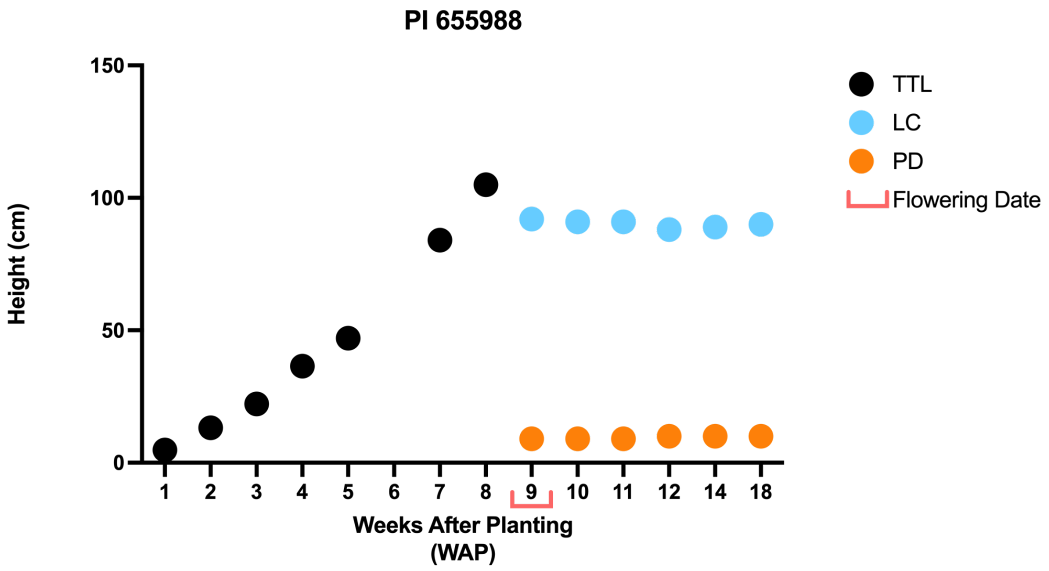


- *The plant was measured up to the last collar (LC) as well as the peduncle to the base of the panicle (PD). Before panicle formation, the height was measured up to the top of the tallest leaf (TTL).*

# PI 656044

**Other name(s):** Kuyuma
**Species:** Sorghum bicolor subsp. bicolor
**Group:** ICRISAT Reference Collection & Sorghum Association Panel

| | |
|---|---|
| **Flowering Days Under Short Days** | 66 days after planting |
| **Seed Color** | White |
| **Seedling Color** | Green |
| **Mid-Rib Color** | Tan |
| **Lodging** | No |
| **Stem diameter (mm) 3rd internode** | 20.4, 19.2 |
| **Stem diameter (mm) 6th internode** | 18.8, 16.8 |
| **Main Stem Height: LC; PD (cm)** | 96 |
| **Total Number of Tillers, with panicles** | 4, 3 |
| **Tiller 1 Stem Height: LC; PD (cm)** | 87; 5 |
| **Tiller 2 Stem Height: LC; PD (cm)** | 84 |
| **Tiller 3 Stem Height: LC; PD (cm)** | 90; 5.5 |

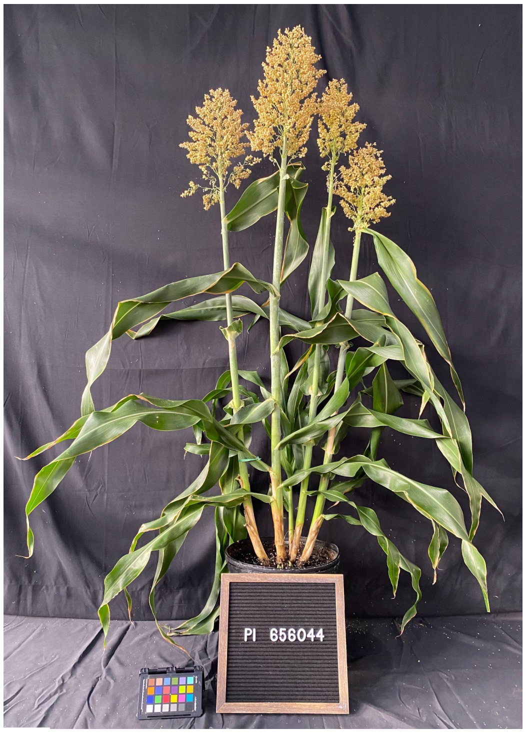


- *Stem diameter: recorded both axes means. if oval-shaped "x, y".*
- *Measurements highlighted in green and photos were recorded 18 weeks after planting (WAP).*
- *Growth conditions: 28°C (day)/22°C (night); 16h light:8h dark; Light Intensity: 400 uMol, 50% RH, Soil type: Berger BM7.*

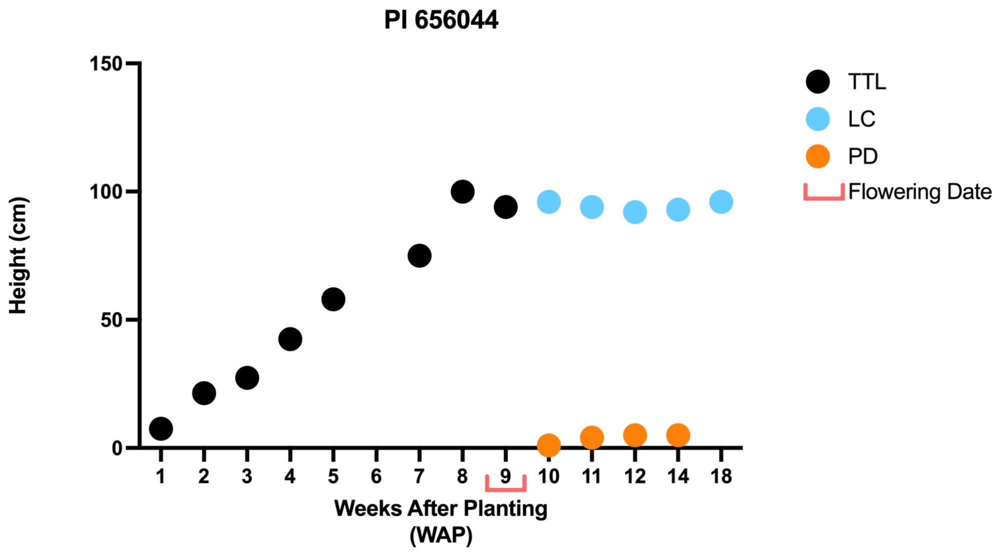


- *The plant was measured up to the last collar (LC) as well as the peduncle to the base of the panicle (PD). Before panicle formation, the height was measured up to the top of the tallest leaf (TTL)*

# PI 656029

**Other name(s):** BTx642
**Species:** Sorghum bicolor subsp. Bicolor
**Group:** ICRISAT Reference Collection & Sorghum Association Panel

| | |
|---|---|
| **Flowering Days Under Short Days** | 66 days after planting |
| **Seed Color** | Purple |
| **Seedling Color** | Green |
| **Mid-Rib Color** | White |
| **Lodging** | No |
| **Stem diameter (mm) 3rd internode** | 32.8, 23.6 |
| **Stem diameter (mm) 6th internode** | 31.5, 21.5 |
| **Main Stem Height: LC; PD (cm)** | 55; 14.5 |
| **Total Number of Tillers, with panicles** | 4, 4 |
| **Tiller 1 Stem Height: LC; PD (cm)** | 44; 28.5 |
| **Tiller 2 Stem Height: LC; PD (cm)** | 53; 31 |
| **Tiller 3 Stem Height: LC; PD (cm)** | 47; 34 |

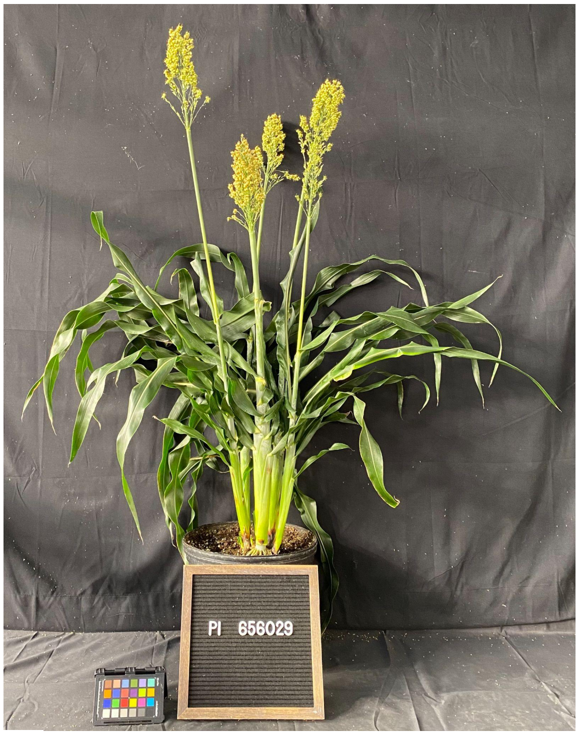


- *Stem diameter: recorded both axes means. if oval-shaped "x, y".*
- *Measurements highlighted in green and photos were recorded 18 weeks after planting (WAP).*
- *Growth conditions: 28°C (day)/22°C (night); 16h light:8h dark; Light Intensity: 400 uMol, 50% RH, Soil type: Berger BM7.*

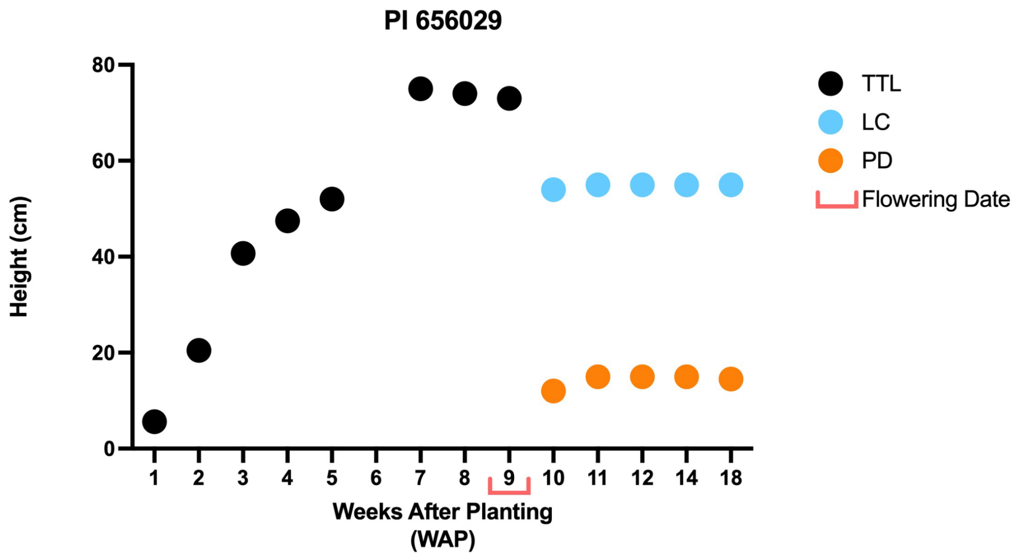


- *The plant was measured up to the last collar (LC) as well as the peduncle to the base of the panicle (PD). Before panicle formation, the height was measured up to the top of the tallest leaf (TTL).*

# PI 329301

**Other name(s):** IS 11059
**Species:** Sorghum bicolor subsp. bicolor
**Group:** ICRISAT Reference Collection, Geo-Reference Collection & Bioenergy Association Panel

| | |
|---|---|
| Flowering Days Under Short Days | 66 days after planting |
| Seed Color | Yellow |
| Seedling Color | Green |
| Mid-Rib Color | White |
| Lodging | No |
| Stem diameter (mm) 3rd internode | 18.4 |
| Stem diameter (mm) 6th internode | 15.7 |
| Main Stem Height: LC; PD (cm) | 250; 34 |
| Total Number of Tillers, with panicles | 10, 4 |
| Tiller 1 Stem Height: LC; PD (cm) | 197; 37 |
| Tiller 2 Stem Height: LC; PD (cm) | 197; 34 |
| Tiller 3 Stem Height: LC; PD (cm) | 239; 12 |

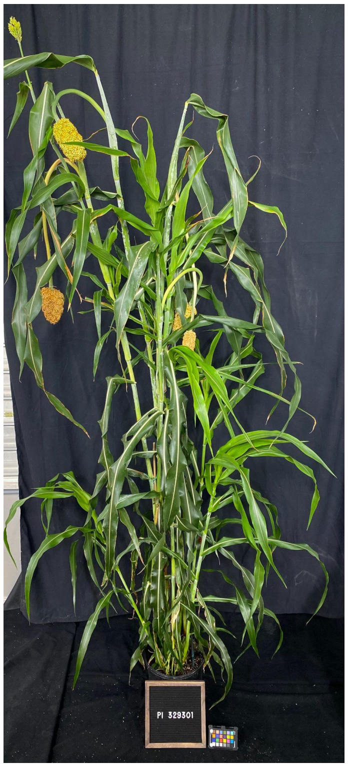


- *Stem diameter: recorded both axes means. if oval-shaped "x, y".*
- *Measurements highlighted in green and photos were recorded 18 weeks after planting (WAP).*
- *Growth conditions: 28°C (day)/22°C (night); 16h light:8h dark; Light Intensity: 400 uMol, 50% RH, Soil type: Berger BM7.*

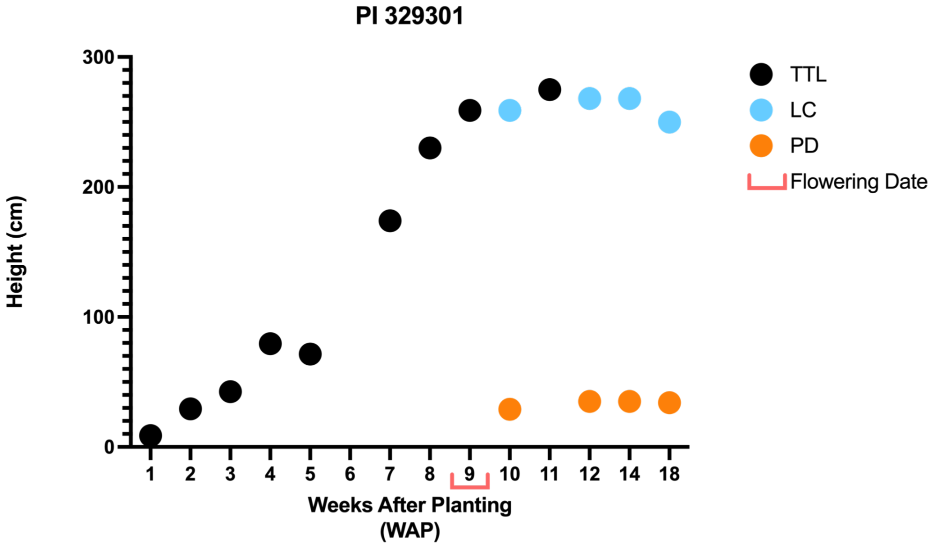


- *The plant was measured up to the last collar (LC) as well as the peduncle to the base of the panicle (PD). Before panicle formation, the height was measured up to the top of the tallest leaf (TTL).*

# PI 655996

**Other name(s):** RTx430
**Species:** Sorghum bicolor subsp. bicolor
**Group:** ICRISAT Reference Collection & Sorghum Association Panel

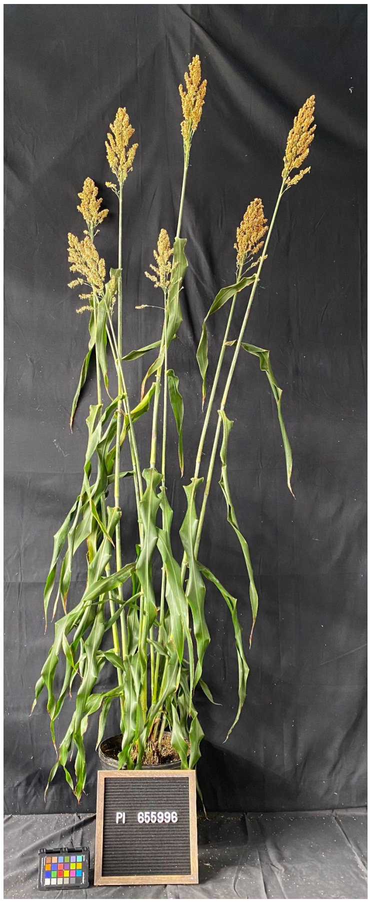


| | |
|---|---|
| **Flowering Days Under Short Days** | 60 days after planting |
| **Seed Color** | Yellow |
| **Seedling Color** | Green |
| **Mid-Rib Color** | Tan |
| **Lodging** | No |
| **Stem diameter (mm) 3rd internode** | 13.3, 12.0 |
| **Stem diameter (mm) 6th internode** | 7.4 |
| **Main Stem Height: LC; PD (cm)** | 64; 5 |
| **Total Number of Tillers, with panicles** | 15, 13 |
| **Tiller 1 Stem Height: LC; PD (cm)** | 51 |
| **Tiller 2 Stem Height: LC; PD (cm)** | 58; 14 |
| **Tiller 3 Stem Height: LC; PD (cm)** | 58; 5 |

- *Stem diameter: recorded both axes means. if oval-shaped "x, y".*
- *Measurements highlighted in green and photos were recorded 18 weeks after planting (WAP).*
- *Growth conditions: 28°C (day)/22°C (night); 16h light:8h dark; Light Intensity: 400 uMol, 50% RH, Soil type: Berger BM7.*

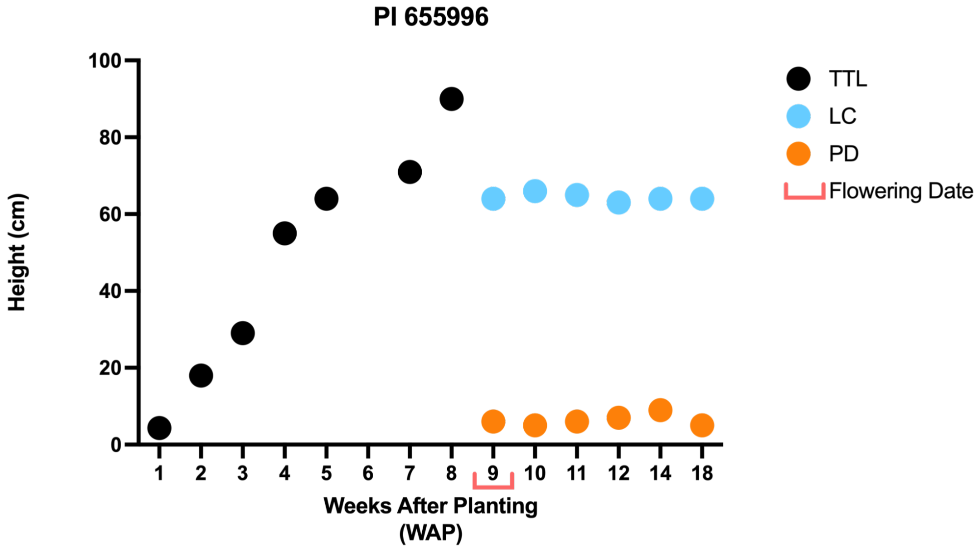


- *The plant was measured up to the last collar (LC) as well as the peduncle to the base of the panicle (PD). Before panicle formation, the height was measured up to the top of the tallest leaf (TTL).*

# PI 564163

**Other name(s):** BTx623
**Species:** Sorghum bicolor subsp. bicolor
**Group:** ICRISAT Reference Collection, Sorghum Association Panel & Bioenergy Association Panel

| | |
|---|---|
| Flowering Days Under Short Days | 75 days after planting |
| Seed Color | Purple |
| Seedling Color | Green |
| Mid-Rib Color | Tan |
| Lodging | No |
| Stem diameter (mm) 3rd internode | 14.8, 13.3 |
| Stem diameter (mm) 6th internode | 11.0, 9.2 |
| Main Stem Height: LC; PD (cm) | 87; 29 |
| Total Number of Tillers, with panicles | 10, 7 |
| Tiller 1 Stem Height: LC; PD (cm) | 98; 35.5 |
| Tiller 2 Stem Height: LC; PD (cm) | 90; 31 |
| Tiller 3 Stem Height: LC; PD (cm) | 94; 42 |

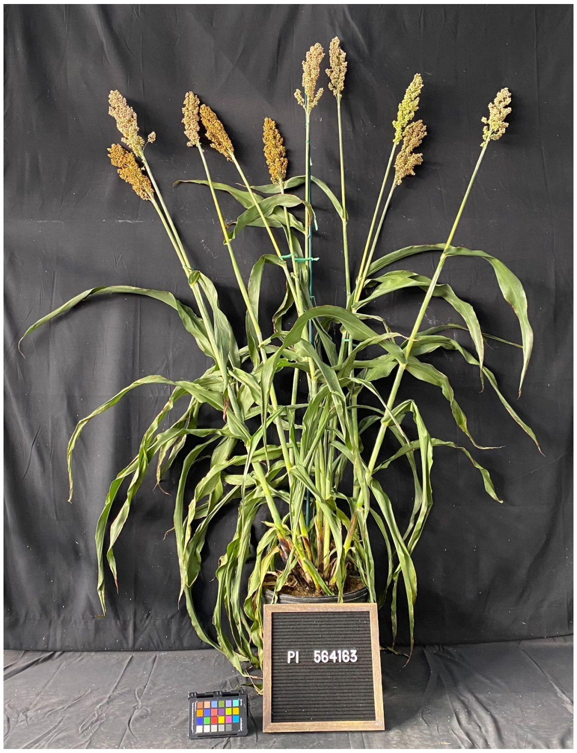


- *Stem diameter: recorded both axes means. if oval-shaped "x, y".*
- *Measurements highlighted in green and photos were recorded 18 weeks after planting (WAP).*
- *Growth conditions: 28°C (day)/22°C (night); 16h light:8h dark; Light Intensity: 400 uMol, 50% RH, Soil type: Berger BM7.*

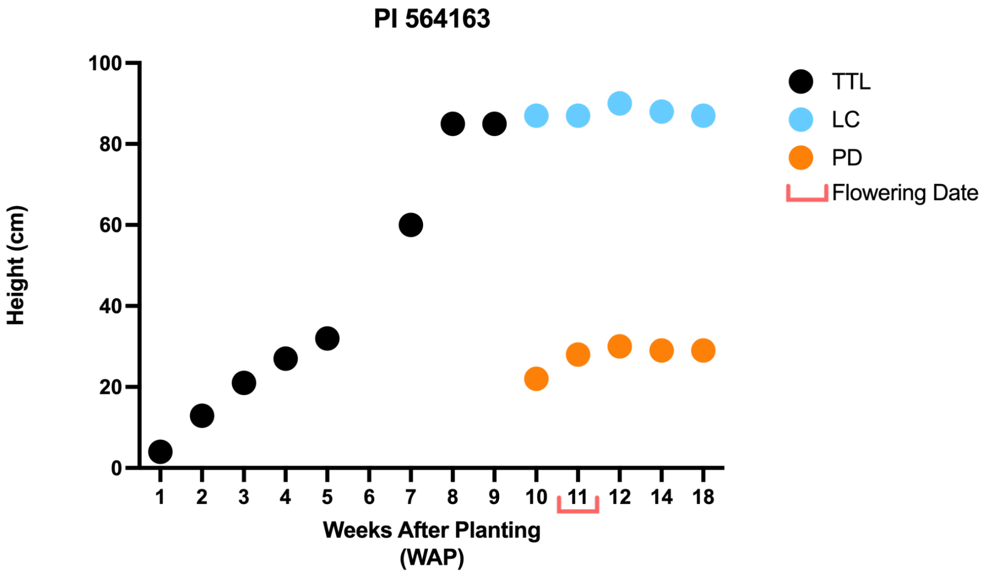


- *The plant was measured up to the last collar (LC) as well as the peduncle to the base of the panicle (PD). Before panicle formation, the height was measured up to the top of the tallest leaf (TTL).*

# PI 585966

**Other name(s):** DJOFELA, IS 26395
**Species:** Sorghum bicolor subsp. bicolor
**Group:** ICRISAT Reference Collection & Bioenergy Association Panel

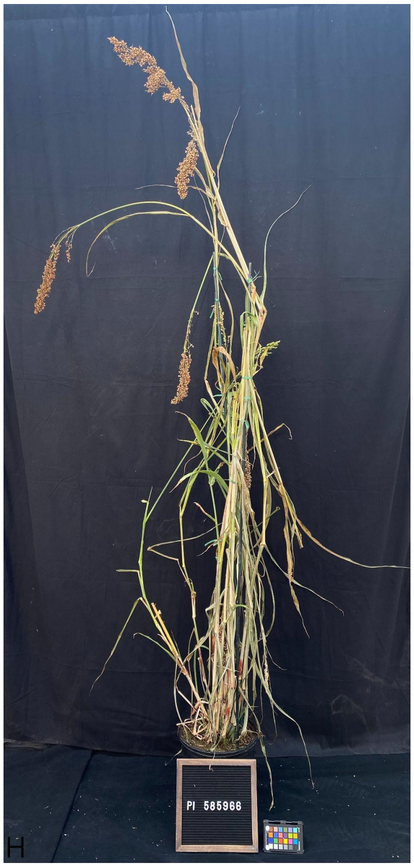


| | |
|---|---|
| Flowering Days Under Short Days | 54 days after planting |
| Seed Color | Tan |
| Seedling Color | Green |
| Mid-Rib Color | White |
| Lodging | Yes |
| Stem diameter (mm) 3rd internode | 9.8 |
| Stem diameter (mm) 6th internode | 7.2 |
| Main Stem Height: LC; PD (cm) | 184; 36.5 |
| Total Number of Tillers, with panicles | 12, 12 |
| Tiller 1 Stem Height: LC; PD (cm) | 164; 20 |
| Tiller 2 Stem Height: LC; PD (cm) | 153; 15 |
| Tiller 3 Stem Height: LC; PD (cm) | 186; 25 |

- *Stem diameter: recorded both axes means. if oval-shaped "x, y".*
- *Measurements highlighted in green and photos were recorded 18 weeks after planting (WAP).*
- *Growth conditions: 28°C (day)/22°C (night); 16h light:8h dark; Light Intensity: 400 uMol, 50% RH, Soil type: Berger BM7.*

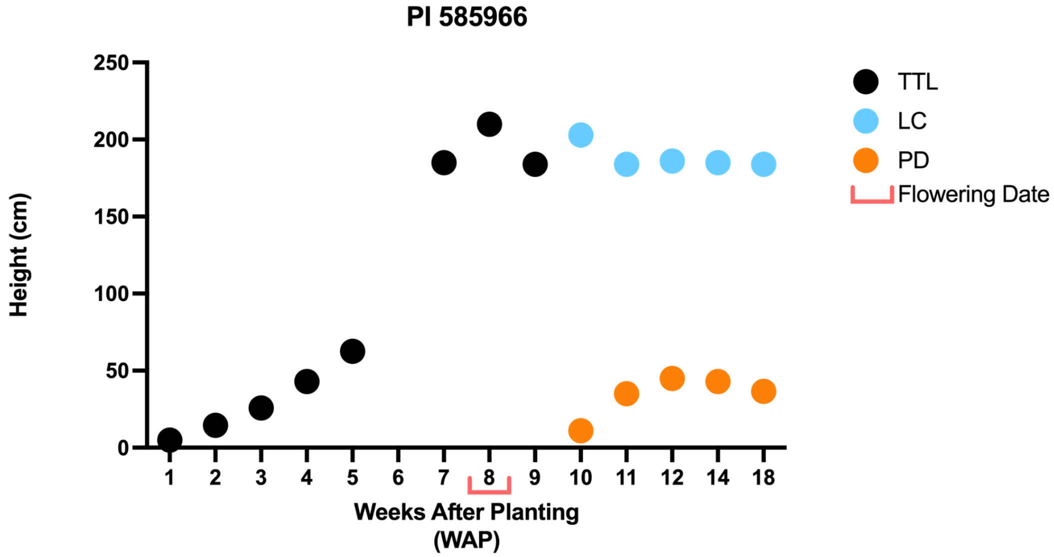


- *The plant was measured up to the last collar (LC) as well as the peduncle to the base of the panicle (PD). Before panicle formation, the height was measured up to the top of the tallest leaf (TTL).*
